# scIGANs: single-cell RNA-seq imputation using generative adversarial networks

**DOI:** 10.1101/2020.01.20.913384

**Authors:** Yungang Xu, Zhigang Zhang, Lei You, Jiajia Liu, Zhiwei Fan, Xiaobo Zhou

## Abstract

Single-cell RNA-sequencing (scRNA-seq) enables the characterization of transcriptomic profiles at the single-cell resolution with increasingly high throughput. However, it suffers from many sources of technical noises, including insufficient mRNA molecules that lead to excess false zero values, termed dropouts. Computational approaches have been proposed to recover the biologically meaningful expression by borrowing information from similar cells in the observed dataset. However, these methods suffer from oversmoothing and removal of natural cell-to-cell stochasticity in gene expression. Here, we propose the generative adversarial networks (GANs) for scRNA-seq imputation (scIGANs), which uses generated cells rather than observed cells to avoid these limitations and balances the performance between major and rare cell populations. Evaluations based on a variety of simulated and real scRNA-seq datasets show that scIGANs is effective for dropout imputation and enhances various downstream analysis. ScIGANs is robust to small datasets that have very few genes with low expression and/or cell-to-cell variance. ScIGANs works equally well on datasets from different scRNA-seq protocols and is scalable to datasets with over 100,000 cells. We demonstrated in many ways with compelling evidence that scIGANs is not only an application of GANs in omics data but also represents a competing imputation method for the scRNA-seq data.

## INTRODUCTION

Single-cell RNA-sequencing (scRNA-seq) revolutionizes the traditional profiling of gene expression, making it able to fully characterize the transcriptomes of individual cells at the unprecedented throughput. A major problem for scRNA-seq is the sparsity of the expression matrix with a tremendous number of zero values. Most of these zero or near-zero values are artificially caused by technical defects including but not limited to insufficient mRNA molecules, low capture rate and sequencing depth, or other technological factors so that the observed zero does not reflect the underlying true expression level, which is called dropout (1). A pressing need in scRNA-seq data analysis remains identifying and handling the dropout events that, otherwise, will severely hinder downstream analysis and attenuate the power of scRNA-seq on a wide range of biological and biomedical applications. Therefore, applying computational approaches to address problems of missingness and noises is very important and timely, particularly considering the increasingly popular and large amount of scRNA-seq data.

Several methods have been recently proposed and widely used to address the challenges resulted from excess zero values in scRNA-seq. MAGIC (1) imputes missing expression values by sharing information across similar cells, based on the idea of heat diffusion. ScImpute (2) learns each gene’s dropout probability in each cell and then imputes the dropout values borrowing information from other similar cells selected based on the genes unlikely affected by dropout events. SAVER (3) borrows information across genes using a Bayesian approach to estimate unobserved true expression levels of genes. DrImpute (4) impute dropouts by simply averaging the expression values of similar cells defined by clustering. VIPER (5) borrows information from a sparse set of local neighborhood cells of similar expression patterns to impute the expression measurements in the cells of interest based on nonnegative sparse regression models. Meanwhile, some other methods aim at the same goal by denoising the scRNA-seq data. DCA (6) uses a deep count autoencoder network to denoise scRNA-seq datasets by learning the count distribution, overdispersion, and sparsity of the data. ENHANCE (7) recovers denoised expression values based on principal component analysis on raw scRNA-seq data. During the preparation of this manuscript, we also noticed another imputation method DeepImpute (8), which uses a deep neural network with dropout layers and loss functions to learn patterns in the data, allowing for scRNA-seq imputation.

While existing studies have adopted varying approaches for dropout imputation and yielded promising results, they either borrow information from similar cells or aggregate (co-expressed or similar) genes of the observed data, which will lead to oversmoothing (e.g. MAGIC) and remove natural cell-to-cell stochasticity in gene expression (e.g. scImpute). Moreover, the imputation performance will be significantly reduced for rare cells, which have limited information and are common for many scRNA-seq studies. Alternatively, SCRABBLE (9) attempts to leverage bulk data as a constraint on matrix regularization to impute dropout events. However, most scRNA-seq studies often lack matched bulk RNA-seq data and thus limit its practicality. Additionally, due to the non-trivial distinction between true and false zero counts, imputation and denoising need account for both the intra-cell-type dependence and inter-cell-type specificity. Given the above concerns, a deep generative model would be a better choice to learn the true data distribution and then generate new data points with some variations, which are then independently used to impute the missing values and avoid overfitting.

Deep generative models have been widely used for missing value imputation in fields (10–12), however, other than scRNA-seq. Although a deep generative model was used for scRNA-seq analysis (13), it’s not explicitly designed for dropout imputation. Among deep generative models, generative adversarial networks (GANs) have evoked increasing interest in the computer vision community since its first introduction in 2014 (14). GANs has become an active area of research with multiple variants developed (15–19) and holds promising in data imputation (20) because of its capability of learning and mimicking any distribution of data. Given the great success of GANs in inpainting, we hypothesize that similar deep neural net architectures could be used to impute dropouts in scRNA-seq data.

In this study, we propose scIGANs, a GANs framework for scRNA-seq imputation (Figures 1A and S1). Inspired by its established applications in inpainting, we convert the expression profile of every single cell to an image, wherein the pixels are represented by the normalized gene expression. And then dropout imputation becomes the process of inpainting an image by recovering the missing pieces that represent the dropout events. Because of the inherent advantages of GANs, scIGANs does not impose an assumption of specific statistical distributions for gene expression levels and dropout probabilities. It also does not force the imputation of genes that are not affected by dropout events. Moreover, scIGANs generates a set of realistic single cells instead of directly borrowing information from observed cells to impute the dropout events, which can avoid overfitting for the cell type of big population and meanwhile promise enough imputation power for rare cells. We systematically evaluate the performance of scIGANs and compare it with 11 state-of-the-art scRNA-seq imputation methods using 3 simulated and 19 real datasets measured across different experimental protocols (Figure 1B and Table S1). We demonstrate its superior performance in recovering the biologically meaningful expression, identifying subcellular states of the same cell types, improving differential expression and temporal dynamics analysis. ScIGANs is robust to the dataset that has a small number of genes with low expression and cell-to-cell variance. Additionally, scIGANs works equally well on datasets generated by different scRNA-seq protocols and is well scalable to data size.

**Figure 1.**
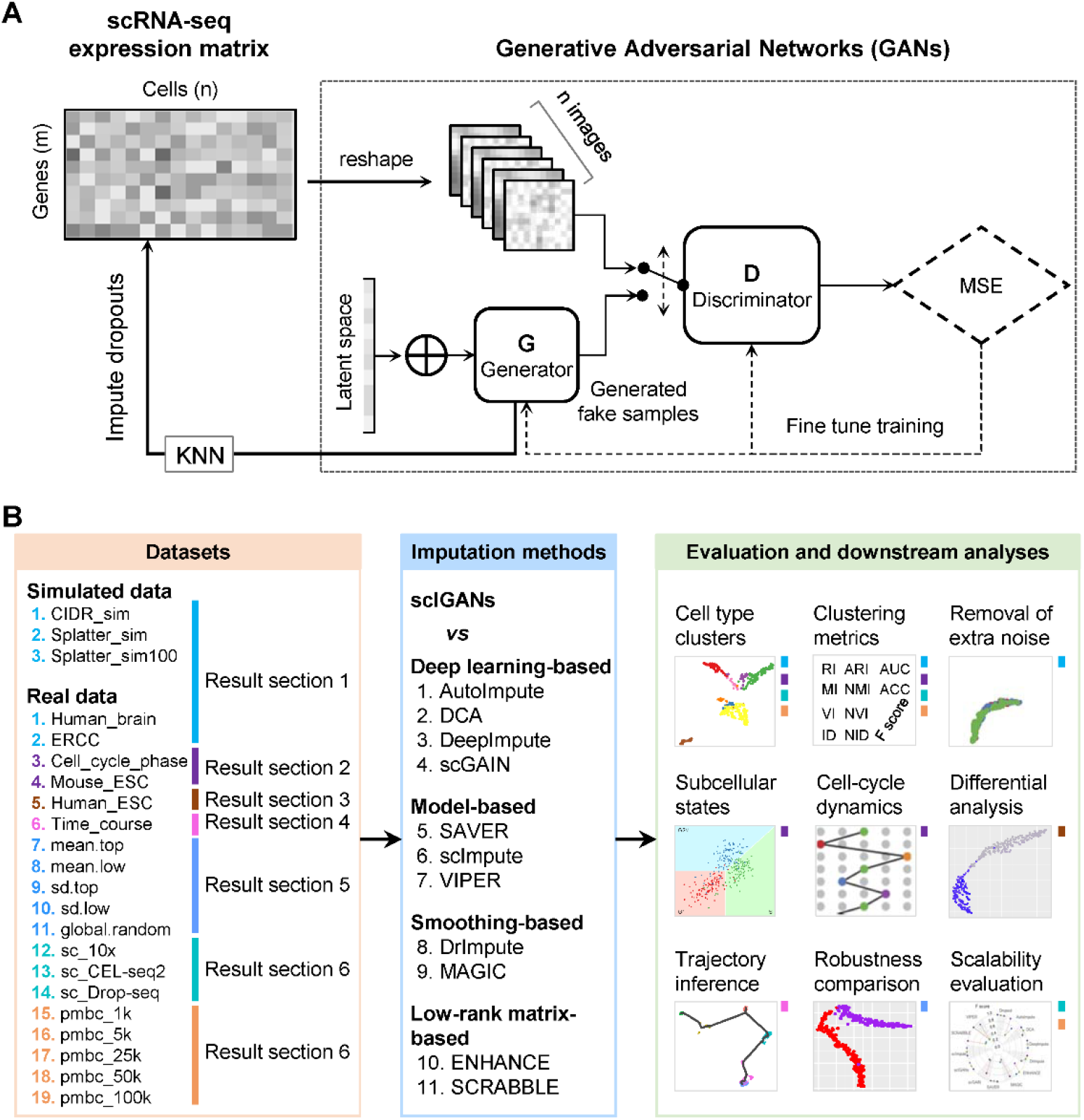
Overview of generative adversarial networks for single-cell RNA-seq imputation and systematic evaluations. **A.** The high-level architecture of scIGANs. The expression profile of each cell is reshaped to a square image, which is fed to the GANs (Supplementary Figure S1A). The trained generator is used to generate a set of realistic cells that are used to impute the raw scRNA-seq expression matrix (Supplementary Figure S1B). KNN, k-nearest neighbors. MSE, mean squared error. **B.** A schematic view of the datasets and strategies used for benchmark comparison between scIGANs and other 11 imputation methods. Colored bars in the right panel indicate the corresponding analyses and evaluations present in the result sections and datasets defined in the left panel. Also see Figures S1.

## MATERIAL AND METHODS

### The idea and design of scIGANs

Generative adversarial networks (GANs), first introduced in 2014 (14), evoked much interest in the computer vision community and has become an active area of research with multiple variants developed (15–19). Inspired by its excellent performance in generating realistic images (21–25) and recent application to generating realistic scRNA-seq data (26), we propose scIGANs, the generative adversarial networks for scRNA-seq imputation (Figure 1A). The basic idea is that scIGANs can learn the non-linear gene-gene dependencies from complex, multi-cell type samples and train a generative model to generate realistic expression profiles of defined cell types (26). To train scIGANs, the real single-cell expression profiles are first reshaped to images and fed to GANs, wherein each cell corresponds to an image with the normalized gene expression representing the pixel (Figures 1A and S1A). The generator generates fake images by transforming a 100-dimensional latent variable into single-cell gene expression profiles (Figure S1A). The discriminator evaluates whether the images are authentic or generated. These two networks are trained concurrently whilst competing against one another to improve the performance of both (Figure 1A).

Once trained, the generative model is used to generate scRNA-seq data of defined cell types. And then we propose to infer the true expression of dropouts from the generated realistic cells. The most important benefit of using generated cells instead of the real cells for scRNA-seq imputation is to avoid overfitting for the cell type of big population but insufficient power for rare cells. The generator can produce a set of cells of any number with the expression profiles faithfully characterizing the demand cell type; then the k-nearest neighbors (KNN) approach is used to impute the dropouts of the same cell type in the real scRNA-seq data (Figure S1B). The scIGANs is implemented in Python and R and compiled as a command-line tool compatible with both CPU and GPU platform. The core model is built on the PyTorch framework and adopted to accommodate scRNA-seq data as input. It’s publicly available at https://github.com/xuyungang/scIGANs.

### The strategy for scIGANs training

Training the GANs is a strategy to define a game between two competing networks. The generator network maps a source of noise to the input space. The discriminator network receives either a generated sample or a true data sample and must distinguish between the two. The generator is trained to fool the discriminator. Formally, the game between the generator *G* and discriminator *D* is the minimax objective 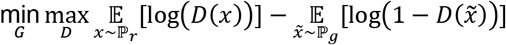; where *D* is the discriminator that can be any network, 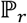 is the real data distribution and 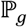 is the model distribution implicitly defined by 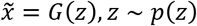; *G* is the generator which can be any network, *z* can be sampled from any noise distribution *p*, such as the uniform distribution or a spherical Gaussian distribution.

It is difficult to train the original GANs model since minimizing the objective function corresponds to minimizing the Jensen-Shannon divergence between 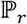 and 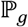 which is not continuous for the generator’s parameters. Earth-Mover (Wasserstein-1) distance *W*(*q, p*) is used to deal with such difficulty (27). Such a model is called Wasserstein GANs(WGANs) which the objective function is constructed as 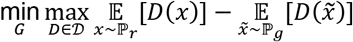; where 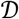 is the set of 1-Lipschitz function, the definition of other symbols are the same as the original GANs model. To enforce the Lipschitz constraint on the critic, one can clip the weights of the critic to lie within a compact space [−*c, c*]. The set of functions satisfying this constraint is a subset of the k-Lipschitz functions for some *k* which depends on *c* and the critic architecture. Researchers introduced an alternative way to enforce the Lipschitz constraint, usually called improved WGANs(IWGANs), which is widely used in training GANs models (19). The objective is 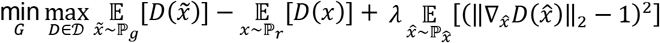; where 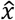 is sampled from the straight lines between pairs of points sampled from the real data distribution and the generator distribution. *λ* is a predefined parameter. BEGAN (28) is an equilibrium enforcing method paired with a loss derived from the Wasserstein distance (19) for training auto-encoder based Generative Adversarial networks. The BEGAN objective is:

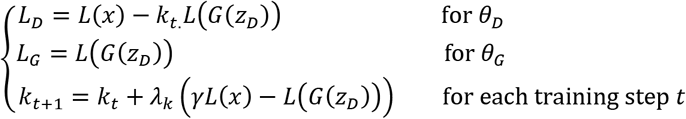

where

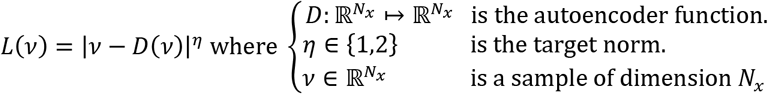

BEGAN uses Proportional Control Theory to maintain the balance between the generator and discriminator losses which is relaxed with the introduction of a new hyper-parameter *γ* ∈ [0,1] defined as 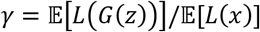. This is implemented using a variable *k_t_* ∈ [0,1] to control how much emphasis is put on *L*(*G*(*z_D_*)) during gradient descent. We initialize *k*_0_ = 0. *λ_k_* is the proportional gain for *k*; in machine learning terms, it is the learning rate for *k*. We used 0.001 as default for scIGANs and all experiments in this manuscript. In essence, this can be thought of as a form of closed-loop feedback control in which *k_t_* is adjusted at each step to maintain equation 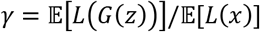. In this work, we use this method to train our scIGANs.

### The procedures for scIGANs training and dropout imputation

To be noted that scIGANs is designed scalable to the datasets with any number of cell types and genes, though we here assume an example dataset with 9 cell types and 32*32=1024 genes to elucidate how it works. The generator network of scIGANs is defined as *G*(*z, La_z_*; *θ*). The inputs of the generator are: *z* ∼ norm(0,1), and label *La_z_* ∼ *U*(1,9). Denote *θ* as the parameters need to be learned. The generator is defined as following the steps (Supplementary Figure S1A):

1. Do transposed convolution on *z* by *GConv1_1* and get the *tensor z_n_* of dimension (32,32,32).
2. Do transposed convolution on *La_z_* by *GConv1_2* and get the *tensor La_n_* of dimension (8,32,32).
3. Concatenate *z_n_* and *La_n_* to get *GConcat1* of dimension (40,32,32).
4. Do convolution on *GConcat1* by *GConv2_1* and *GConv2_2* to get the tensor of dimension (1,32,32), which is the output of the Generator.

The discriminator network is defined as *D*(*x,La_x_;w*). The inputs of discriminator are samples of real data 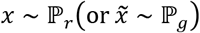 representing the reshaped expression profile of an individual cell, and label of *x*(or 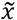) denoted by *La_x_* representing the cell type or subpopulation. Denote *w* as the parameters need to be learned. The discriminator is defined as following the steps (supplementary Figure S1A):

1. Do convolution on *x* or 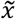 by *DConv1_1* and get the *tensor* of dimension (16,32,32).
2. Do convolution on *Lα_x_* by *DConv1_2* and get the tensor of dimension (16,32,32).
3. Concatenate results of steps (1) and (2) as *Dconcat1*, which is a tensor of the dimension (32,32,32).
4. Convert the *Dconcat1* to a vector of length 16 using a fully connected network (FCN).
5. Do convolution on the result of step (4) by *GConv2_1* and *GConv2_2* to get the tensor of dimension (1,32,32), which is the output of the Discriminator.

With a well-trained GANs model, for a given cell *c_i_* which belongs to the subpopulation *K_c_i__*, we generate a candidate set *A^K_c_i__^* with *n_can_* expression profiles. Denote *c′_i_knn__* as the *k* nearest neighbors using Euclidian distance in the set *A^K_c_i__^*. We then use the following equation to impute *j*th gene in the cell *c_i_* (Supplementary Figure S1B): 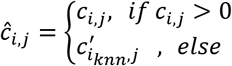.

### Data processing and normalization

The data of a scRNA-seq study are usually organized as a read count matrix with *N* rows representing genes and *M* columns representing cells, which is the input of scIGANs. Since scIGANs is trained similarly to the training for image processing, we need to transfer the expression profile of each cell to a grayscale image (Supplementary Figure S1A). To this end, scIGANs firstly normalizes the raw count matrix by the maximum read count of each sample (cell) so that all genes of each sample will have the expression values in a [0,1] range. scIGANs then reshapes the expression profile of each cell to a square image in a column-wise manner, with the normalized gene expression values representing the pixels of the image. The image size will be *n* × *n*, where *n* is the minimum integer so that *n* × *n* ≥ N. If the gene number is less than *n* × *n*, extra zeroes will be filled. Then, a scRNA-seq matrix with *M* cells will be represented as *M* grayscale images and used to train a conditional GANs with the cell labels.

### Simulated scRNA-seq datasets

We first simulated a simple scRNA-seq data with 150 cells and 20180 genes using the default CIDR simulation function *scSimulator(N=3, k=50)* (29). Three cell types are generated with 50 cells for each [dataset in Table S1: CIDR_sim]. The raw data has a dropout rate of 52.8%. Figures 2A, S2A, and Table S2 are derived from this data. We then tested the performance of different imputation methods on different dropout rates simulated by Splatter (30). We took the same simulation strategy used by SCRABBLE (9) with the same parameters for the Splatter simulator. Specifically, three scRNA-seq datasets with three different dropout rates (71%, 83%, and 87%) were simulated; each dataset has 800 genes and 1000 cells grouped into three clusters (cell types) [dataset in Table S1: Splatter_sim]. Figures S2B-E and Table S3 were derived from these datasets. To test the robustness of imputation methods, we repeated 100 times of the above Splatter simulations and generated 100 datasets for each of the above three different dropout rates [dataset in Table S1: Splatter_sim100]. Figures 2B, S3A-F, and Table S4 (EXCEL) were derived from these datasets.

**Figure 2.**
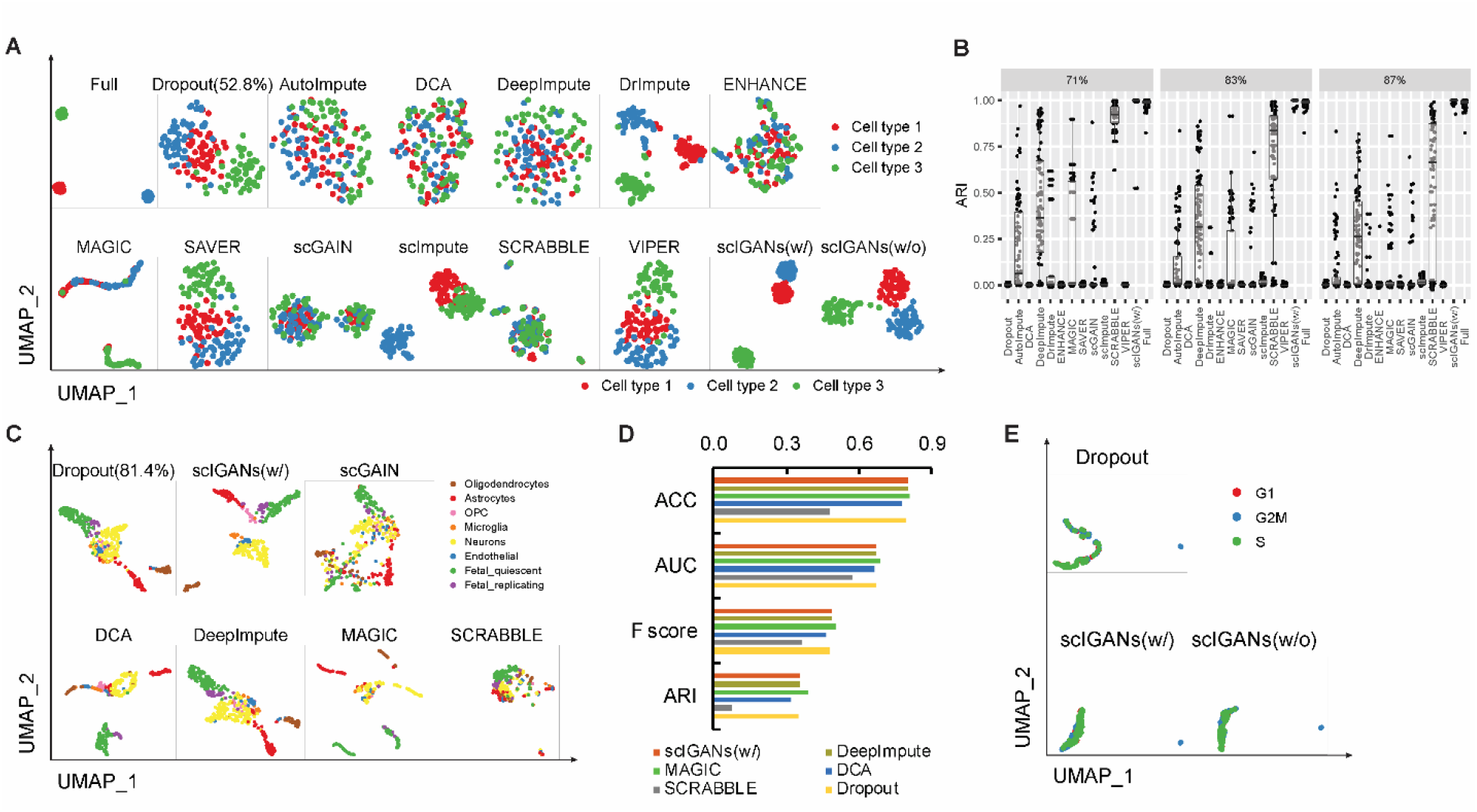
ScIGANs recovers single-cell gene expression from dropouts without extra noise. **A.** The UMAP plots of the CIDR simulated scRNA-seq data for Full, Dropout, and imputed matrix by 12 methods. Multiple clustering measurements are provided in Supplementary Figure S2A and Table S2. **B.** The adjusted rand index (ARI), a representative clustering measurement to indicate performance and robustness of all methods on the Splatter simulated data with three different dropout rates (71%, 83%, and 87%) and 100 replicates for each. Full list of clustering measurements provided in Supplementary Table S4. **C.** The selected UMAP plots of real scRNA-seq data for the human brain. **D.** The selected clustering measurements for the scRNA-seq data of the human brain. ACC, accuracy; AUC, area Under the Curve of ROC (receiver operating characteristics); F score, the harmonic mean of precision and recall; NMI, normalized mutual information. The full list of all considered clustering measurements is provided in Supplementary Table S5. **E.** The evaluation of robustness in avoiding extra noise using scRNA-seq data of spike-in RNAs. All UMAP plots are provided in Supplementary Figure S3H. Also see Figures S2-S3.

### Real scRNA-seq datasets

#### Human brain scRNA-seq data

We used scRNA-seq data of 466 cells capturing the cellular complexity of the adult and fetal human brain at a whole transcriptome level [dataset in Table S1: Human_brain] (31). Tag tables were downloaded from the data repository NCBI Gene Expression Omnibus (GEO access number: GSE67835) and combined into one table with columns representing cells and rows representing genes. We excluded the uncertain hybrid cells and remained 420 cells in eight cell types with the expression of 22085 genes. This dataset was used to generate Figures 2C-D, S3G, and Table S5.

#### ERCC spike-in RNAs scRNA-seq data

In a scRNA-seq dataset for mESCs (32) [dataset in Table S1: ERCC], ERCC spike-in RNAs were added to each cell and sequenced. ERCC spike-in RNAs consist of 92 RNA transcripts in the length of 250 to 2,000 nt, which are widely used in scRNA-seq experiments to remove the confounding noises from biological variability. Since spike-in RNAs are added to samples with the identical amount to capture the technical noise, the readout for the spike-in RNAs should be free of cell-to-cell variability and the detected variance of expression, if exists, should only come from technical confounders other than biological contexts (e.g. cell types). Therefore, the expression profiles of spike-in RNAs that were added to individual cells should not be able to cluster these cells into different subgroups regarding cell types or other biological states. Therefore, we used the ERCC spike-in read counts from the real scRNA-seq data for mESCs to evaluate the imputation methods on denoising the technical variation without introducing extra noise. This data was used to generate Figures 2E and S3H-I, and Table S6.

#### Cell-cycle phase scRNA-seq data

To evaluate the performance of different imputation methods on identifying different cellular states of the same cell type, we analyzed a single-cell RNA-seq data from mESCs (32) [dataset in Table S1: Cell_cycle_phase]. A set of 96 asynchronously dividing cells for each cell-cycle phase of G1, S, and G2M was captured using the Fluidigm C1 system, and sequencing libraries were prepared and processed. In this dataset, 288 mESCs were profiled and characterized by 38293 transcripts with a dropout rate of 74.4%. This dataset was used to generate Figures 3A-B and S4, and Table S7. Specifically, the cell states of individual cells in Figures 3B and S4B were inferred by R package Seurat (v3.1) based on a collection of predefined cell cycle marker genes (33,34).

**Figure 3.**
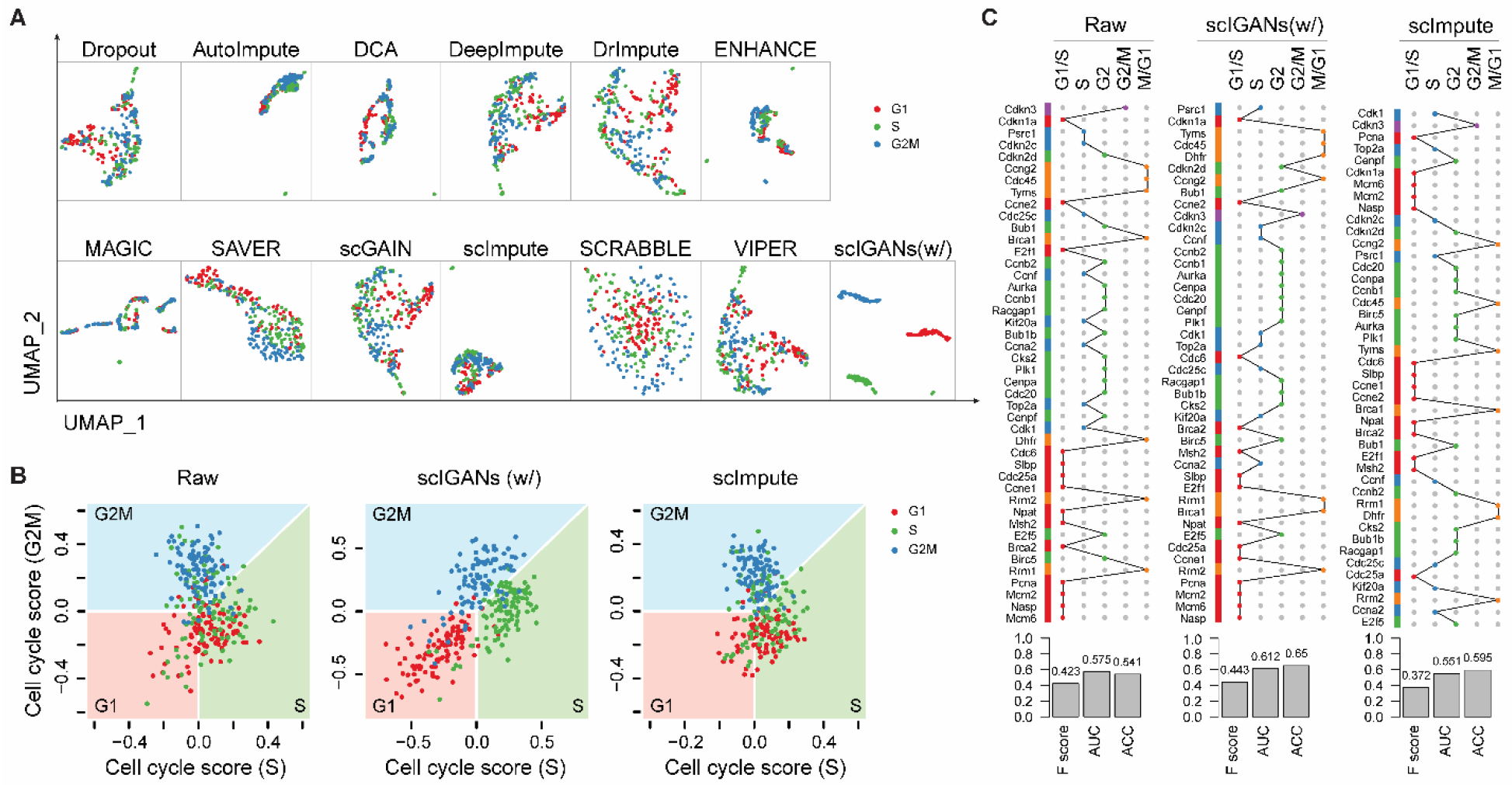
ScIGANs enables the identification of cell-cycle states and dynamics. **A.** The UMAP plots of the real scRNA-seq data for cell-cycle states of homogeneous mouse ESCs. The full list of all clustering metrics is provided in Supplementary Figure S4A and Table S7. **B.** Cells are projected to the cell-cycle phase spaces based on a collection of cell-cycle marker genes. The plots for all other methods are provided in Supplementary Figure S4B. **C.** Cell cycle dynamics are shown as the hierarchical clustering of 44 cell-cycle-regulated genes across 6.8k mouse ESCs. Full dynamic cell-cycle profiles from the scRNA-seq data before and after imputation by different methods are provided in Supplementary Figure S4C-N. The barplots show the quantitative concordance between the assigned cell-cycle phases by hierarchical clustering and the true phases that these genes serve as the markers. F score, the harmonic mean of precision and recall; AUC, area Under the Curve of ROC (receiver operating characteristics); ACC, accuracy. Also see Figure S4.

#### Mouse ESC scRNA-seq dataset for cell-cycle dynamics

Mouse embryonic stem cells (mESC) were profiled using the droplet-microfluidic scRNA-seq approach with 1 biological replicate (933 cells) and 2 technical replicates (2509 and 3443 cells for each) [dataset in Table S1: Mouse_ESC]. The processed count matrix was downloaded from Gene Expression Omnibus (GEO) with the access ID GSE65525. All other 11 imputation methods and scIGANs were used to impute the raw matrix with an exception that SCRABBLE and DrImpute failed to impute this data because take longer than a month to finish the imputation. This data was used to generate Figures 3C and S4C-N.

Cell cycle dynamics assessment was performed according to Figure 6E-F of reference (35). Briefly, the Pearson’s correlation was applied among a list of previously categorized 44 cell-cycle genes based on their expression across these ∼6.8k cells. Genes were ordered by hierarchical clustering on the correlation matrix and their previously categorized cell-cycle phases were indicated as linked dots representing cell-cycle oscillations (Figures 3C and S4C-N). Clustering measurements were also applied to the gene clusters against their pre-assigned cell-cycle phased (barplots in Figures 3C and S4C-N), which represent the performances of imputation methods by clustering the cell-cycle genes across cells.

#### Human ESC scRNA-seq dataset for differential expression analysis

To compare the performance of different imputation methods on detecting differentially expressed genes (DEGs), we utilized a dataset with both bulk and single-cell RNA-seq experiments on human embryonic stem cells (ESC) and the differentiated definitive endoderm cells (DEC) (36) [dataset in Table S1: Human_ESCs]. This dataset includes six samples of bulk RNA-seq (four for H1 ESC and two for DEC) and scRNA-seq of 350 single cells (212 cells for H1 ESC and 138 cells for DEC). The percentage of zero expression is 14.8% for the bulk dataset and 49.1% for the single-cell dataset. This dataset was used to generate Figures 4 and S5-S7.

**Figure 4.**
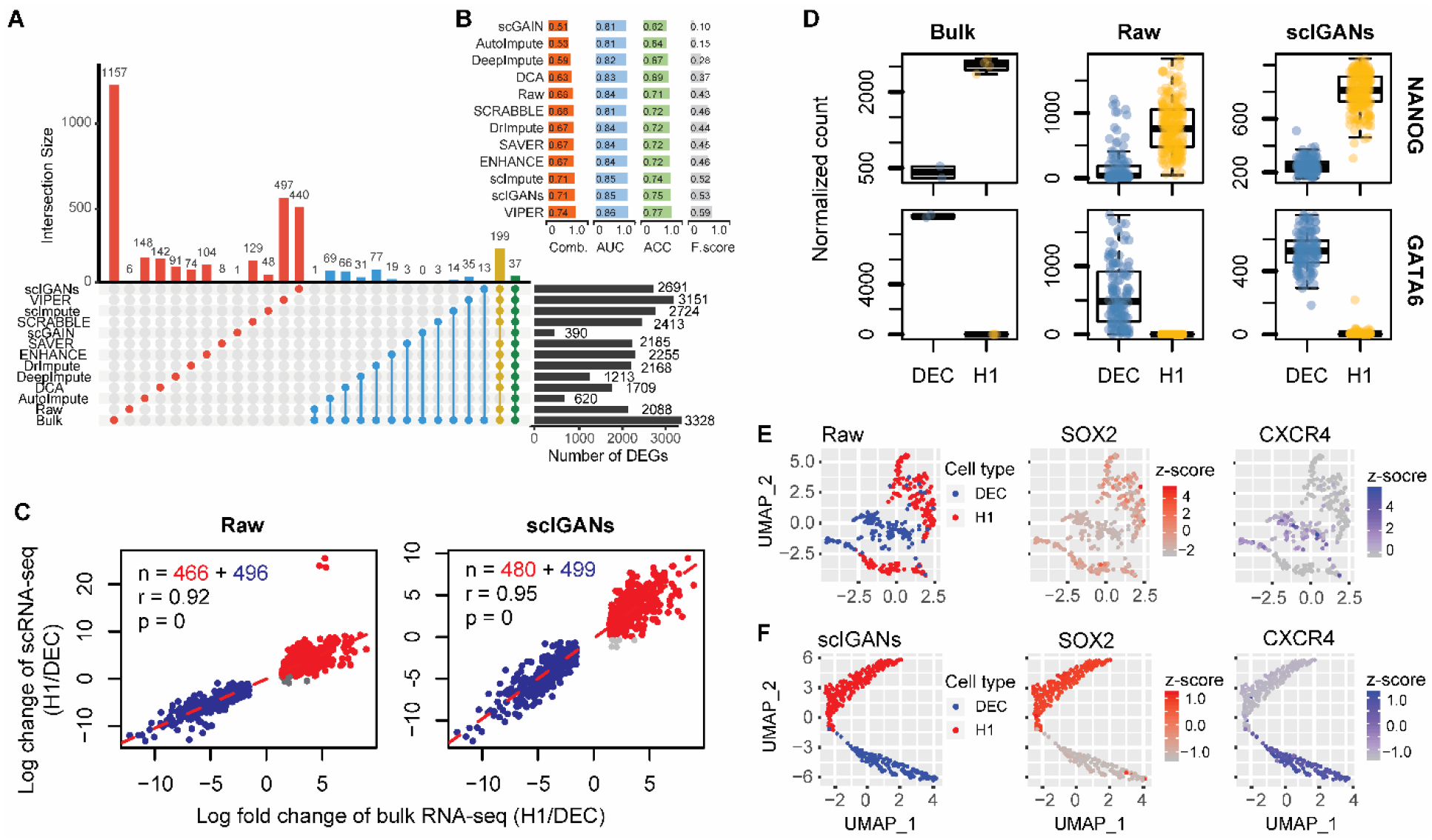
ScIGANs increases the correspondence between single-cell and bulk differential expression analysis. **A.** The correspondence of differentially expressed genes (DEGs) between bulk and single-cell RNA-seq with different imputation approaches. The yellow and green bars (connoted dots) highlight the poorest agreement of AutoImpute and scGAIN with all other methods. **B.** The barplots show the performances of DEG detections from raw and imputed scRNA-seq datasets based on the gold standard defined by the bulk RNA-seq dataset. F score, the harmonic mean of precision and recall; AUC, area Under the Curve of ROC (receiver operating characteristics); ACC, accuracy; Comb., the combined overall performance as the average of the AUC, ACC, and F score. **C.** The correlations between log fold-changes of differentially expressed genes from bulk and single-cell RNA-seq datasets. Detailed legends and the plots of results from all other imputation methods are provided in Supplementary Figure S5. **D.** The expression for one of five selected signature genes of H1 and DEC cells, respectively. All plots of other genes with different imputation methods are provided in Supplementary Figure S6. **E-F.** The UMAP plots of the single cells overlaid by the expression of SOX2 and CECR4, which is the marker gene of H1 and DEC cells, respectively. Raw (E) and scIGANs imputed (F) data are shown and data from all other methods are provided in Supplementary Figure S7. Also see Figure S5-S7.

We used scIGANs and 11 other imputation methods to impute the gene expression for single cells and then used DESeq2 (37) to perform differential expression analysis on the raw and 11 imputed data, respectively. DEGs are genes with the absolute log fold changes (H1/DEC) ≥ 1.5, adjust-p ≤ 0.05, and base mean ≥ 10. Taking the DEGs from bulk RNA-seq data as the gold standard, the overall performances of DEG detection from imputed scRNA-seq datasets were defined as the correspondences of DEGs between bulk RNA-seq and scRNA-seq (Figure 4A). To more fairly quantitate the performance of DEG detection using scRNA-seq data, we took the DEG detection as the process of predicting a gene is DEG or not, based on the gold standard from bulk RNA-seq. Then we calculated the accuracy (ACC), F score (also F1-score or F-measure), and the area under the receiver operating characteristic curve (AUC) for each DEG detection from the imputed scRNA-seq datasets. The overall performance was defined as the average of the above three measurements (Figure 4B). Additionally, a set of top 1000 DEGs (500 best up-regulated and 500 best down-regulated genes based on the adjust-p values) from bulk RNA-seq data were used to calculate the correlations between log fold changes of DEGs from bulk and single-cell RNA-seq datasets. between scRNA-seq and bulk RNA-seq data (Figures 4C and S5). To further illustrate the improvement of imputation on DEG detection, five signature genes highlighted in Figure 1c of the source paper (37) for H1 and DEC, respectively, were plotted out (Figures 4D and S6). The expression of two marker genes (SOX2 for H1 cell and CXCR4 for DEC cell) were overlaid to the UMAP space of single cells to show the expression signature of these two cell types (Figures 4E-F and S7).

#### Time-course scRNA-seq data for cellular trajectory analysis

We utilize a time-course scRNA-seq data derived from the differentiation from H1 ESC to definitive endoderm cells (DEC) (36) [dataset in Table S1: Time_cousre]. A total of 758 cells were profiled at 0 (cell number n=92), 12 (n=102), 24 (n=66), 36 (n=172), 72 (n=138), and 96 (n=188) hours after inducing the differentiation from H1 ESCs to DECs (Figure 5A). We apply scIGANs and all other 11 imputation methods to the raw scRNA-seq data with known time points and then reconstruct the trajectories using Monocle3 (38). This dataset was used to generate Figures 5 and S8.

**Figure 5.**
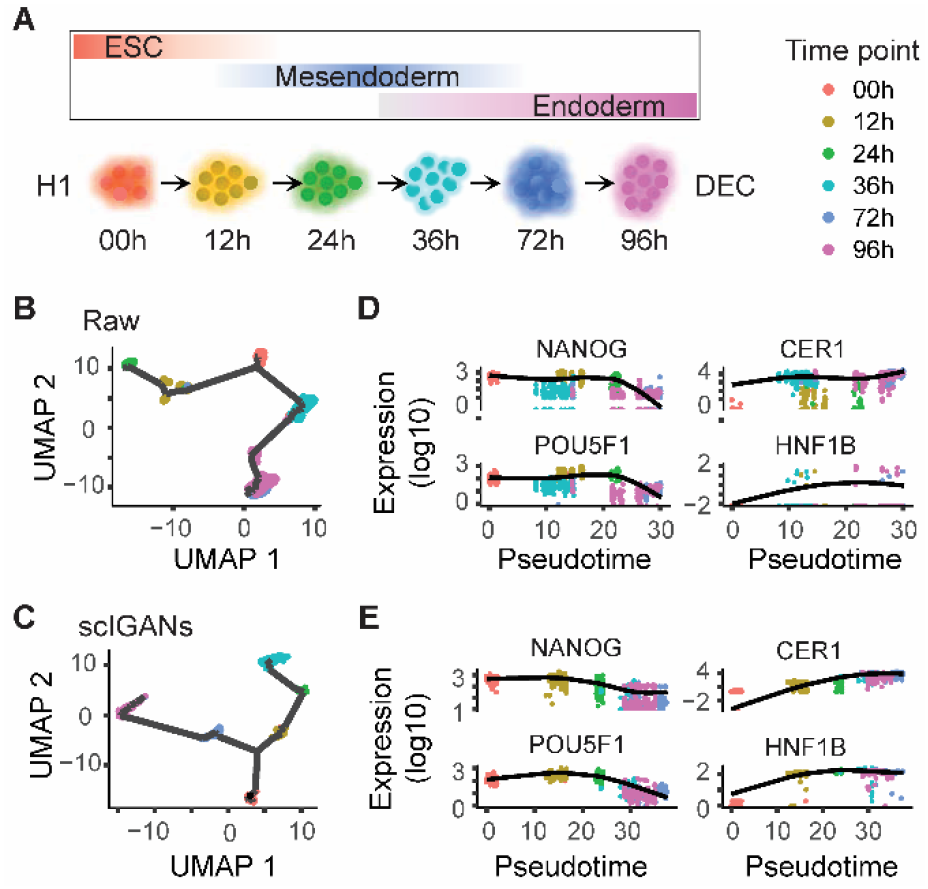
scIGANs improves time-course scRNA-seq data analysis and reconstructing differentiation trajectory. **A.** The time points of scRNA-seq sampling along with the differentiation from the pluripotent state (H1 cells) through mesendoderm to definitive endoderm cells (DEC). **B-C.** The trajectories reconstructed by monocle3 from the raw (C) and scIGANs imputed (D) scRNA-seq data. **D-E.** The expression dynamics of two pluripotent (left) and DEC (right) signature genes are shown in the order of the pseudotime. The plots of all other imputation methods are provided in Supplementary Figure S8. Also see Figure S8.

### Subsampled scRNA-seq datasets

We subsampled the scRNA-seq data derived from human embryonic stem cells (ESC) and the differentiated definitive endoderm cells (DEC) (36). This dataset has expression profiles of 350 single cells (212 for H1 ESC and 138 for DEC) across 19097 genes. Three different sampling strategies were used to generate different sub-datasets for robustness tests. These datasets were used to generate Figures 6 and S9.

**Figure 6.**
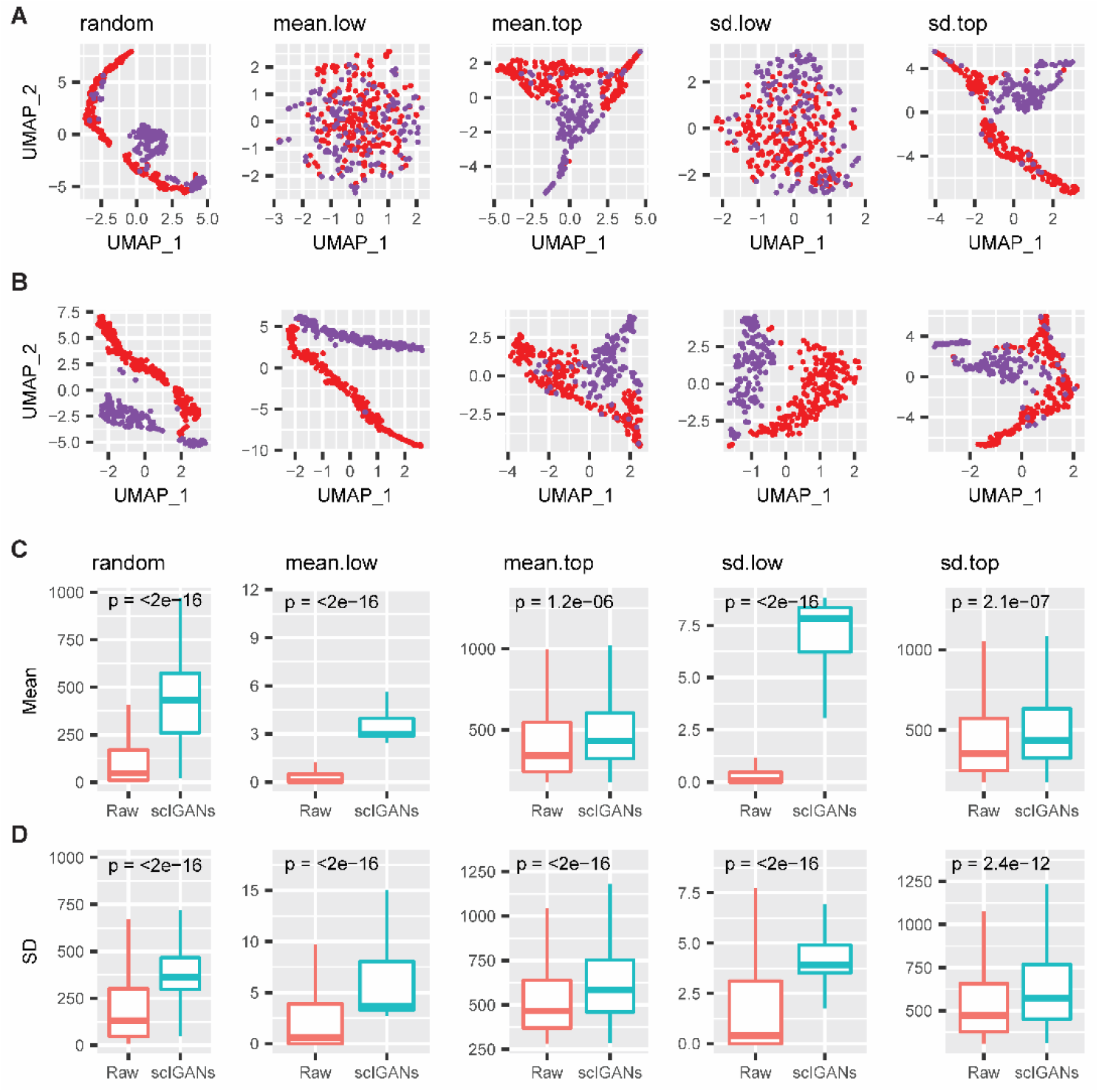
scIGANs is robust to a small set of genes with very low expression or cross-cell variance. **A-B.** The UMAP visualizations of H1 and DEC cells using only 1024 genes from raw (A) or scIGANs imputed (B) expression matrix based on three different sampling strategies. The sampling strategies are described in Methods. **C-D.** The boxplots show the mean (C) or standard deviation (sd, D) of the 1024 sampled genes before and after scIGANs imputation. The same series of plots for all other imputation methods are provided in Supplementary Figure S9. Also see Figure S9.

1) datasets with a subset of genes that have top- and lower-mean of expressions across all 350 cells, denoted as mean.top and mean.low (Table S1). Specifically, the expression matrix (genes in rows and cells in columns) was sorted by the row-wise means (descending) and the first and last 5000 genes were selected, representing two subsets with high and low expressions, respectively. Then 1024 (32*32) genes were randomly picked from these 5000 genes to generate the two test datasets, mean.top and mean.low with the zero-rate of 6.34% and 97.25%, respectively.

2) datasets with a subset of genes that have top- and lower-standard deviation (sd) of expressions across all 350 cells, denoted as sd.top and sd.low (Table S1). Specifically, the expression matrix (genes in rows and cells in columns) was sorted by the row-wise standard deviation (descending) and the first and last 5000 genes were selected, representing two subsets with high and low expression standard deviations, respectively. Then 1024 (32*32) genes were randomly picked from these 5000 genes to generate the two test datasets, sd.top and sd.low with the zero-rate of 8.72% and 92.42%, respectively.

3) dataset with a subset of 1024 genes randomly selected from all 19097 genes, denoted as global.random (Table S1). It has the zero-rate of 49.51%.

### Cross-platform scRNA-seq datasets

To evaluate how scIGANs works equally well for different scRNA-seq protocols/platforms, we collected a cross-platform benchmark dataset derived from 3 human lung adenocarcinoma cell lines, including H1975, H2228, and HCC827 (39). The three cell lines were mixed equally and processed by 10X Genomics/Chromium, CEL-seq2/Fluidigm, and Drop-seq/Droplet, by which the datasets were generated and referred to as sc_10X, sc_CEL-seq2 and sc_Drop-seq, respectively (Table S1). We clustered the cells and calculate the clustering metrics based on the raw matrix and matrix imputed by different methods (Figures 7A-B, S10-S11, and Table S8). We also calculated the Spearman correlation coefficients between the expression profiles of the same cell types from different sequencing methods (Figures 7C and S12A). Taking H1975 cell between sc_10X and sc_CEL-seq2 as an example, the Spearman correlation coefficients were calculated for each cell-cell pair, of which one H1975 cell from the sc_10X dataset and the other H1975 cell from the sc_CEL-seq2 dataset. Figures 7C and S12A are plotted using a random subsample of 500 cell-cell pairs for each comparison.

**Figure 7.**
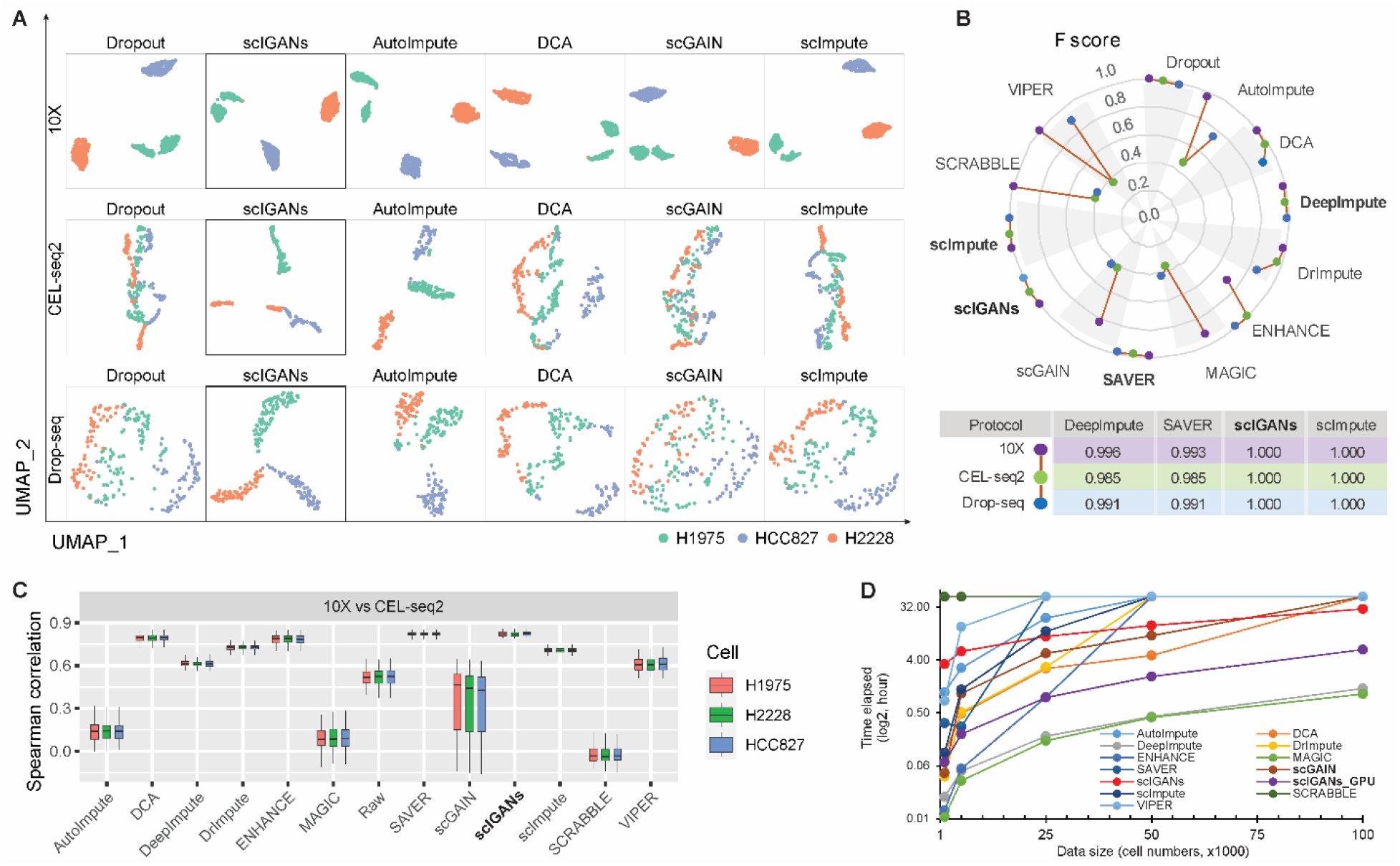
scIGANs is scalable to scRNA-seq methods and data sizes. **A.** The UMAP visualizations of the same cell populations sequenced by three different scRNA-seq methods (protocols/platforms). 10X, 10X Genomics/Chromium; CEL-seq2, CEL-seq2/Fluidigm; Drop-seq, Drop-seq/Droplet. **B.** Cross-platform performances of different methods represented as the F scores for cell type clusters against the pre-defined cell types. The same series of plots and source data for all other clustering metrics are provided in Supplementary Figure S11 and Table S8. **C.** Cell-cell correspondence of the expression profiles between 10X and CEL-seq2, shown as the Spearman correlation coefficients from the expression profiles of the same cell types before and after imputation by different imputation methods. Correlations between other sequencing methods are provided in Supplementary Figure S12A. **D.** The running time (in hour) of the imputation methods on datasets with different sizes (cell numbers). The failed jobs in Supplementary Table S9 are assigned auxiliary values of 49 hours for plotting purposes. Also see Figure S10 - S12.

### scRNA-seq datasets with different cell numbers

To test the scalability of scIGANs and 11 other imputation methods to data sizes, we created a set of scRNA-seq datasets by randomly sampling 1000 (pmbc_1k), 5000 (pmbc_5k), 25000 (pmbc_25k), 50000 (pmbc_50k), and 100000 (pmbc_100k) cells from the PMBC 10k dataset (Table S1, https://support.10xgenomics.com/single-cell-gene-expression/datasets/3.0.0/pbmc_10k_v3). These datasets were used to test the running time and memory usage of each method on different data sizes (cell numbers). All methods were tested on a single node of the TACC Lonestar 5 (LS5) system (https://portal.tacc.utexas.edu/user-guides/lonestar5). Specifically, we assessed the scalability to the data sizes of each method by computational time (in hours) and memory usage (in gigabytes, GB) using the values “Elapsed” and “AveRSS” from the job information returned from Slurm command *sacct*. To be noted that, some methods were failed on some datasets because they need either more than 48 hours (Failed_T) or more than 64 GB of the memory (Failed_M), which exceeds the limit of the TACC Lonestar system (Table S9). Correspondingly, these failed jobs were assigned 49 hours for running time and 65 GB for memory usage, respectively, so that all methods can be plotted at all data points in Figures 7D and S12B. scIGANs was also tested on an in-house GPU system configured with one NVIDIA Tesla V100 32GB Graphic Card, from which the measurements are labeled as scIGANs_GPU in Figures 7D, S12B and Table S9.

### Implementation

ScIGANs is implemented in Python (>2.7) and R (>3.6) with a Linux/Unix wrapper script. An expression matrix of the single cells is the only required input file. Optionally, a file with the cell labels (cell type or subpopulation information) can be provided to direct scIGANs for cell type-specific imputation. If there are no prior cell labels provided, scIGANs will pre-cluster the cells using a spectral clustering method. ScIGANs can run on either CPUs or GPUs. The output is the imputed expression matrix of the same dimensions, of which only the true zero values will be imputed without change other expression values.

### Quantitative measurements of single-cell clusters

We used 11 numeric metrics to quantitate the clustering of single cells. RI, the Rand index, is a measure of the similarity between two data clusters. ARI, the adjusted Rand index, is adjusted for the chance grouping of elements. MI, mutual information, is used in determining the similarity of two different clusters of a dataset. As such, it provides some advantages over the traditional Rand index. AMI, adjusted mutual information, is a variation of mutual information used for comparing clusters. VI, the variation of information, is a measure of the distance between two clusters and a simple linear expression involving the mutual information. NVI the normalized VI. ID and NID refer to the information distance and normalized information distance. All these metrics are computed using clustComp() from R package ‘aricode’ (https://cran.r-project.org/web/packages/aricode/). F score (also F1-score or F-measure) is the harmonic mean of precision and recall. AUC, the area under the receiver operating characteristic (ROC) curve, is the probability that a classifier will rank a randomly chosen positive instance higher than a randomly chosen negative one. ACC, accuracy. The above three classification metrics are defined by comparing the independent clustering of cells to the true cell labels. Clustering was done using *prediction()* from the R package SC3 (40). The in-house R scripts for these metrics are provided in the codes for reproducibility (https://github.com/xuyungang/scIGANs_Reproducibility).

### Statistical information

All statistical tests are implemented by R (version 3.6.1). Specifically, the Pearson correlation tests (Figures 4C and S5) were done by the R function *cor.test()* with default parameters; the student’s t-tests (Figures 6C-D and S9) were done by the R function *t.test()* with default parameters; the differentially expressed genes (DEGs) were identified by DESeq2 with the p-adjust <=0.05, log2FoldChange >= 1.5, and baseMean >= 10 (Figures 4A-C and S5); the Spearman correlation were done by the R function *cor()* with default parameters (Figures 7C and S12A).

## RESULTS

### ScIGANs recovers single-cell gene expression from dropouts without inflicting extra noise

Recovery of the biologically meaningful expression from dropout events is the major goal of scRNA-seq imputation to benefit the downstream analyses and biological discoveries. We use both simulated and real scRNA-seq datasets to illustrate the performance and robustness of scIGANs in rescuing dropouts and avoiding additional noise from imputation (Table S1).

First, simulated datasets are used to evaluate the imputation performance since they have known ‘truth’ and can thus benchmark different methods. In a single dataset with a 52.8% zero rate that was simulated according to an independent single-cell clustering method CIDR (29), scIGANs performs superiorly over all other 11 methods in recovering the gene expression and cell population clusters (Figures 2A and S2A; Table S2). Although GANs is a supervised model that requires pre-defined cell labels, we implemented scIGANs to accommodate scRNA-seq data without prior labels, instead to learn the labels by applying spectral clustering (41) on the input data. Trained by the same simulated data without labels (scIGANs w/o), scIGANs slightly reduces the performance but still holds the superiority over the other 11 compared methods, except for scImpute (2) (Figures 2A and S2A; Table S2).

Second, we test the performance of scIGANs and other methods on datasets with different dropout rates simulated by Splatter (30). ScIGANs ranks in the top in rescuing the population clusters (Figures S2B-D) and has the highest resistance to dropout rate increase (Figure S2E; Table S3). Moreover, to evaluate the robustness of imputation methods, we use the same simulation strategy described by SCRABBLE (42) to repeat the above Splatter simulation 100 times for each dropout rate. We evaluate the performance by multiple quantitative clustering metrics (Table S4). The second-ranked SCRABBLE performs superiorly over all other 10 methods; however, it has worse concordance among simulated replicates with a higher dropout rate (Figure 2B). In contrast, scIGANs ranks at the top among all methods and has the most robust performance among the replicates with increasing dropout rates (Figures 2B and S3A-F; Table S4).

Third, we evaluate the imputation methods using a real scRNA-seq dataset from the Human brain, which contains 420 cells in eight well-defined cell types after we exclude uncertain hybrid cells (31). However, the raw data doesn’t show clear clustering of all cell types because of the dropouts and technical noise. After imputation, scIGANs enhances the cell type clusters to the maximum extent so that all 8 cell types could be separated and identified (Figure 2C). Quantitative evaluations of the clustering following different imputation methods highlight the superiority of scIGANs over the others, even trained without the prior cell labels (Figures 2D and S3G; Table S5).

Last, we test another important yet difficult to quantify robustness, i.e. to what extent the imputation method will not introduce extra noise by, for example, mistakenly imputing biological “zeros” or over-imputation. None of the existing imputation methods evaluated their robustness in avoiding extra noise using real scRNA-seq data. Spike-in RNAs (e.g. ERCC spike-in developed by the External RNA Controls Consortium) are a common set of external RNA controls that are equally added to an RNA analysis experiment after sample isolation. It is widely used in scRNA-seq experiments to remove the confounding noises from biological variance. Because the spike-in RNAs are added to samples with the identical amount to capture the technical noise, the readout for the spike-in RNAs should be free of cell-to-cell variability and the detected variances of expression, if exists, should only come from technical confounders other than biological contexts (e.g. cell types). Therefore, the expression of spike-in RNAs that were added to individual cells should not be able to cluster these cells into different subgroups, such as cell types or other biological states. We here use the ERCC spike-in read counts from a real scRNA-seq study (32) to evaluate the imputation methods on denoising the technical variance without introducing extra noise. These 92 ERCC RNAs were added to 288 single-cell libraries of three sets of 96 cells with different cell-cycle states. However, the raw counts failed to cluster these cells into one cluster due to the dropouts of spike-in RNAs (Figure 2E). We expect that the imputation can impute the artificial zeros without exposing the variability of cell states to spike-in profiles and thus all cells should have the same spike-in profiles and will be clustered into a single group. ScIGANs successfully recovers the spike-in profiles with minimum cell-to-cell variability and clustered all cells closely into one group, even though it was trained with supervisory cell labels (Figures 2E and S3H-I). However, other imputation methods suffer from introducing extra noises and thus make the clustering even worse (Figures S3H-I; Table S6). Altogether, scIGANs performs superiorly on imputing the dropouts and avoiding extra noise.

### ScIGANs enables the identification of subcellular states of the homogeneous cell population

Single-cell RNA-seq is typically used to identify different cell types from heterogeneous tissues or cell populations. However, cell populations that seem homogeneous, in terms of expression of cell surface markers, comprise many different cellular states with hidden cell-to-cell variability that can have significant effects on cell function (43,44), such as cellular functions, developmental stages, cell cycle phase, and adjacent microenvironments. Therefore, many biological discoveries require deeper investigation beyond the cell types towards implied cellular states, such as cell-cycle phases of the same cell type. It was reported that cell cycles contribute to phenotypic and functional cell heterogeneity even in monoclonal cell lines (45–47). However, identifying the cell-cycle phases of individual cells from a homogeneous cell population is more challenging for scRNA-seq data due to the prevalence of dropout and high technical variance, which was recently reported more attributable than cell cycle to the single-cell transcriptomic variability (46). We thereby test how imputation could benefit the identification of cell cycle variability from two scRNA-seq studies (Table S1).

First, we reanalyze a scRNA-seq dataset from mouse embryonic stem cells (mESC) that were sorted for G1, S, and G2M phases of the cell cycle (32). Due to the dropout and other technical noise, the raw data does not show cluster structures regarding the three different cell-cycle phases (Figure 3A) and has the poorest clustering measurements (Figure S4A). All other imputation methods fail to recover the cluster structure regarding the cell-cycle states (Figures 3A and S4A; Table S7). Only scIGANs shows significant improvement in detecting cell-cycle states with the best performance (Figures 3A and S4A). Using a collection of independently predefined cell-cycle marker genes from Seurat (33,34), scIGANs significantly improves the identification of the cell cycle states superior over all other methods, shown as the most of sorted cells are correctly assigned in the cell-cycle phase spaces (Figures 3B and S4B).

Second, we assess the performance of different imputation methods on pinpointing the cell-cycle dynamics using a large scRNA-seq data of about 6.8k mouse ESCs (35). The previous work confirmed that ES cells lack strong cell-cycle oscillations in mRNA abundance, but they do show evidence of limited G2/M phase-specific transcription (35). Imputation by scIGANs significantly improved the cell-cycle oscillations with especially a more obvious G2/M phase-specific transcription (Figures 3C and S4C-N). All the above results demonstrate that scIGANs performs better than all other methods on recovering and capturing the subcellular states and very subtle cell-cycle dynamics among single cells of the homogeneous population.

### ScIGANs improves the differential expression analysis

Differential expression analysis refers broadly to the task of identifying those genes with expression levels that depend on some variables, like cell type or state. Ultimately, most single-cell studies start with identifying cell populations and characterizing genes that determine the cell types and drive them different from one to another. Using the scRNA-seq data (36) that have matched bulk RNA-seq data (Table S1), we compare the performances of different imputation methods on improving the identification of differentially expressed genes (DEGs). This dataset has six samples of bulk RNA-seq (four for H1 ESC and two for definitive endoderm cells, DEC) and 350 samples of scRNA-seq (212 for H1 ESC and 138 for DEC). DESeq2 (37) is used to identify DEGs for both bulk and single-cell RNA-seq data between the H1 and DEC cells. The raw scRNA-seq has a much higher zero expression rate than bulk RNA-seq (49.1% vs 14.8%) and shares fewest DEGs with bulk samples (Figure 4A). After imputation, the number of DEGs is increased toward the DEGs numbers of bulk samples (except the four other deep learning-based methods, AutoImpute (48), DCA (6), DeepImpute (8), and scGAIN (49), which detect much fewer DEGs than raw data). Especially highlighted by the yellow and green bars in Figure 4A, the AutoImpute and scGAIN detect significantly fewer DEGs and show the poorest agreement with other methods. In contrast, scIGANs imputation identifies the highest number of dataset-specific DEGs and shares a significant number of DEGs with bulk RNA-seq (Figure 4A). To more fairly quantitate the performance of DEG detection using scRNA-seq data, we define the accuracy (ACC), F score (also F1-score or F-measure), and the area under the receiver operating characteristic curve (AUC) for each DEG detection from the imputed scRNA-seq datasets, by taking DEGs from bulk RNA-seq as the gold standard (refer to Methods for details). The overall performance was defined as the average of the above three measurements. ScIGANs ranks in second place with scImpute, slight behind VIPER (Figure 4B).

Additionally, we use a set of top 1000 DEGs from bulk samples (500 up-regulated and 500 down-regulated genes) as a benchmark to evaluate the correspondence of DEG detection between single-cell and bulk RNA-seq data. Without exception, scIGANs-imputed scRNA-seq data show the highest correspondence with bulk RNA-seq, depicted as the most number of shared top 1000 DEGs and highest correlation of the fold-changes versus bulk RNA-seq (Figures 4C and S5). Moreover, the expressions of five marker genes for H1 and DEC cells are investigated to compare the extent to which the imputation methods recover the expression patterns of signature genes. Results show that scIGANs best reflect the expression signatures of both H1 and DEC cells by removing undesirable variation resulted from dropouts (Figures 4D and S6). Projection of cells to the UMAP space overlaid by the expression of signature genes furtherly highlights the performance of scIGANs on recovering the expression patterns of signature genes (Figures 4E-F and S7). In summary, scIGANs improves the identification of DEGs from scRNA-seq data with the best performance among other competing methods.

### ScIGANs enhances the inference of cellular trajectory

Beyond characterizing cells by types, scRNA-seq also largely benefits organizing cells by time-course or developmental stages, i.e. cellular trajectory. In general, trajectory analysis starts with reducing the dimensionality of the expression data, then reconstructs a trajectory along which the cells are presumed to travel, and finally projects each cell onto this trajectory at the proper position. Although single-cell experiments can illuminate trajectories in a wide variety of biological settings (38,50–52), none of the single-cell trajectory inference methods account for dropout events. We speculate that inferring the cellular trajectory on scRNA-seq data after imputation could improve the accuracy of pseudotime ordering. We utilize a time-course scRNA-seq data derived from the differentiation from H1 ESC to definitive endoderm cells (DEC) (36). A total of 158 cells were profiled at 0, 12, 24, 36, 72, and 96 hours after inducing the differentiation from H1 ESCs (Figure 5A and Table S1). We apply scIGANs and all other 11 imputation methods to the raw scRNA-seq data with known time points and then reconstruct the trajectories using Mococle3 (38). Imputation by scIGANs produces the highest correspondence between the inferred pseudotime and real-time course (Figures 5B-C and S8), suggesting that scIGANs recovers more accurate transcriptome dynamics along the time course. We also investigate the signature genes of pluripotency (e.g. NANOG and POU5F1) and DECs (e.g. CER1 and HNF1B) and find that scIGANs improves the gene expression dynamics after imputation (Figures 5D-E) and has better performance than other imputation methods (Figure S8). These results demonstrate that scIGANs can help to improve the single-cell trajectory analysis and recover the temporal dynamics of gene expression.

### ScIGANs is robust to the small dataset of few genes with low expression or cell-to-cell variance

In general, other imputation methods (e.g. SAVER (3) and scImpute (2)) heavily rely on a set of pre-selected informative genes that are highly expressed and unlikely to suffer from the dropout. Imputation is then performed from the most similar cells defined by these informative genes. In contrast, scIGANs automatically learns the gene-gene and cell-cell dependencies from the whole dataset. More important, scIGANs converts each single-cell expression profile to an image so that a 1-dimension “feature” vector is reshaped to a 2-dimension matrix with each element representing the expression of a single gene (Figure S1A). Like image processing, scIGANs is then trained by convolution on the matrix so that the 2-dimension gene-gene relations within each cell are captured. Therefore, we hypothesize that scIGANs is more robust to genes of low expression or with less cell-to-cell variance.

From the aforementioned scRNA-seq data with 350 cells (212 H1 ESC and 138 DEC) (36), we randomly sample small sets of genes (n=1024 for each) from the 5000-gene sets with top/lower means or variances, as well as a set of 1024 genes randomly picked from all expressed genes (Table S1; refer to Methods for details). Visualizations based on the 1024 genes (only ∼5% of detected genes) with very low expression or variance show that the two types of cells are almost mixed up without any cluster characterization for the raw expression profiles (Figures 6A and 4E). Imputation by scIGANs successfully recovered the two cell clusters for both datasets with only 1024 genes of low expression and variance, respectively (Figure 6B). However, all other methods failed in identifying the two cell types from these datasets (Figure S9). Moreover, scIGANs significantly changes the mean and variance of expression after imputation, while they are not always the same cases for other methods (Figures 6C-D and S9). All these results show that scIGANs is robust to a small dataset composed of very few genes (∼5% of detected genes) with very low expression or cell-to-cell variance, which are less informative for other imputation methods. It’s the strong support to the expectation that scIGANs can learn very limited gene-gene and cell-cell dependencies from a small set of lowly or close-to-uniform expressed genes.

### ScIGANs is scalable to scRNA-seq methods and data sizes

The last but not least concern on scRNA-seq imputation raises in terms of the scalability to data sizes and sequencing methods, since the scRNA-seq becomes increasingly high-throughput in cell numbers and available on different protocols/platforms. To evaluate scIGANs and compare it with other methods regarding this concern, we collected a set of real scRNA-seq data to test the scalability to data sizes and sequencing methods (Table S1).

We first test how scIGANs works equally well on scRNA-seq data from different single-cell methods. We use the datasets from three 3 human lung adenocarcinoma cell lines (H2228, H1975, HCC827) generated by 10X Genomics/Chromium, CEL-seq2/Fluidigm, and Drop-seq/Droplet, respectively, which represent two single-cell platforms, i.e. droplet-based and plate-based. The cell clusters from raw data with dropouts show that 10X Genomics/Chromium generate the best outcomes while CEL-seq2/Fluidigm and Drop-seq/Droplet are more affected by the dropouts (Figures 7A and S10), which is consistent with the observation from a very recently publish benchmarking work (34). Imputation is expected to attenuate the dropout effects between different scRNA-seq methods; however, the performance varies largely across imputation methods. ScIGANs is among the four top-ranked methods showing a minor difference in the performances across different sequencing methods (Figures 7B and S11; Table S8). To further test how imputation could improve the correspondence of single-cell gene expression profiled by different sequencing methods, we calculated the Spearman correlation coefficient from the expression profiles of the same cell type generated by different sequencing methods (refer to Methods for details). With a high agreement with the clustering metrics (Figures 7B and S11; Table S8), scIGANs is top-ranked in recovering the highly correspondent expression profiles of the same cell types generated from different scRNA-seq methods (Figures 7C and S12A). These results demonstrate that scIGANs shows equally high performance on scRNA-seq datasets generated by different scRNA-seq methods (protocols/platforms).

Second, we assess the computational intensity of imputation methods using five datasets of 1000 (1k), 5000 (5k), 25000 (25k), 50000 (50k), and 100000 (100k) cells created by random sampling from the PMBC 10k dataset (Table S1, Methods). As shown in Table S9, many methods, including SAVER and scImpute, failed the jobs on the datasets with 50k or more cells because of the high requirements of memory (> 64GB). Specifically, SCRABBLE failed on all datasets because of requiring more than 48 hours, even on the 1k dataset. Only scIGANs, DeepImpute, and MAGIC complete all jobs on five datasets. Among these three outstanding methods, scIGANs shows comparable scalability to data sizes in terms of the running time and memory usage, especially when accelerated by GPU (Figures 7D and S12B; Table S9). Therefore, superior over most of the other tested methods, scIGANs can impute the scRNA-seq dataset with more than 100k cells, which covers the throughputs of most scRNA-seq studies. To sum up, scIGANs is scalable to data sizes and scRNA-seq methods and promises equal performance.

## DISCUSSION

Here we propose the generative adversarial networks for scRNA-seq imputation (scIGANs). ScIGANs converts the expression profiles of individual cells to images and feeds them to generative adversarial networks. The trained generative network produces expression profiles representing the realistic cells of defined types. The generated cells, rather than the observed cells, are then used to impute the dropouts of the real cells. We assess scIGANs regarding its performances on the recovery of gene expression and various downstream applications using simulated and real scRNA-seq datasets. We provide compelling evidence that scIGANs performs superior over the other 11 peer imputation methods. Most importantly, using generated rather than observed cells, scIGANs avoids overfitting for the cell type of big population and meanwhile promise enough imputation power for rare cells.

While there are many methods for scRNA-seq imputation, we specifically show how the GANs can improve the imputation and downstream applications, representing one of three pioneering applications of GANs to genomic data. Two other recent manuscripts used GANs to simulate (generate) realistic scRNA-seq data with the applications of either integrating multiple scRNA-seq datasets (18) or augmenting the sparse and underrepresented cell populations in scRNA-seq data (26). We, for the first time, advance the applications of GANs to scRNA-seq for dropout imputation. Inspired by the great success of GANs in inpainting and highly relevant work that applied GANs for ‘realistic’ generation of scRNA-seq data (26), we speculate that the generated realistic cells can not only augment the observed dataset but also benefit the dropout imputation since it was proved that the generated data mimics the distribution of the real data in their original space with stable fidelity (26). Our multiple downstream assessments and applications on simulated and real scRNA-seq datasets demonstrated its advantage in dropout imputation, superior over other peer methods. Especially for cells coming from very small populations, generated data were proved to faithfully augment the sparse cell populations (26) and thus reduce the sampling bias and improve the imputation power, which, however, are suffered by all other imputation methods. Additionally, GANs can learn dependencies between genes beyond pairwise correlations (26), which enables scIGANs more sensitive and robust to small datasets with very low or uniform expressions. We demonstrated these advantages by ERCC spike-in RNAs (Figures 2E and S3H-I) and downsampling real scRNA-seq data (Figures 6 and S9).

The underlying basis of scIGANs is that the real scRNA-seq data is derived from sampling, which doesn’t have enough cells to characterize the true expression profiles of each cell type, even for the major cell populations; and the generated realistic cells could augment the observations, especially for sparse and underrepresented cell populations, and thus improve the dropout imputation of scRNA-seq data. There are many benefits of using realistic rather than the observed cells for imputation. First, the generated cells characterize the expression profiles of real cells, and faithfully represent the cell heterogeneity. Therefore, the realistic cells are ideal to serve as extra samples and independently impute the observed dropouts to avoid the “circular logic” issue (overfitting) suffered by other methods (e.g. scImpute), which borrow information from the observed data per se. Second, the realistic cells will augment the rare cell-types, and thus overcome potential sampling biases to avoid imputation performance skewed to dominant cell populations. Additionally, benefitting from the power of GANs in adversarially discriminating between real and realistic data, and the augmentation from generated data, scIGANs is more sensitive to subcellular states like the cell-cycle phases investigated in this study. Imputation by scIGANs enables the investigation of scRNA-seq data beyond the identification and characterization of cell types but go deeper into subcellular states and capture cell-to-cell variability of the homogenous cell populations. This is critical for the applications of scRNA-seq to pinpoint the state transitions along the cellular trajectory or identify and remove the subcellular confounding factors (e.g. cell-cycle phases) (46). Our evaluations on cell-cycle phase detection and trajectory construction show the superiority of scIGANs over all other 11 tested methods.

During the submission of this work, another work using GANs for scRNA-seq imputation (referred to as scGAIN) was posted in BioRxiv (49). We added it in all evaluations and downstream analyses in our work and found that scGAIN is not as good as scIGANs in all evaluations. We, therefore, want to specifically discuss the reasons for this difference.

First, scGAIN and scIGANs use different network architectures in their GANs. ScIGANs uses convolutional neural networks (CNN) while scGAIN uses full connected networks (FCN). FCN may hold good performance on high-quality data with low or no dropout and noise. However, we focus on detecting and imputing the dropouts. For this purpose, FCN will not promise good performance. Instead, through multi-layer networks, convolutional neural networks (CNN) will map the features to adjacent positions, thereby forming representative local features and being captured by subsequent networks. This idea is widely used in stock market prediction (53,54). Specifically, in scRNA-seq data imputation, we extract high-quality information from features by concatenating a pooling layer to each convolutional layer. We use max pooling, which means to retain the highest expressions from the local area of the feature space. After multilayer mapping of the feature space, the effect of dropout in a local area will be attenuated by the high-quality information (high expression). Therefore, CNN with convolution-pooling architecture will automatically filter out the dropouts and noise and not compromise feature extraction. This explains why scIGANs works better than scGAIN and other methods in identifying subcellular states (Figures 3 and S4), such as the cell-cycle phase, which always is implied and confounded with technical noise in scRNA-seq data.

Second, during either training or imputation steps, scGAIN requires a mask matrix to identify which entries in the input matrix are targets for imputation, which was pre-defined by a hard threshold. In contrast, the expression matrix is the only required input for scIGANs. Once trained, scIGANs uses the KNN algorithm to heuristically define and impute dropouts. The uniform expression cutoff for dropouts may largely compromise the performance since genes have different expression patterns among cells. This is why scGAIN has the worst performance in DEG detection, which heavily relies on the expression patterns across cells (Figures 4A-B).

Third, along with the changes in the generator and discriminator models from a standard GAN, scGAIN uses different loss functions. The discriminator no longer takes two sets of samples (real or fake); instead, the discriminator distinguishes between observed and imputed portions (probability) of that sample. In contrast, scIGANs takes each cell as a whole and distinguishes the fake from real based on the overall expression profile of each cell rather than individual genes. Additionally, scIGANs reshapes the expression profiles of individual cells into a 2-dimension “images” and takes these “images” rather than the original 1-dimension vectors as samples for training. In this way, scIGANs can learn the non-linear gene-gene dependencies from complex and multi-cell type samples, which enable scIGANs more robust to a small set of genes with very low expression or cross-cell variance (Figures 6 and S9).

In summary, scIGANs is a method that takes advantage of both the gene-to-gene and cell-to-cell relationships to recover the true expression level of each gene in each cell, removing technical variation without compromising biological variabilities across cells. ScIGANs is also compatible with other single-cell analysis methods since it does not change the dimension (i.e., the number of genes and cells) of the input data and it effectively recovers the dropouts without affecting the non-dropout expressions. Additionally, ScIGANs is robust to small datasets that have few genes with low expression and/or cell-to-cell variance. Last but not least, scIGANs is also scalable to data sizes and works equally well on datasets generated by different scRNA-seq protocols/platforms. Altogether, scIGANs is not only an application of GANs in omics data but also represents a competing imputation method for the scRNA-seq data.

## AVAILABILITY

ScIGANs is an open-source tool available in the GitHub repository with a usage tutorial (https://github.com/xuyungang/scIGANs). The sources and pre-processes of all data are described in Methods. The processed datasets and codes used to reproduce the Figures and Tables are available at GitHub (https://github.com/xuyungang/scIGANs_Reproducibility).

## AUTHOR’S CONTRIBUTIONS

YX, ZZ, and XZ conceived the study. ZZ developed the scIGANs model and YX wrapped it up to a package. YX analyzed all scRNA-seq datasets, interpreted the results, and wrapped up the reproducibility codes on GitHub. ZZ, LY, JL, and ZF helped to test the methods and reproduce the analyses. YX wrote the manuscript and all authors revised it. All authors read and approved the final version of the manuscript.

## SUPPLEMENTARY DATA

Supplementary Data are available online.

Additional file 1 (PDF): Figures S1-S12.

Additional file 2 (PDF): Tables S1-S3, and Tables S5-S9.

Additional file 3 (XLSX): Table S4.

