## Additional file 1: Figures S1-S12 for "scIGANs: single-cell RNA-seq imputation using generative adversarial networks"

**Figure S1 The schematic view of scIGANs. A.** The architecture of Generative Adversarial Networks trained using the single-cell RNA-seq expression matrix. **B.** Impute the scRNA-seq expression based on the generated expression matrix using the mean of k-nearest neighbors. Refer to Methods for a detailed description. Numbers in parentheses indicate the dimensions.

**Related to Figure 1.**

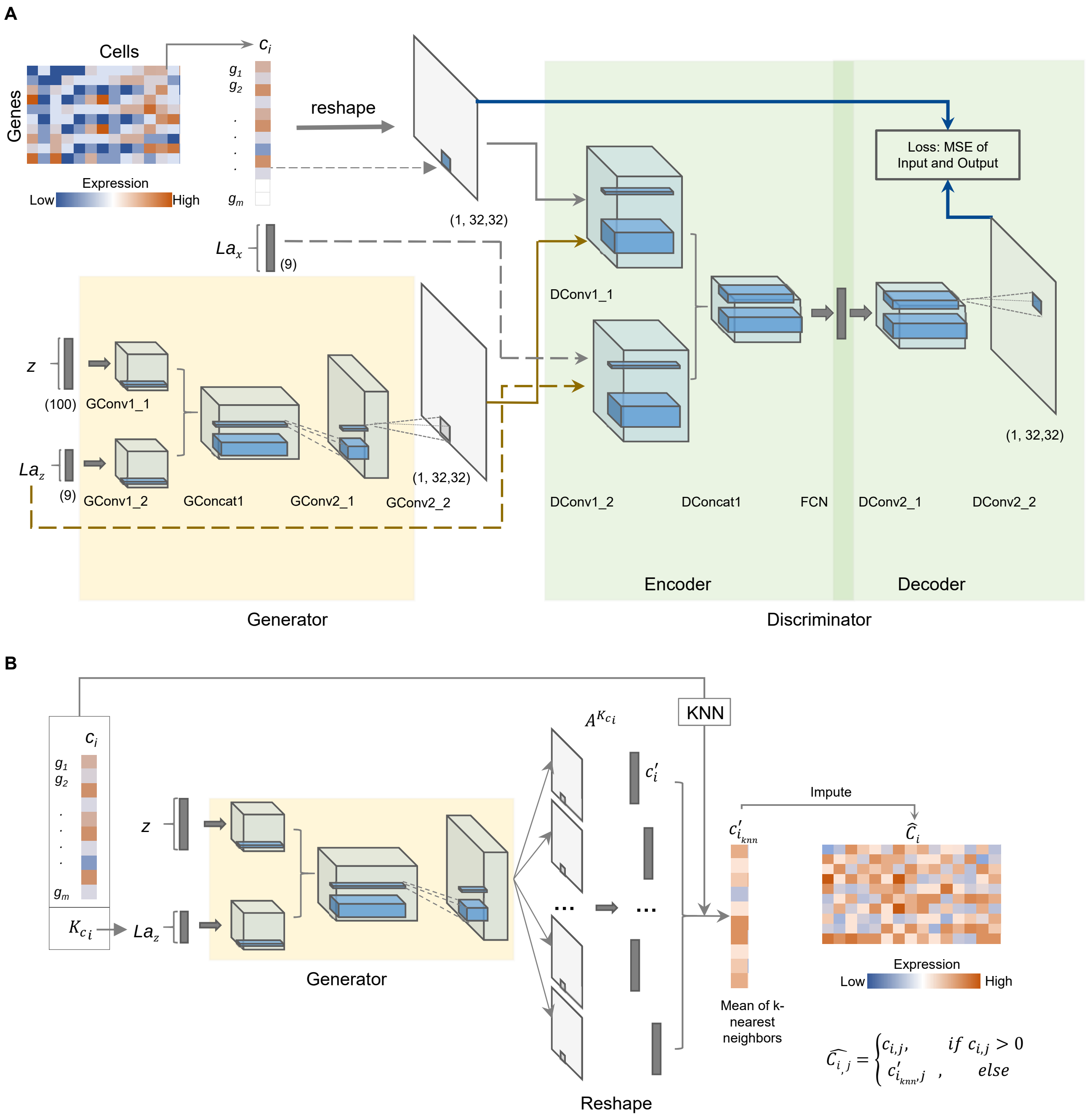

**Figure S2. Performance evaluation using simulated data. A.** Multiple metrics are used to measure the clustering outputs for each of the compared methods on CIDR simulated data. ARI, adjusted rand index; NMI, normalized mutual information; F score, the harmonic mean of precision and recall; AUC, area Under the Curve of ROC (receiver operating characteristics); ACC, accuracy. The full list of all considered clustering metrics is provided in **Supplementary Table S2**. **B-D.** The UMAP plots of the Splatter simulated data with three different dropout rates (71%, 83%, and 87%). The full list of all considered clustering metrics is provided in **Supplementary Table S3**. **E.** scIGANs is robust to the dropout rate increase, shown as the adjusted rand index (ARI).

**Related to Figure 2.**

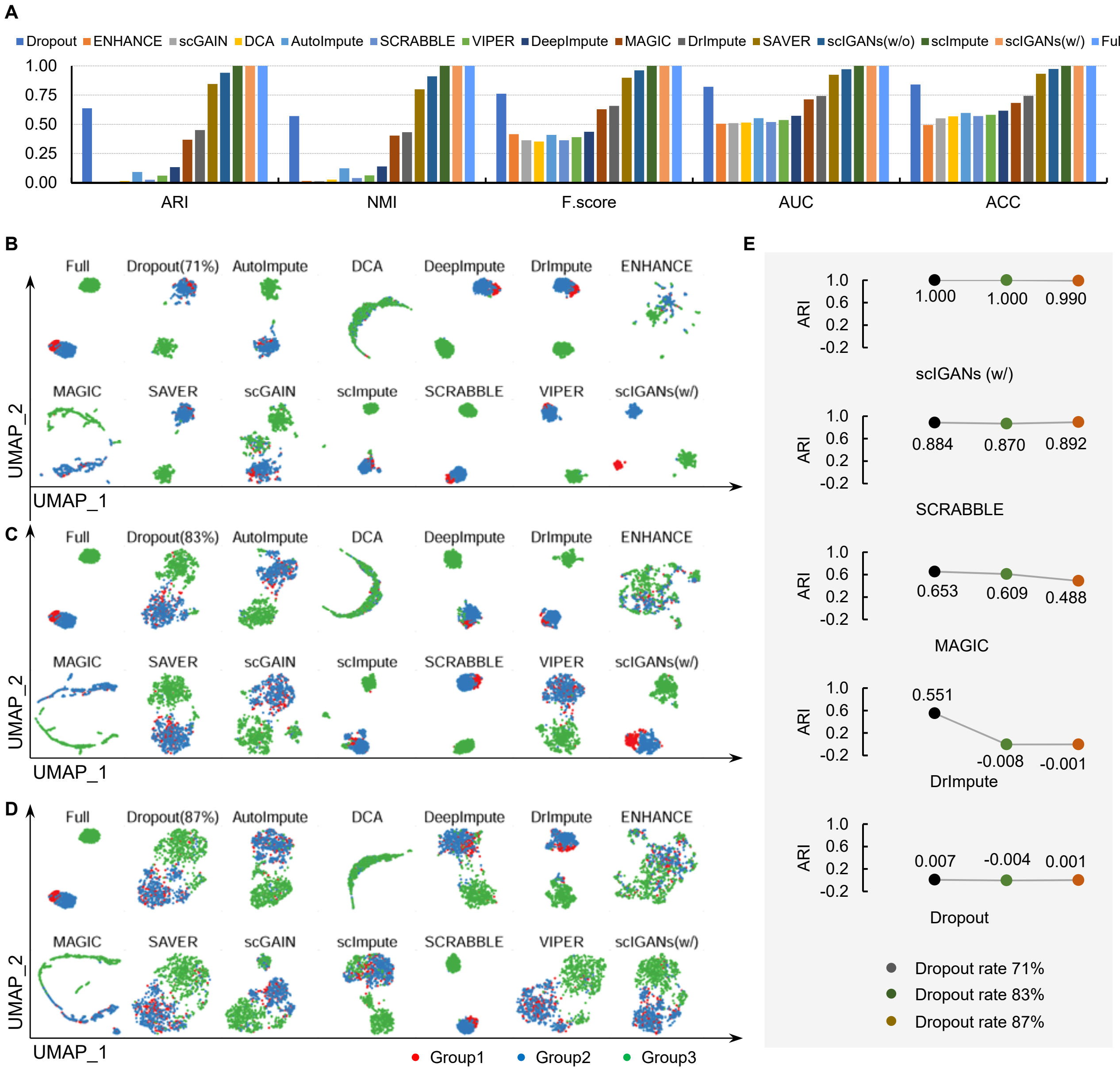

**Figure S3. Robustness evaluation using simulated and real scRNA-seq data. A-F.** Multiple metrics are used to measure the clustering output for each of the compared methods on Splatter simulated data (100 times for each dropout rate). RI, rand index; NMI, normalized mutual information; MI, mutual information; F score, the harmonic mean of precision and recall; AUC, area Under the Curve of ROC (receiver operating characteristics); ACC, accuracy. The full list of all considered clustering metrics is provided in **Supplementary Table S4**. **G.** The UMAP plots of the real scRNA-seq data from the human brain. Clustering measures are provided in **Supplementary Table S5**. The color scheme for cell types is the same as in Figure 2C. **H.** The UMAP plots of the real scRNA-seq data for ERCC spike-in RNAs. Cluster measures are provided in **Supplementary Table S6**. I. Cell-to-cell variabilities before and after imputation by different methods represented as the coefficient variances (CV) of all 92 spike-in RNAs across 288 cells. The vertical dotted line indicates the median CV of scIGANs trained with cell labels.

**Related to Figure 2.**

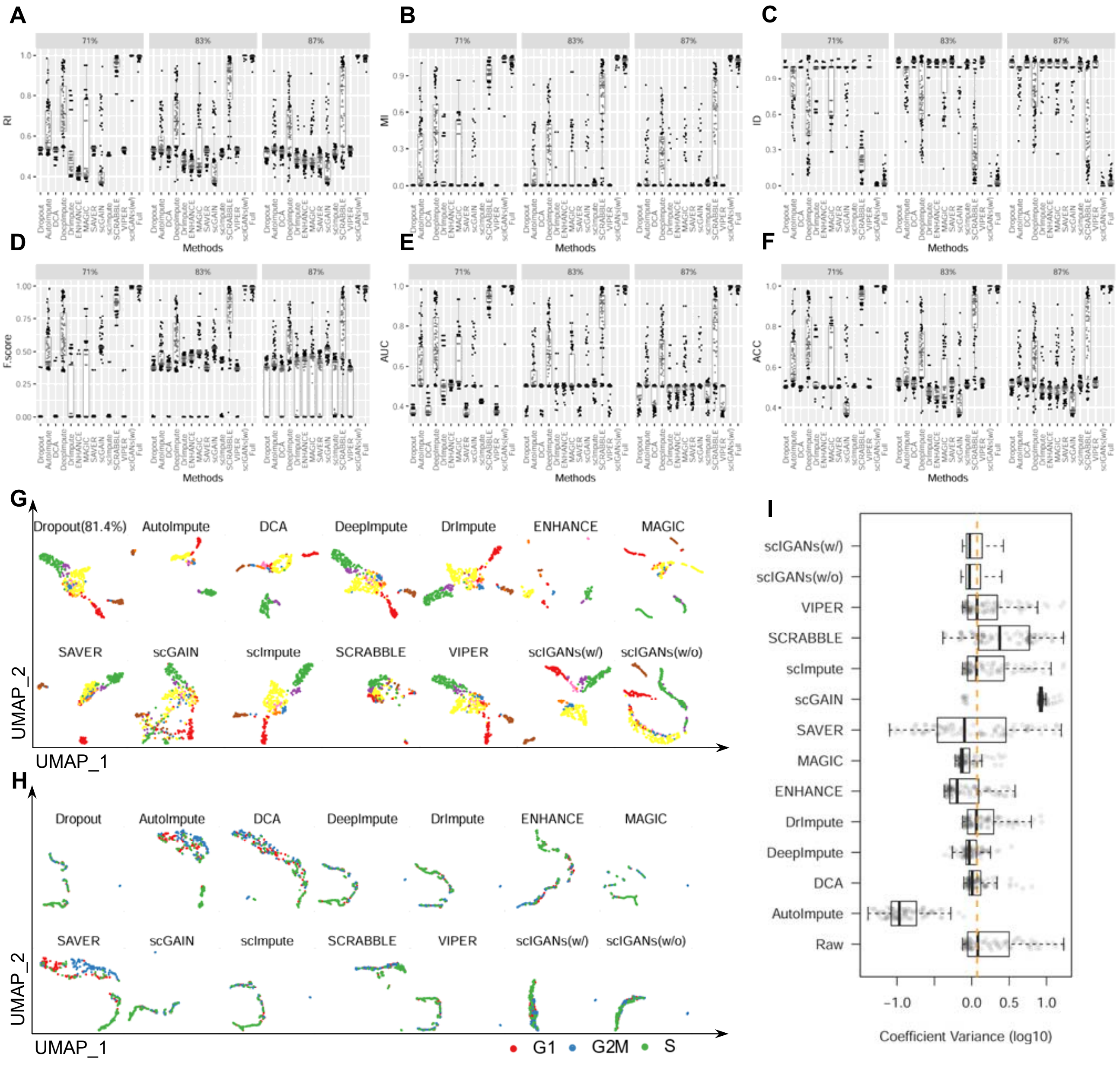

**Figure S4. Identification of subcellular stats of the homogeneous cell population. A.** Multiple metrics are used to measure the clustering output for each of the compared methods on cell-cycle data. The full list of all considered clustering metrics is provided in **Supplementary Table S7**. **B.** Cells are projected to the cell-cycle phase spaces based on a collection of cell-cycle marker genes. **C-N.** Full dynamic cell-cycle profiles shown as the hierarchical clustering of Pearson correlations among 44 cell-cycle-regulated genes across ~6.8k mouse ESCs from the scRNA-seq data before and after imputation by different methods. The barplots show the quantitative concordance between the assigned cell-cycle phases by hierarchical clustering and the true phases that these genes serve as markers. F score, the harmonic mean of precision and recall; AUC, area Under the Curve of ROC (receiver operating characteristics); ACC, accuracy.

**Related to Figure 3.**

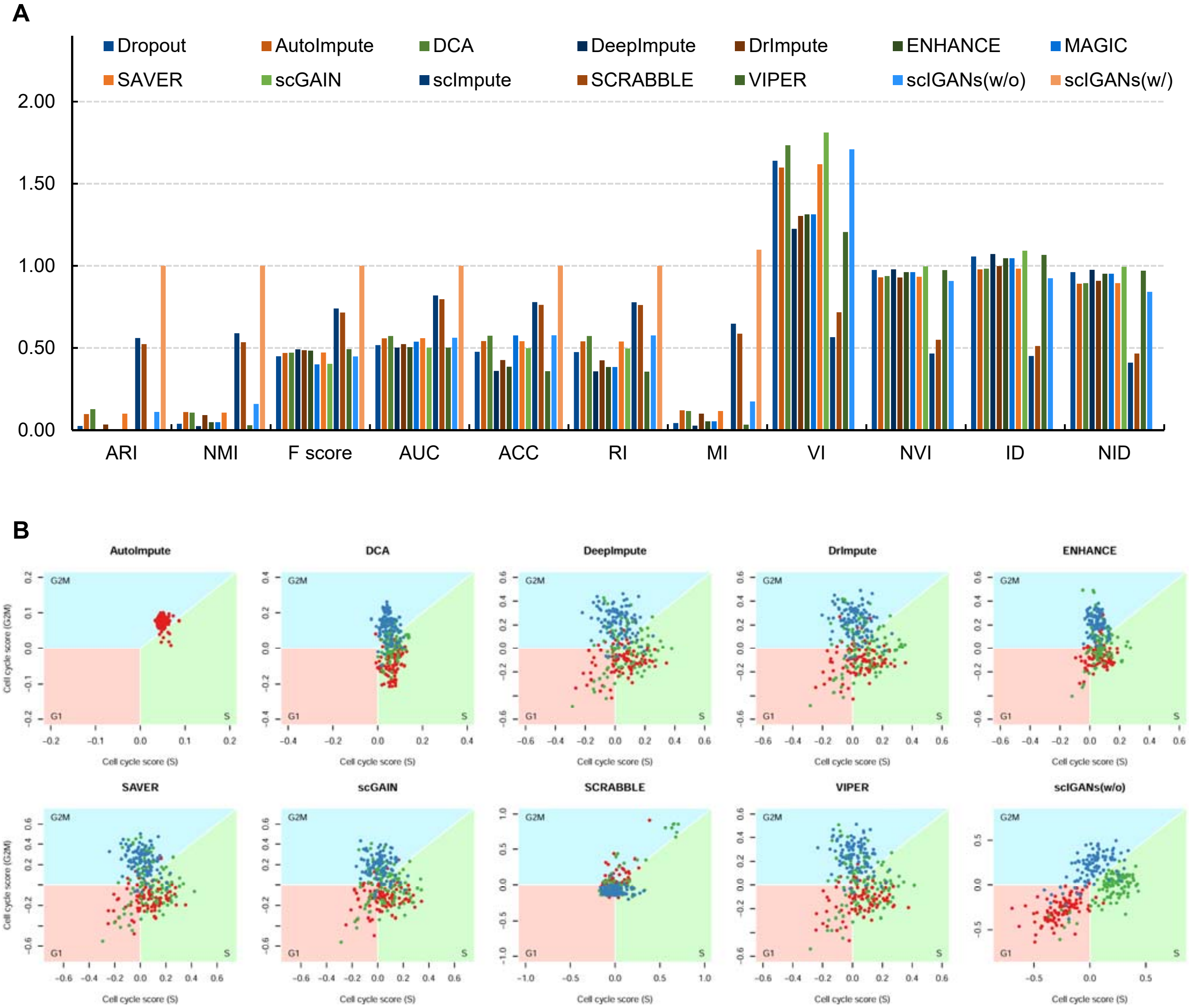

**Figure S4 (continue)**

**C. Raw data**

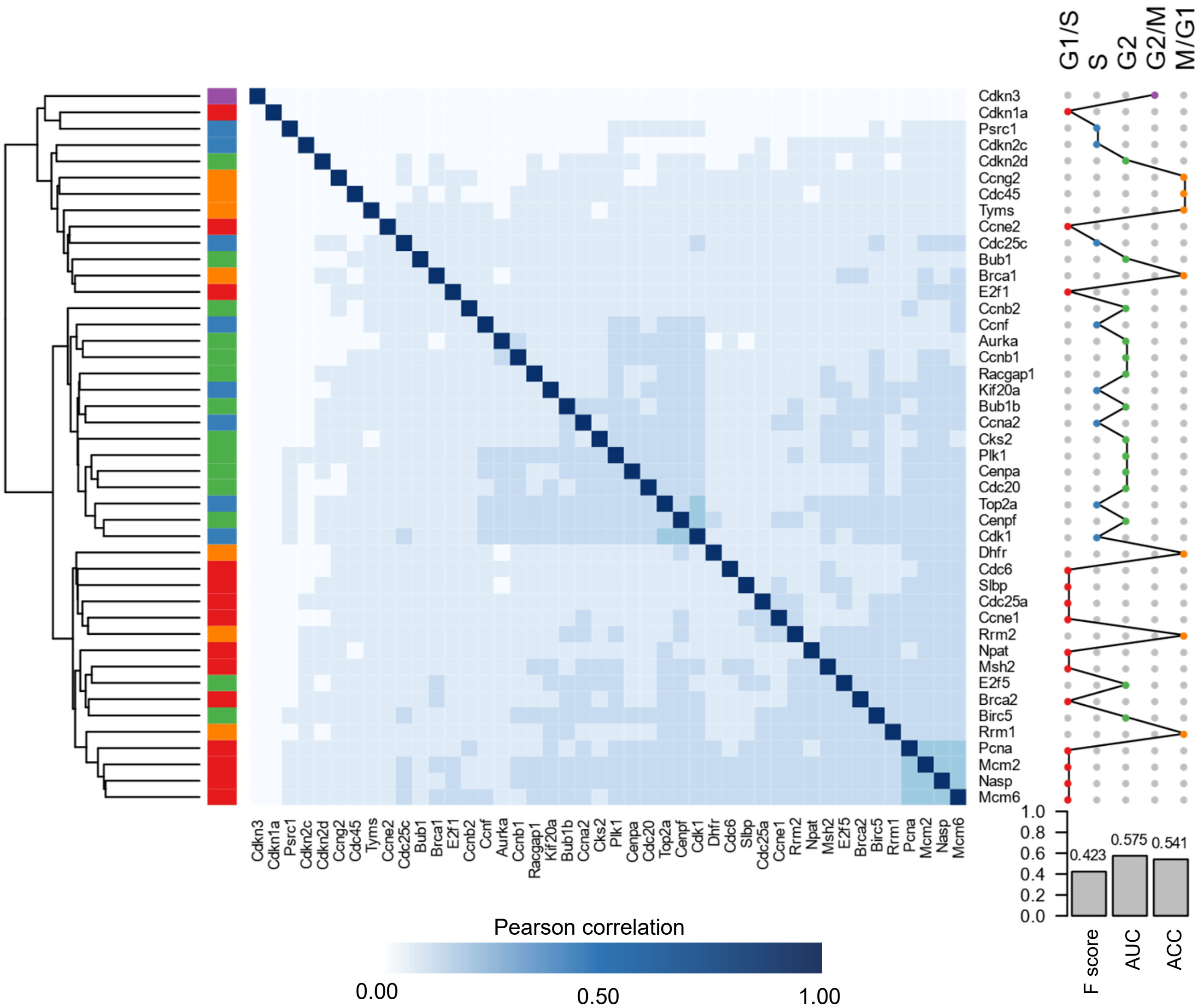

**Figure S4 (continue)**

**D. Imputed by scIGANs (w/)**

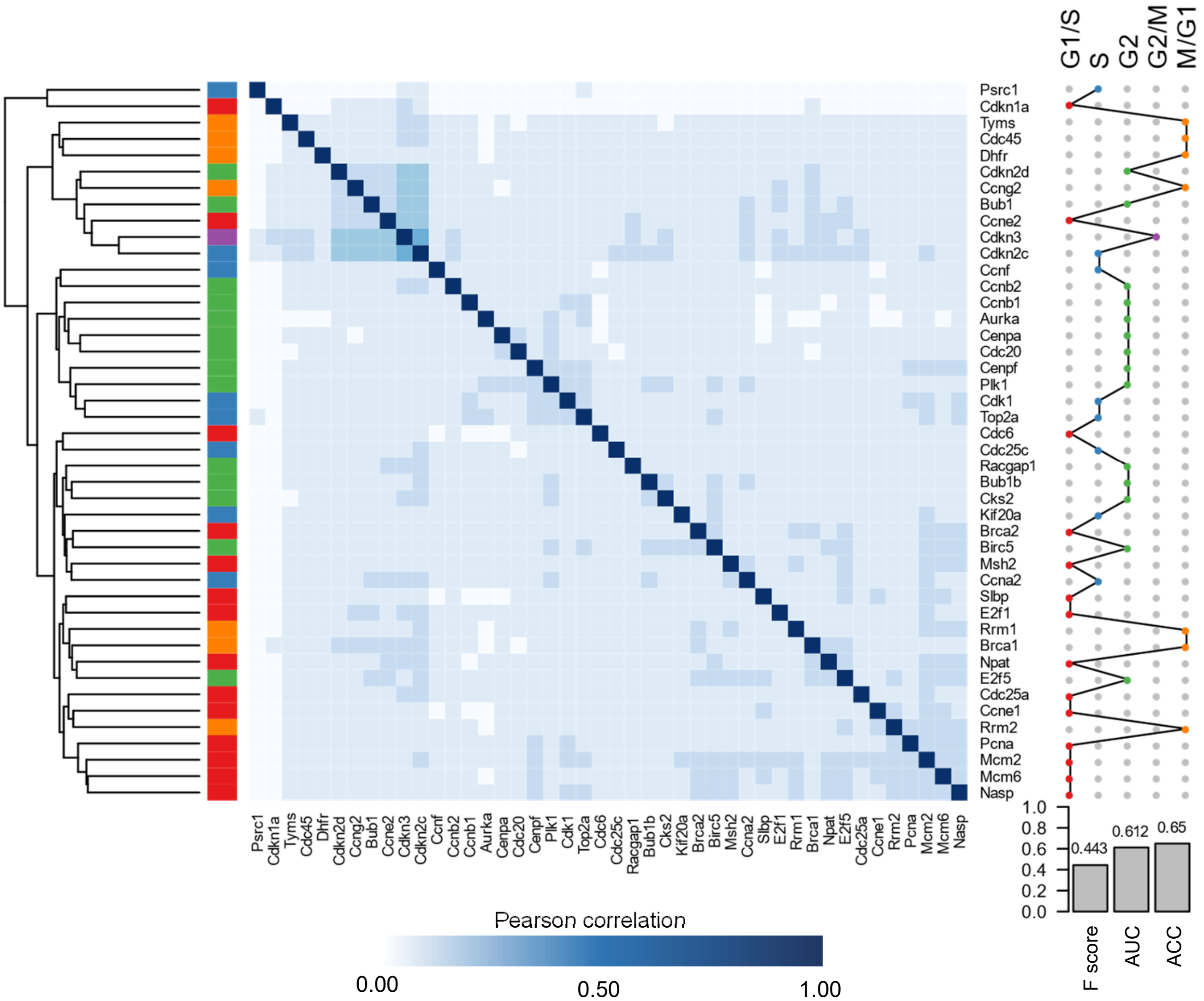

**Figure S4 (continue)**

**E. Imputed by scIGANs (w/o)**

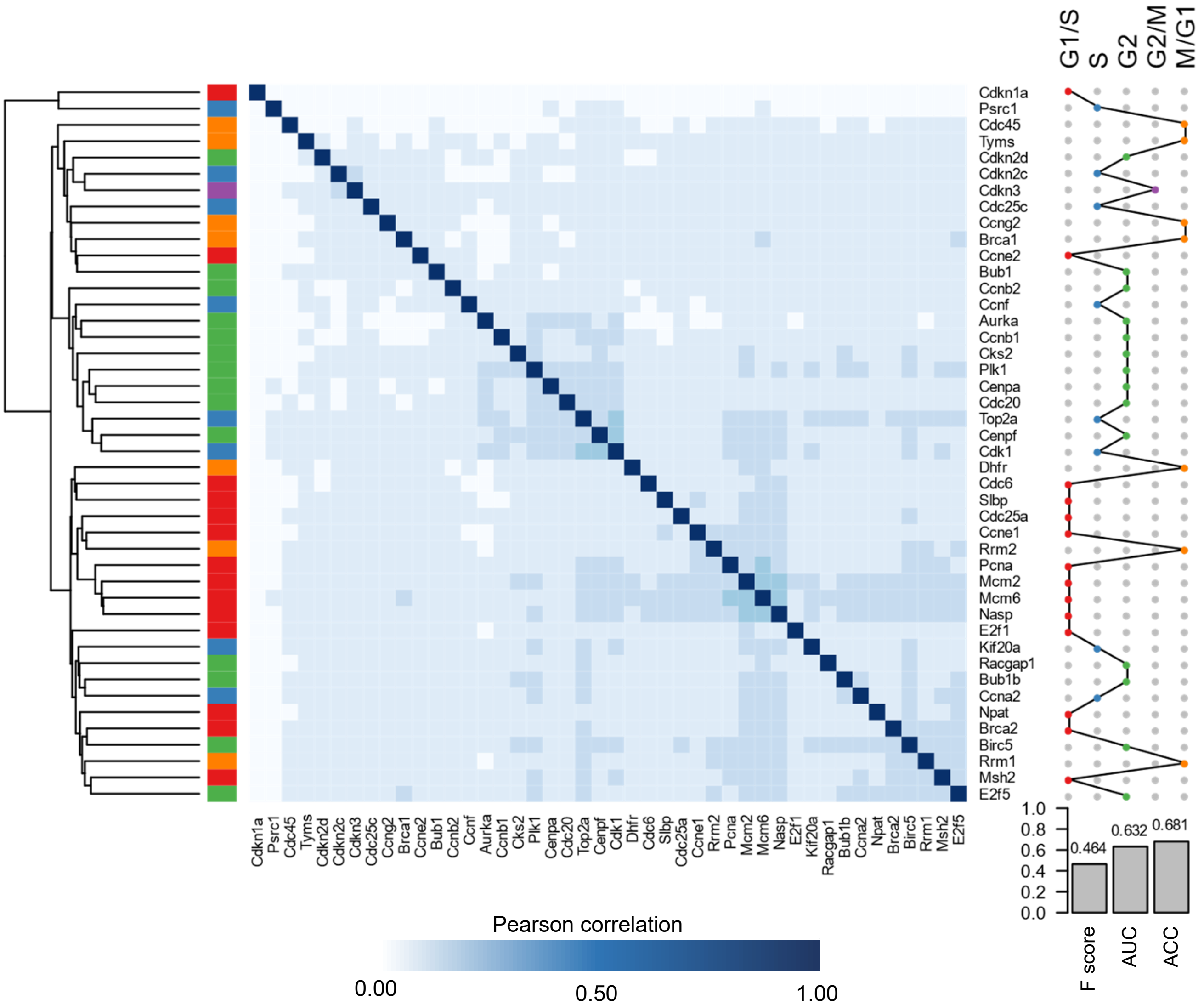

**Figure S4 (continue)**

**F. Imputed by DCA**

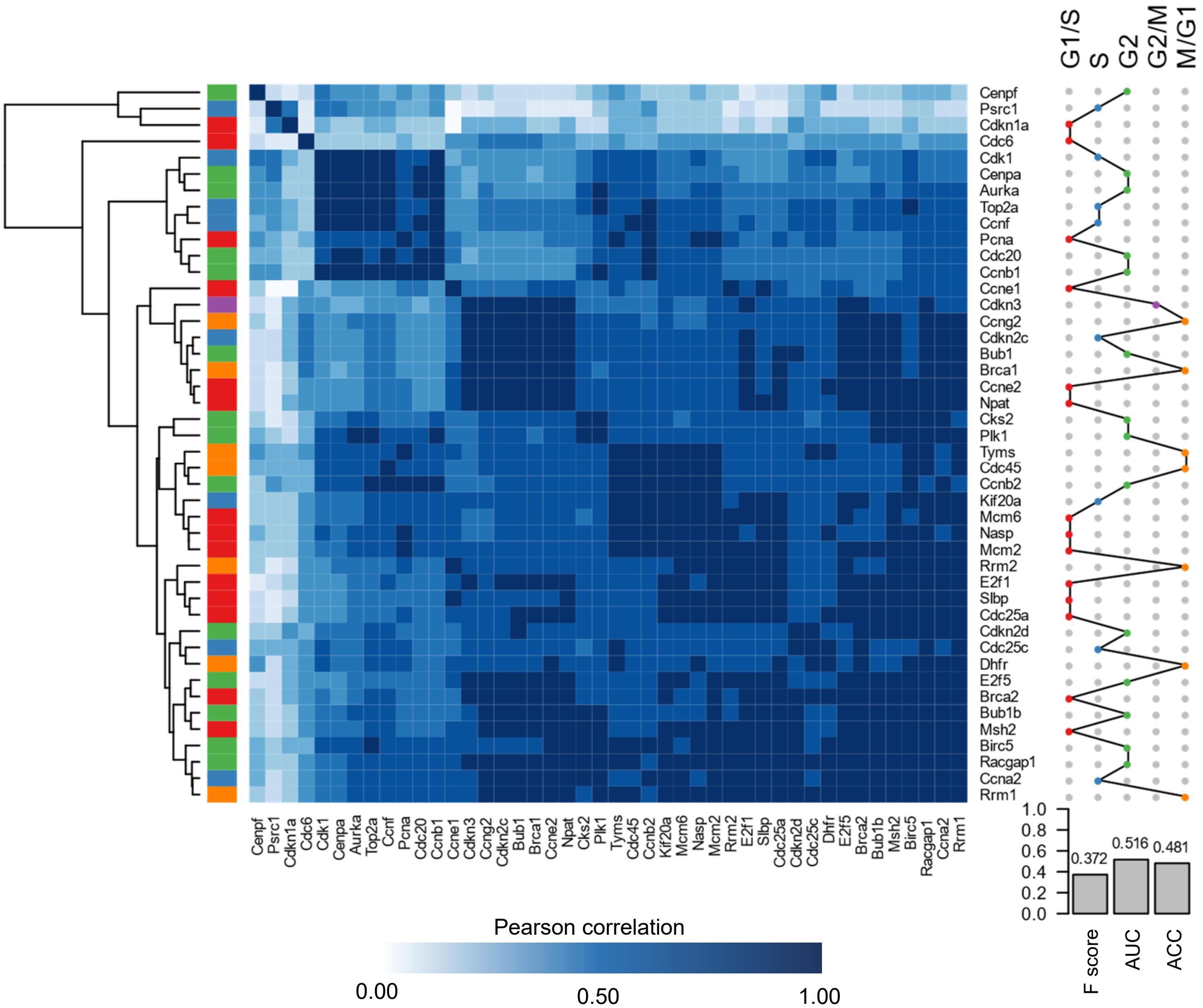

**Figure S4 (continue)**

**G. Imputed by ENHANCE**

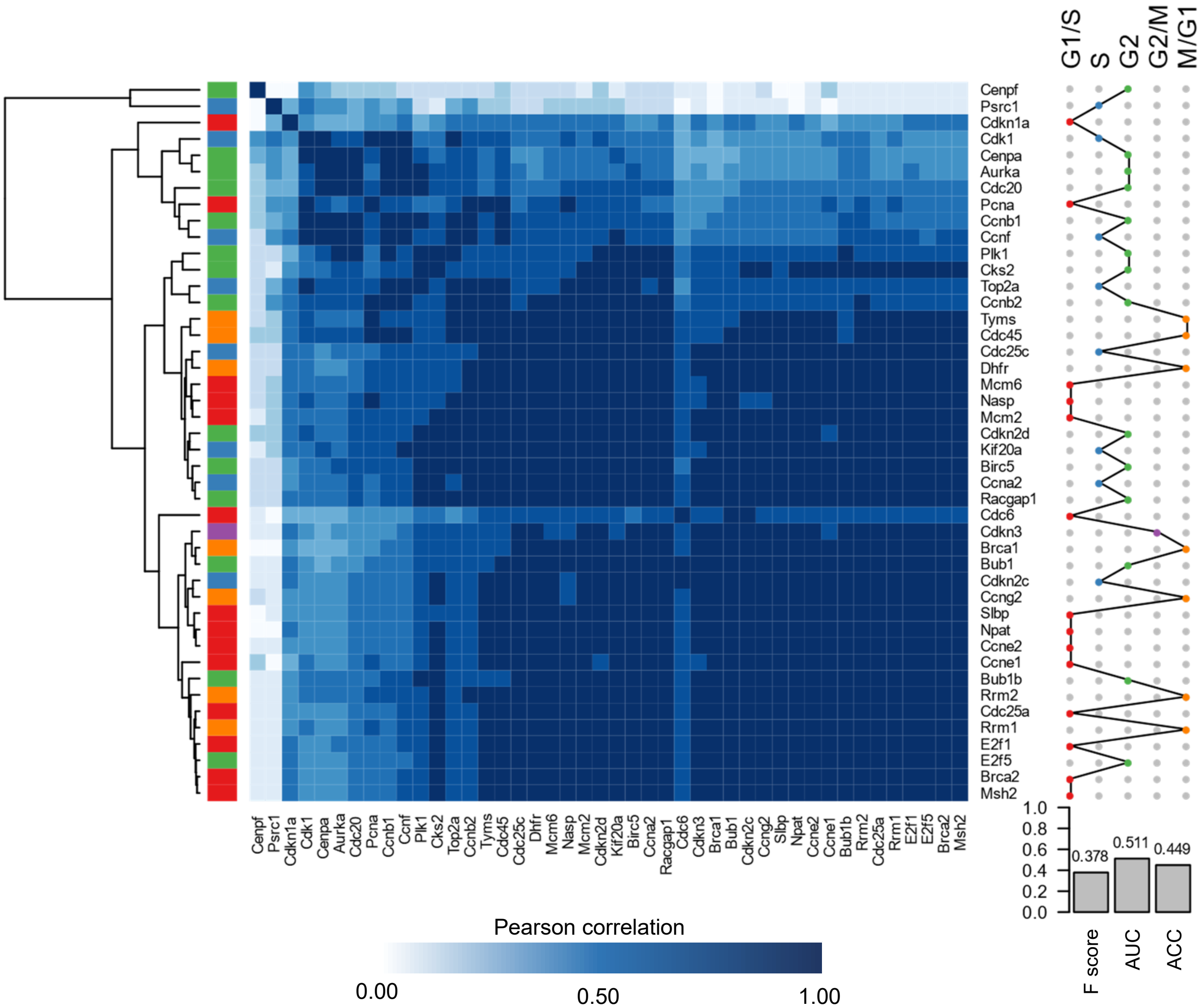

**Figure S4 (continue)**

**H. Imputed by MAGIC**

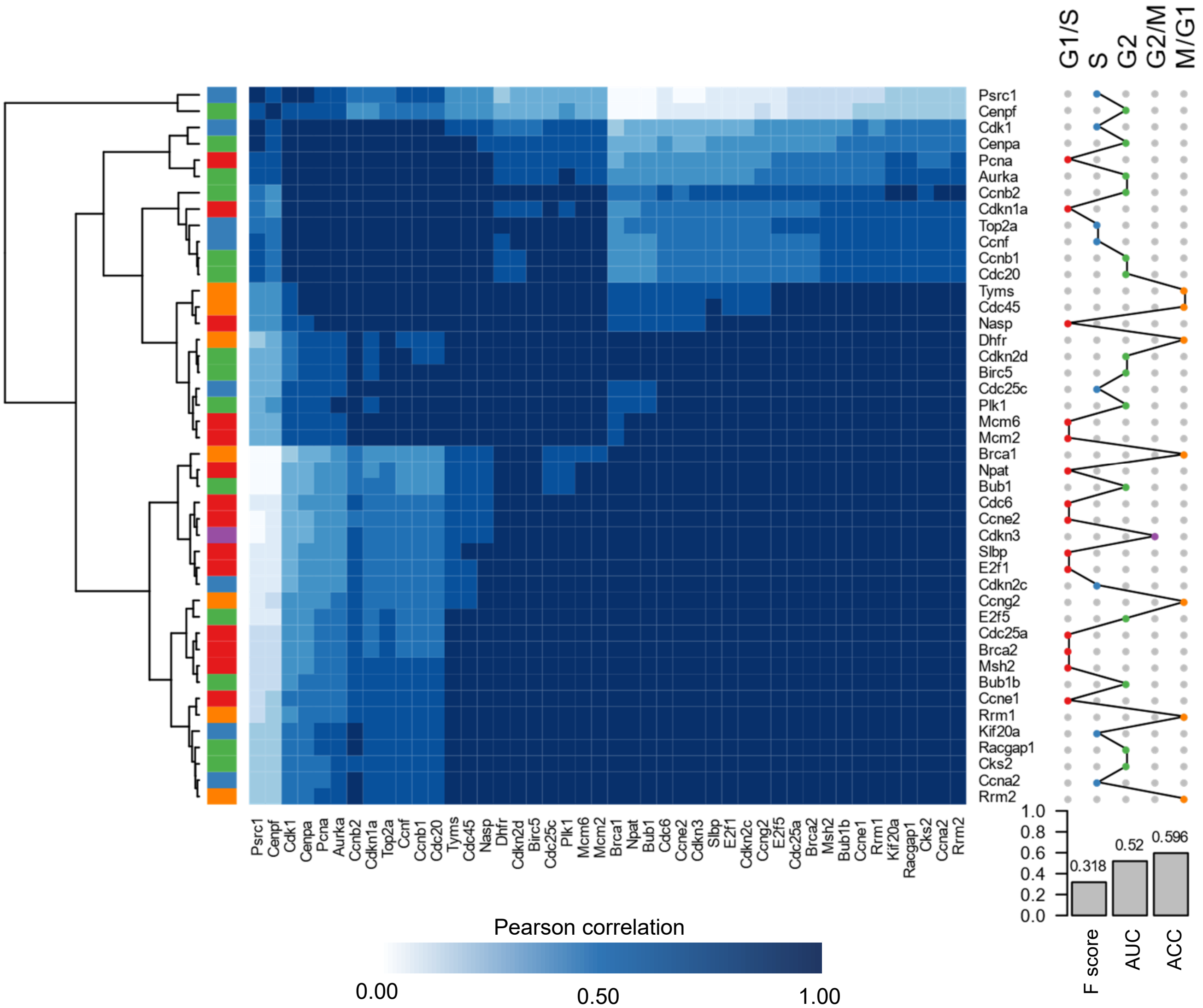

**Figure S4 (continue)**

**I. Imputed by SAVER**

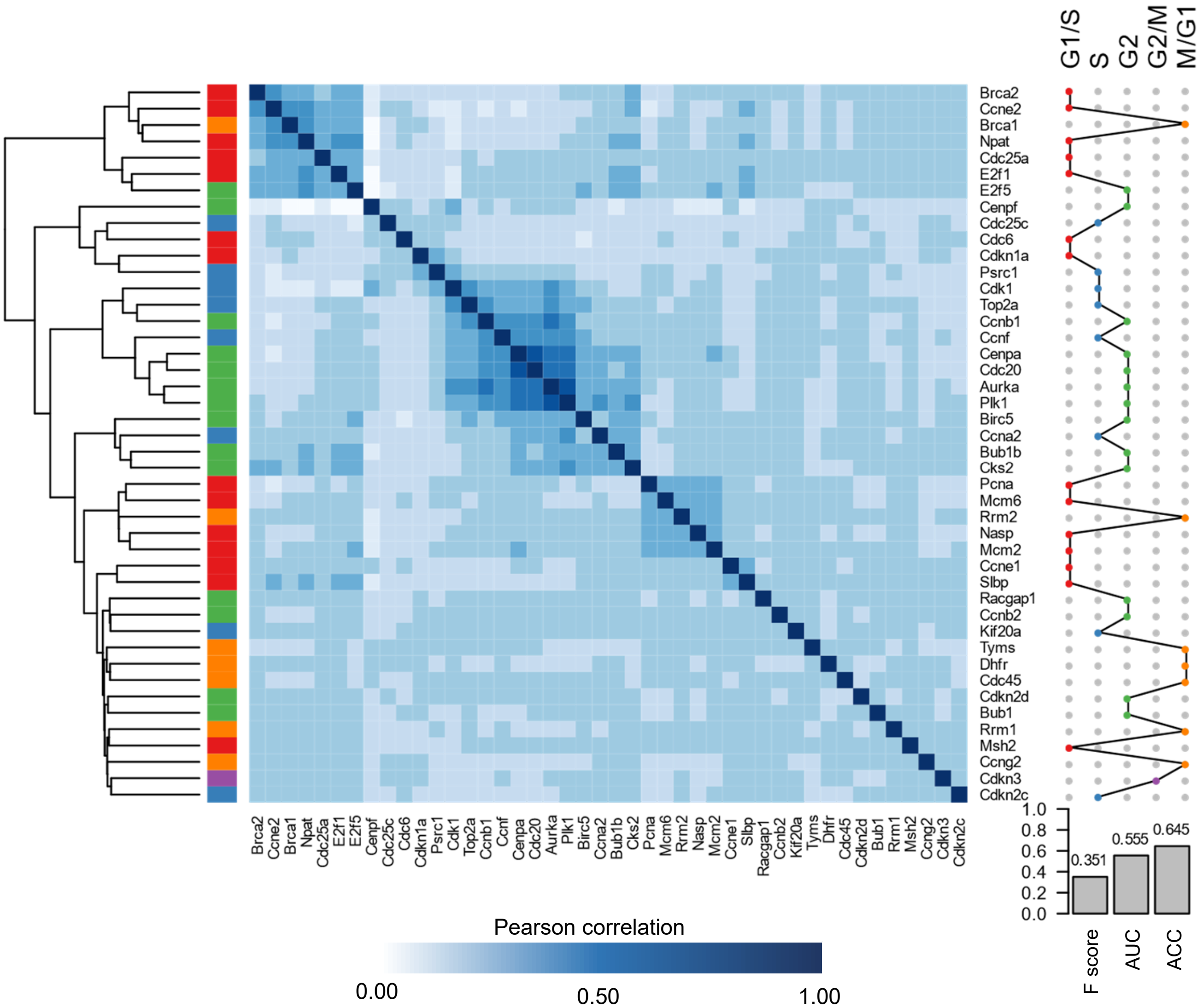

**Figure S4 (continue)**

**J. Imputed by scImpute**

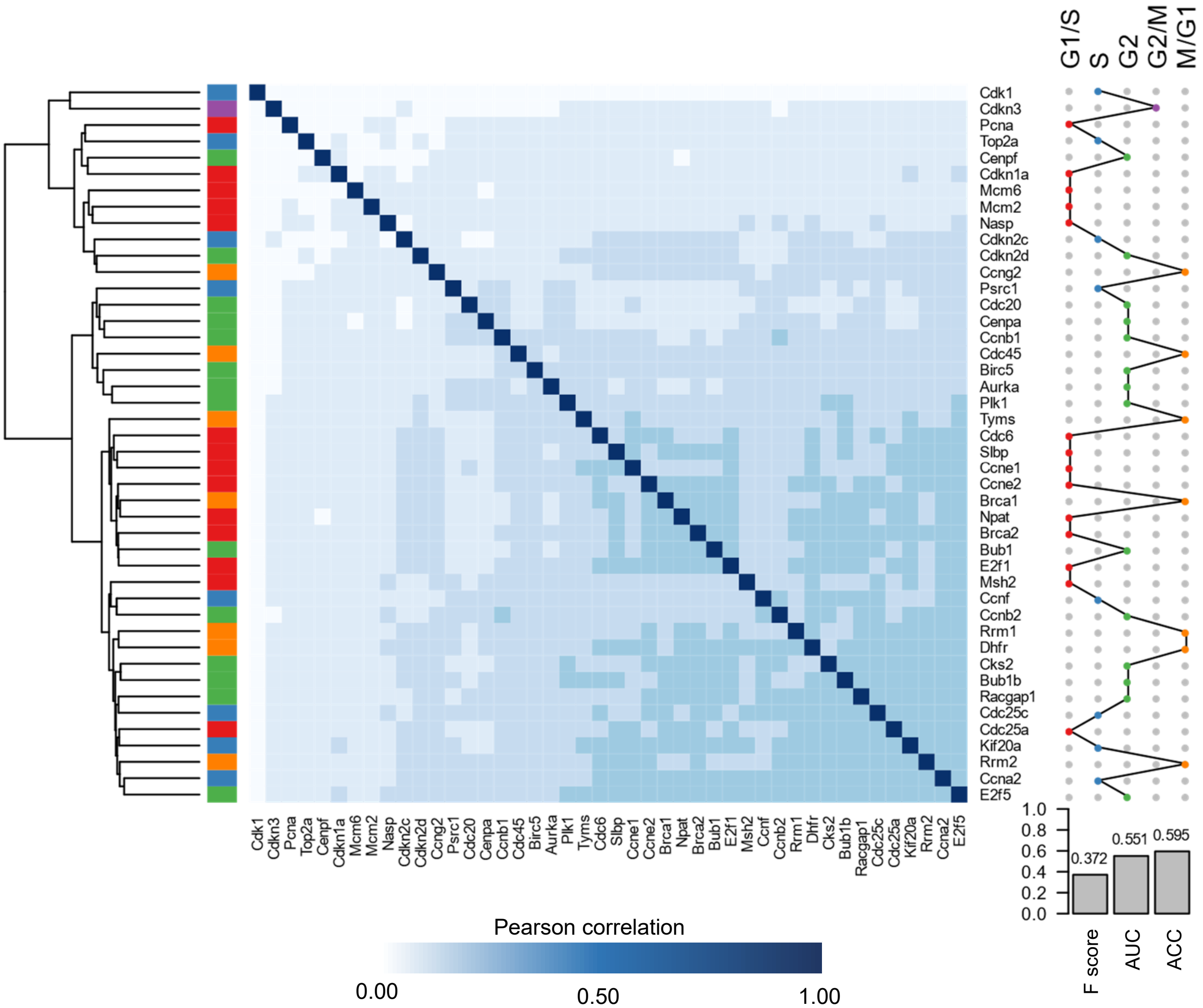

**Figure S4 (continue)**

**K. Imputed by VIPER**

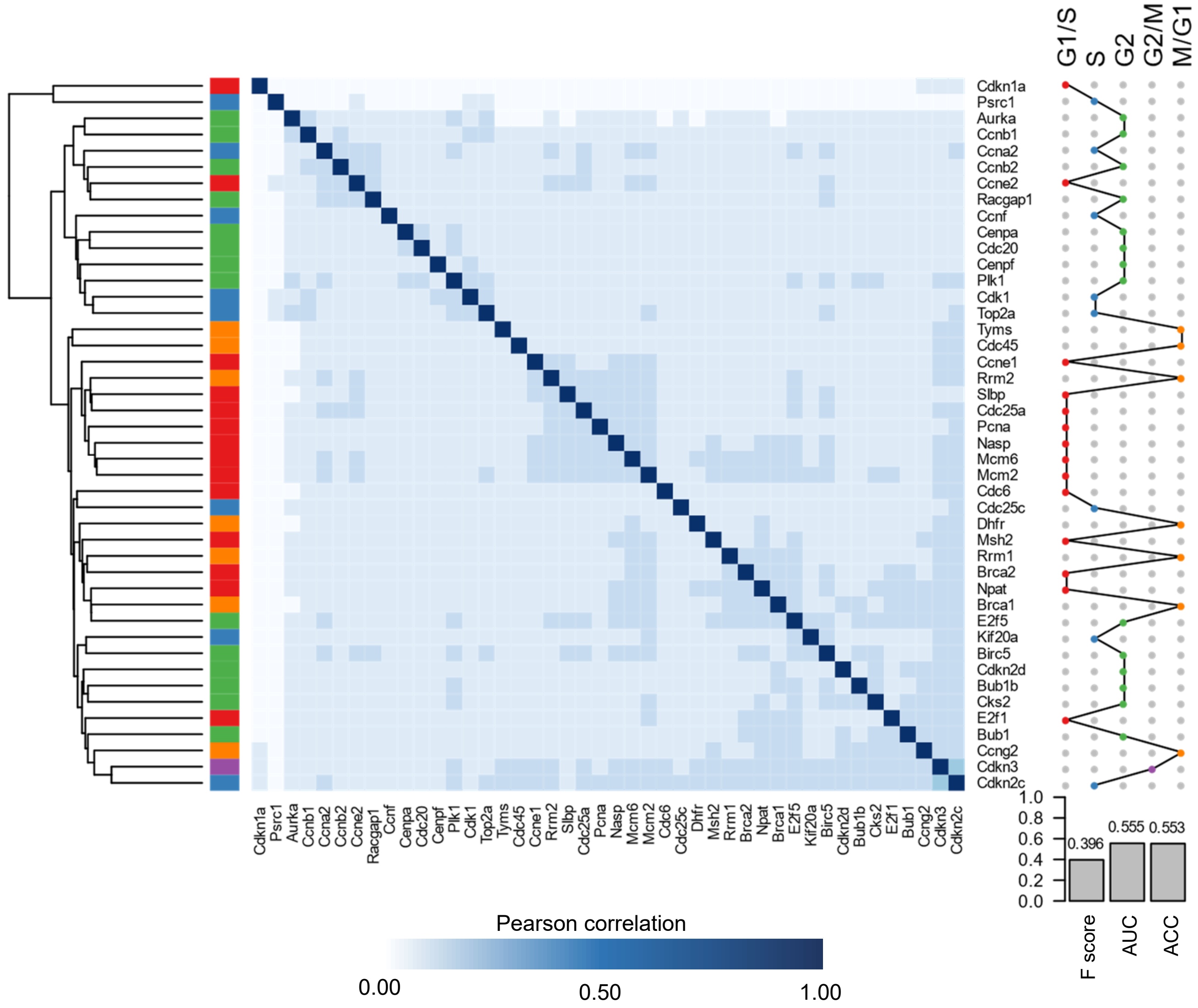

**Figure S4 (continue)**

**L. Imputed by DeepImpute**

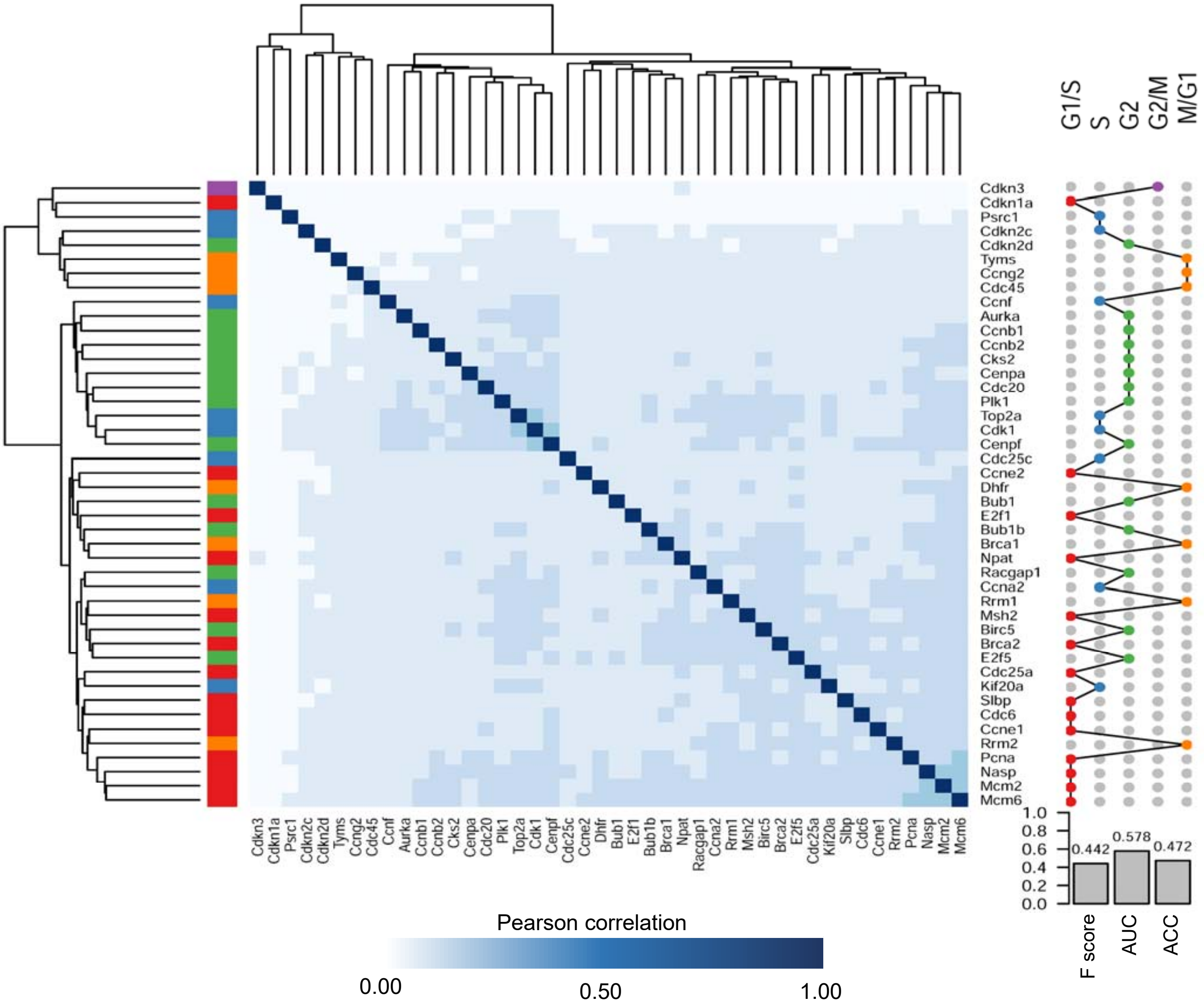

**Figure S4 (continue)**

**M. Imputed by AutoImpute**

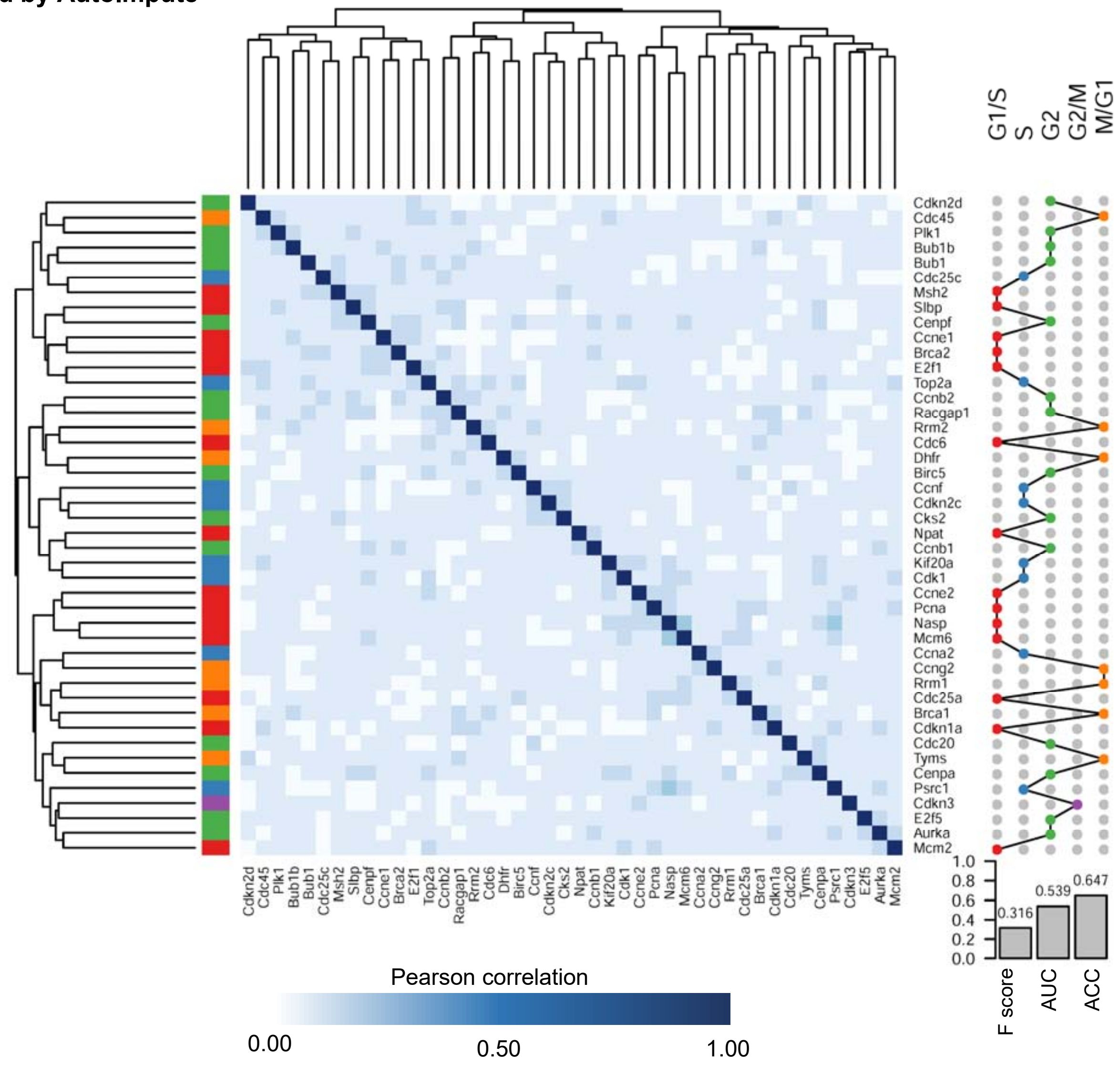

**Figure S4 (continue)**

**N. Imputed by scGAIN**

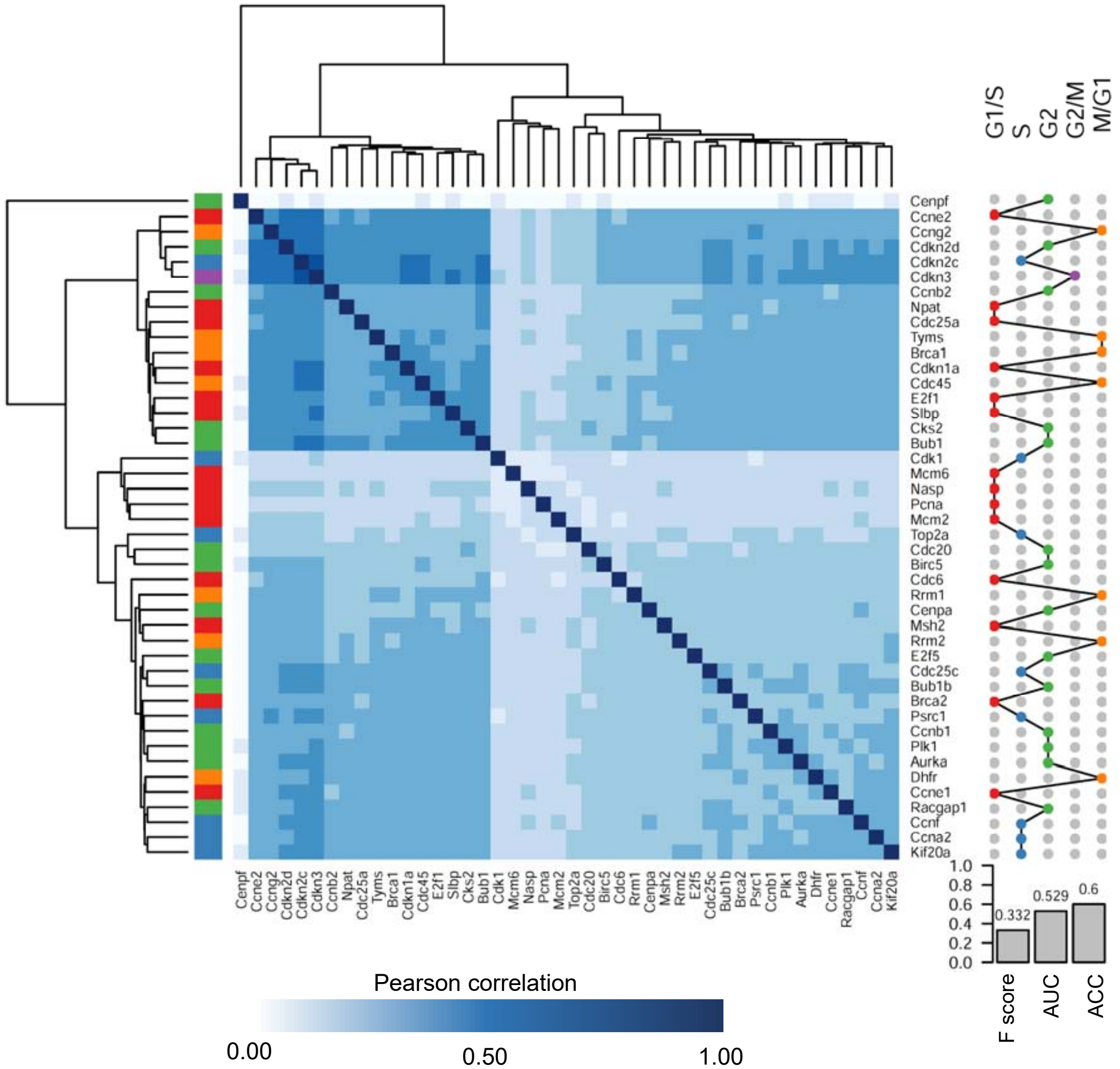

**Figure S5. Correspondence between single-cell and bulk differential expression analysis.** Scatterplots depict the estimated log fold change for each gene derived from differential analysis using bulk (x-axis) and single-cell count matrix imputed by different methods. n, the number of differentially expressed genes (p-adjust <= 0.05), red for up-regulated and blue for down-regulated genes; r, the Pearson correlation coefficient; p, the p-value of Pearson's correlation test. Red dotted line, the linear regression line. Gray dots are those genes of inconsistently differential expression in two studies.

**Related to Figure 4C.**

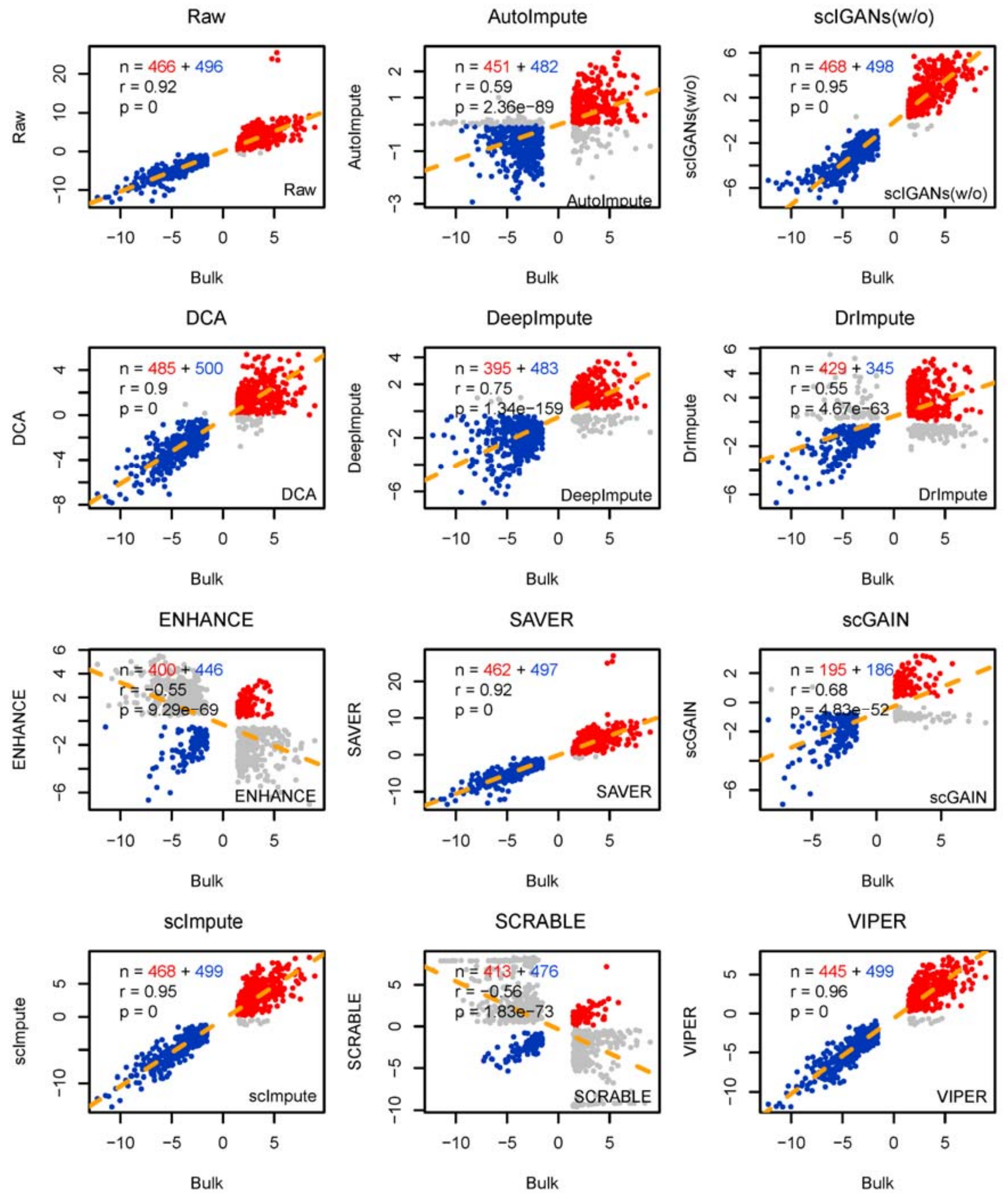

**Figure S6. Expression of five signature genes for bulk and single-cell count matrix imputed by different methods. A.** expression of signature genes of H1 cells**. B.** expression of signature genes of DEC cells.

**Related to Figure 4D.**

**A signature genes of H1 cells.**

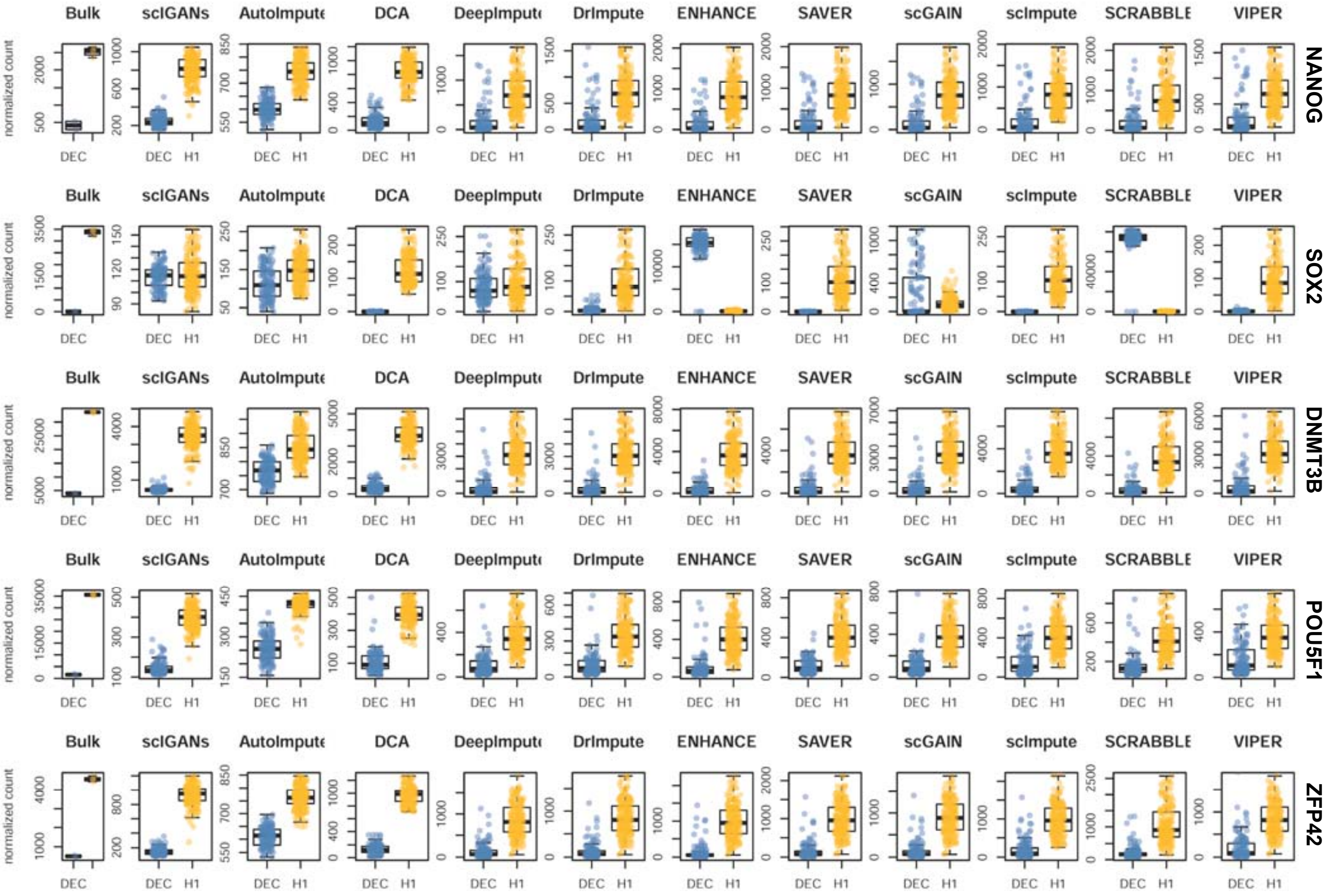

**Figure S6 (continue)**

**Related to Figure 4D.**

**B signature genes of DEC cells.**

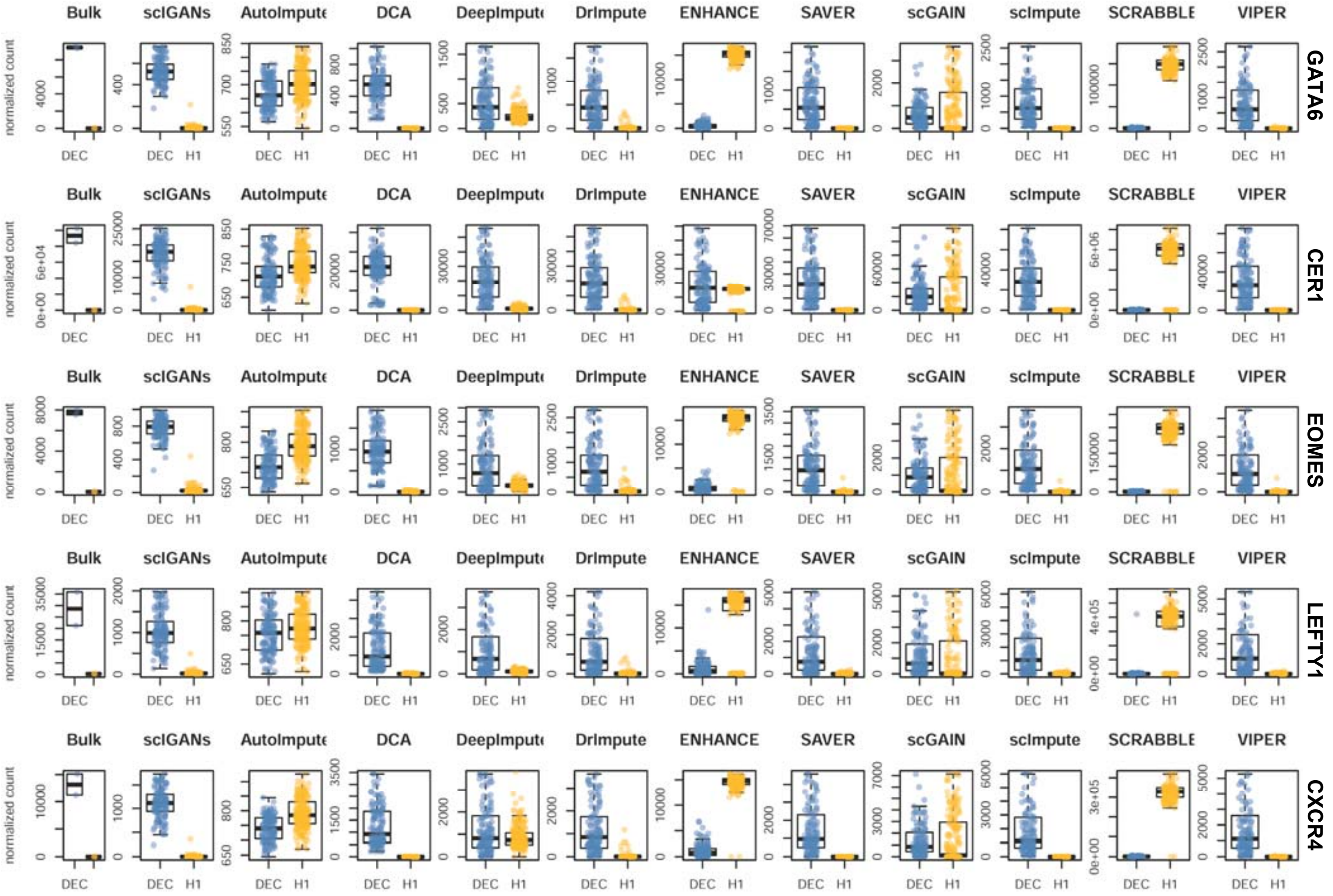

**Figure S7. Expression of signature genes across single cells. A-L.** The UMAP plots of the single cells overlaid by the expression of SOX2 and CECR4, which is the marker gene of H1 and DEC cells, respectively. Plots are for the other 11 imputation methods.

**Related to Figure 4E-F.**

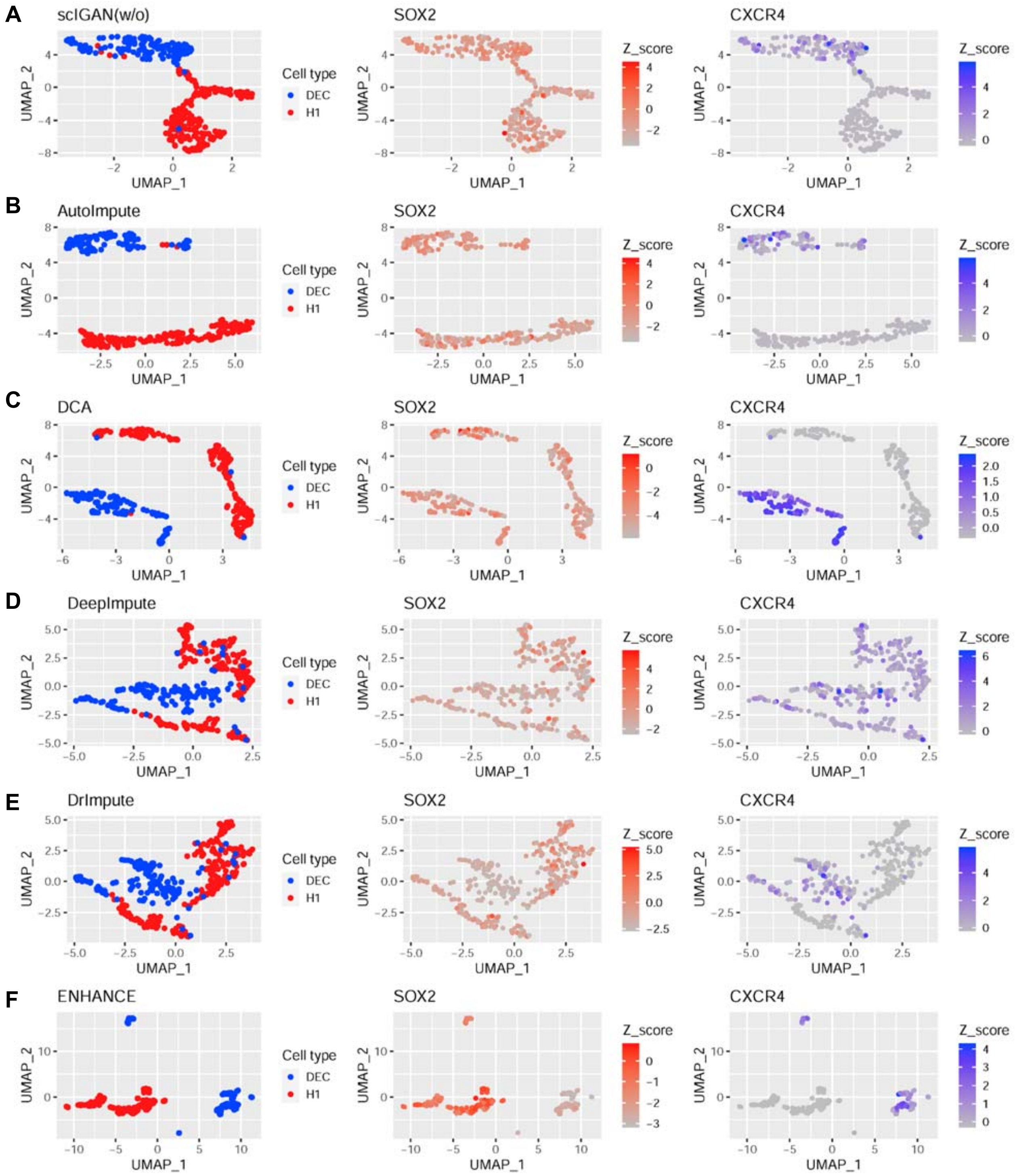

**Figure S7 (continue)**

**Related to Figure 4D-E.**

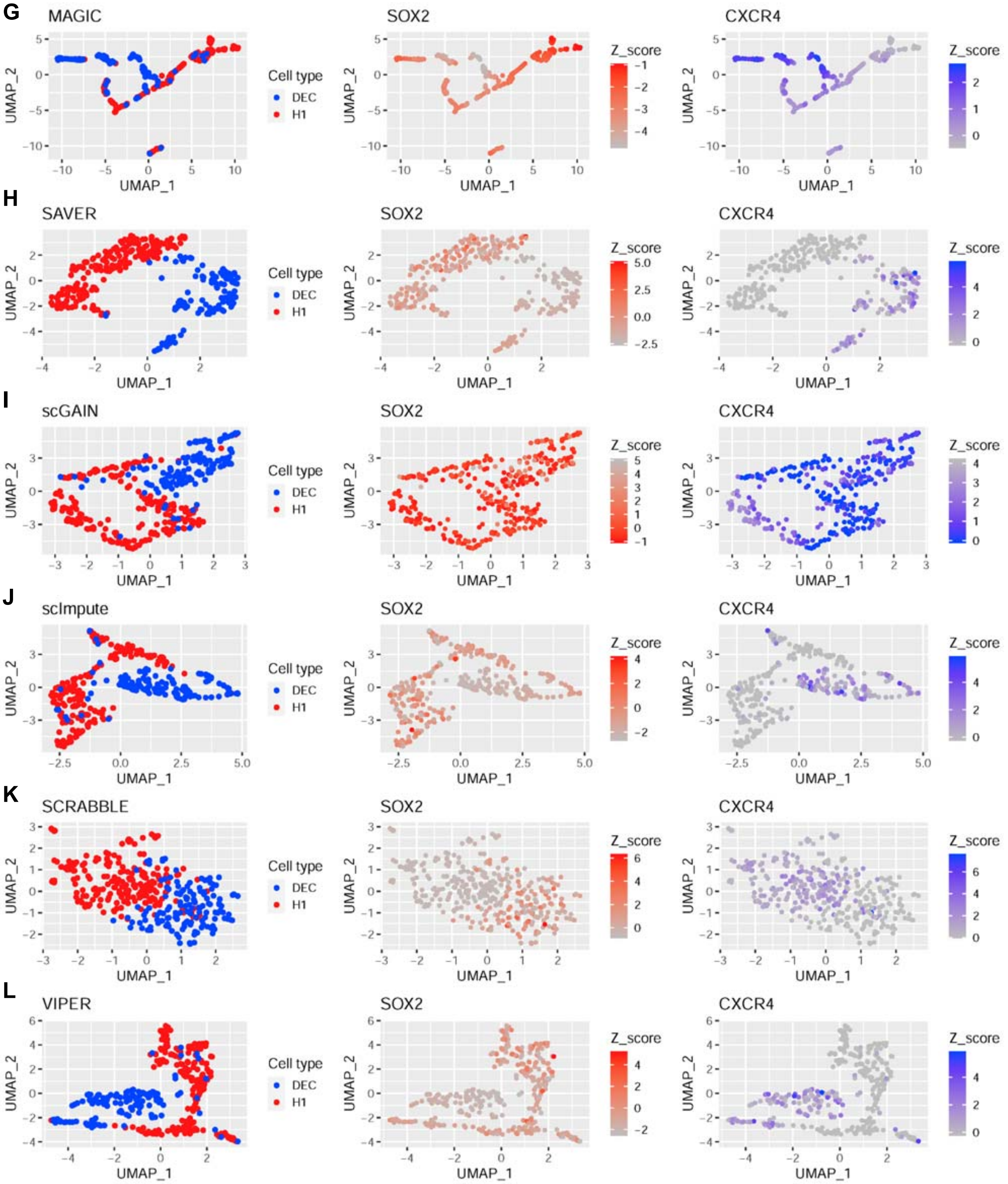

**Figure S8. Trajectories and expression dynamics of signature genes during H1 cell differentiation. A-L.** The trajectories reconstructed by monocle3 from scRNA-seq data imputed by the other 11 methods. The right two panels show the expression dynamics of signature genes of H1 cells (middle) and DEC cells (right) along with the pseudotime.

**Related to Figure 5.**

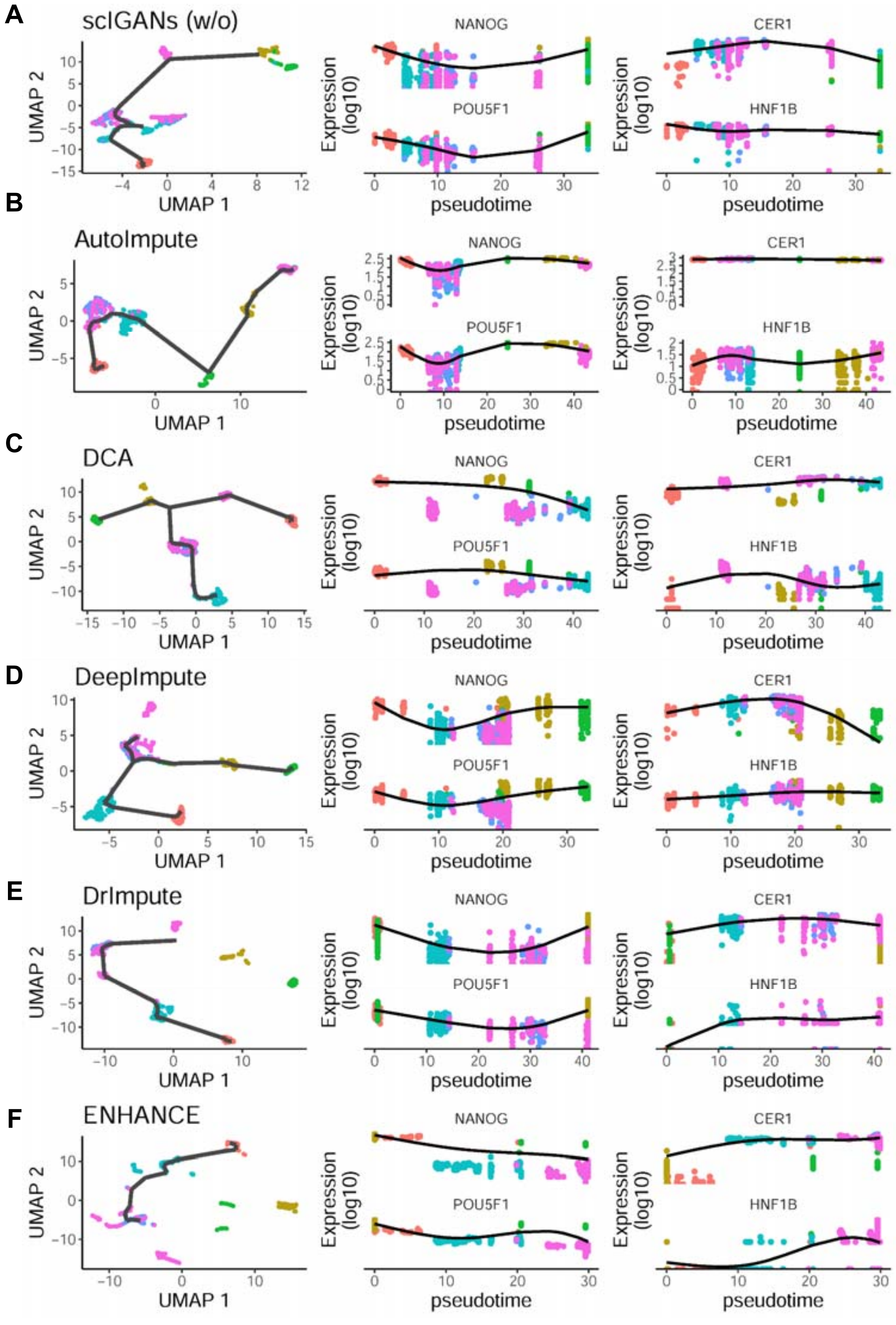

**Figure S8 (continue)**

**Related to Figure 5.**

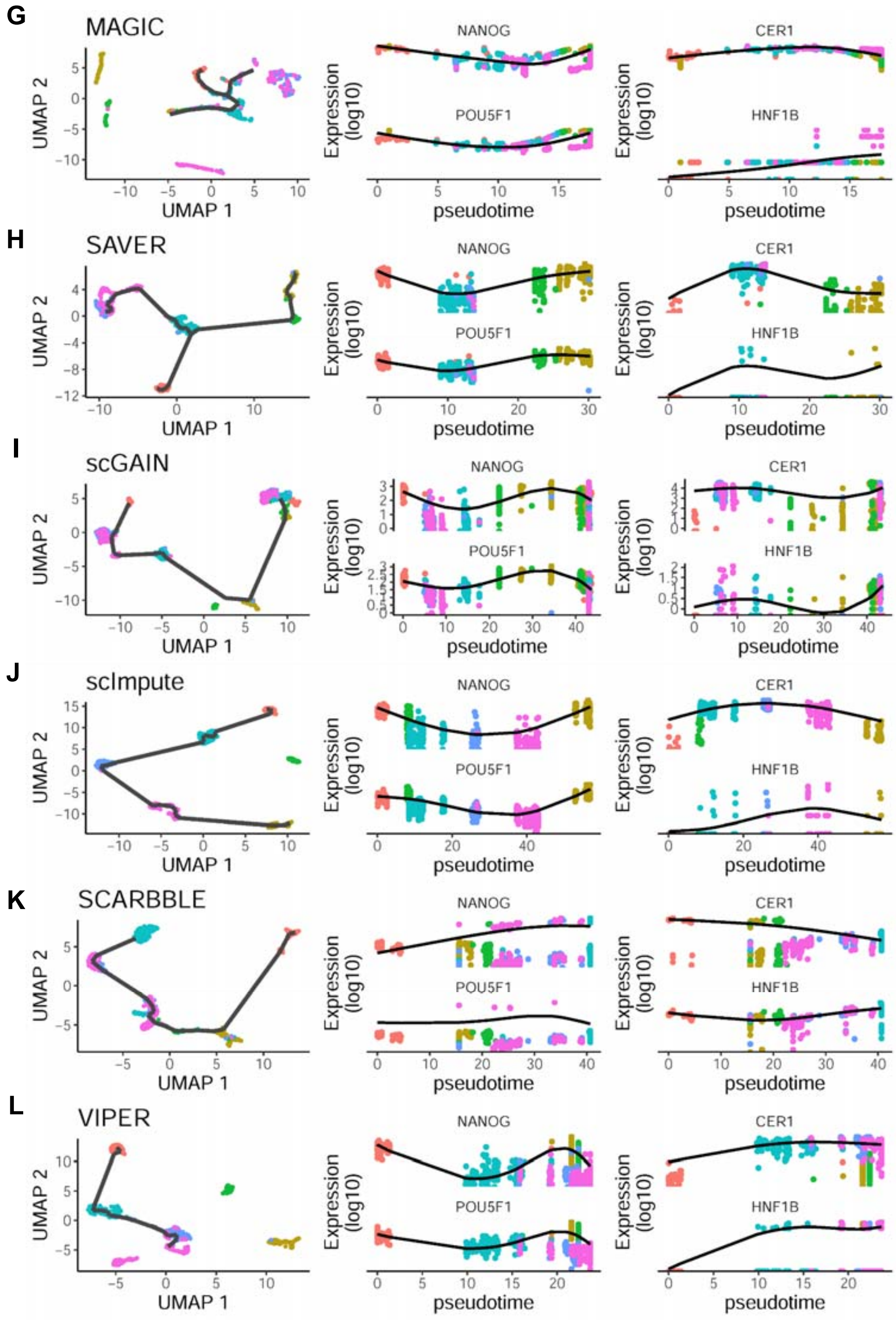

**Figure S9. The performance of imputation methods on cell-type identification using a small set of genes with very low expression or cross-cell variance. A-H.** The same series of plots as **Figure 6** for the other 11 tested methods and scIGANs trained without labels. Random, randomly picked 1024 genes from all expressed genes; mean.low, randomly picked 1024 genes from 5000 genes with lowest expression means; mean.top, randomly picked 1024 genes from 5000 genes with highest expression means; sd.low, randomly picked 1024 genes from 5000 genes with lowest expression variance (standard deviation) across all cells; sd.top, randomly picked 1024 genes from 5000 genes with highest expression variance across all cells.

**Related to Figure 6.**

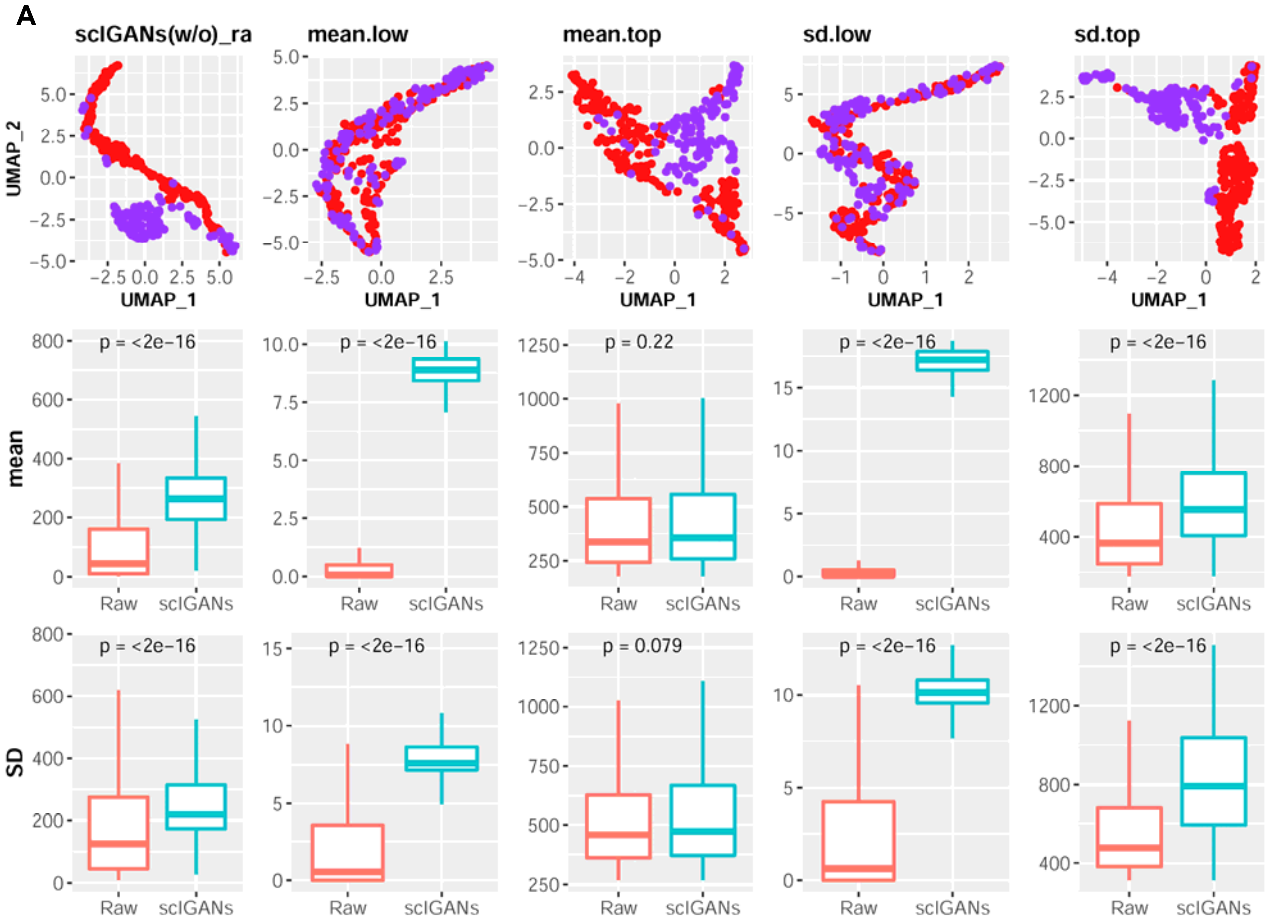

**Figure S9 (continue)**

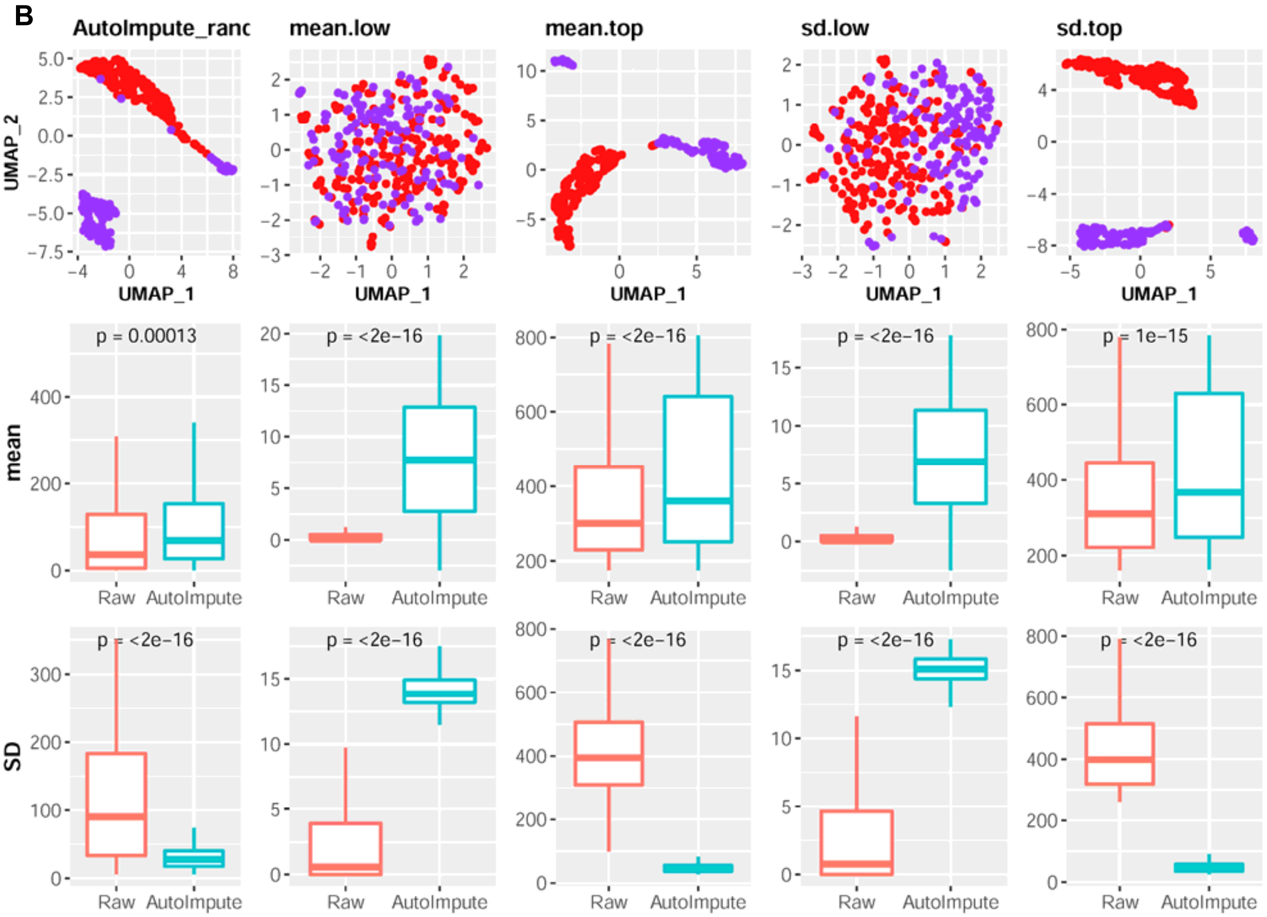

**Figure S9 (continue)**

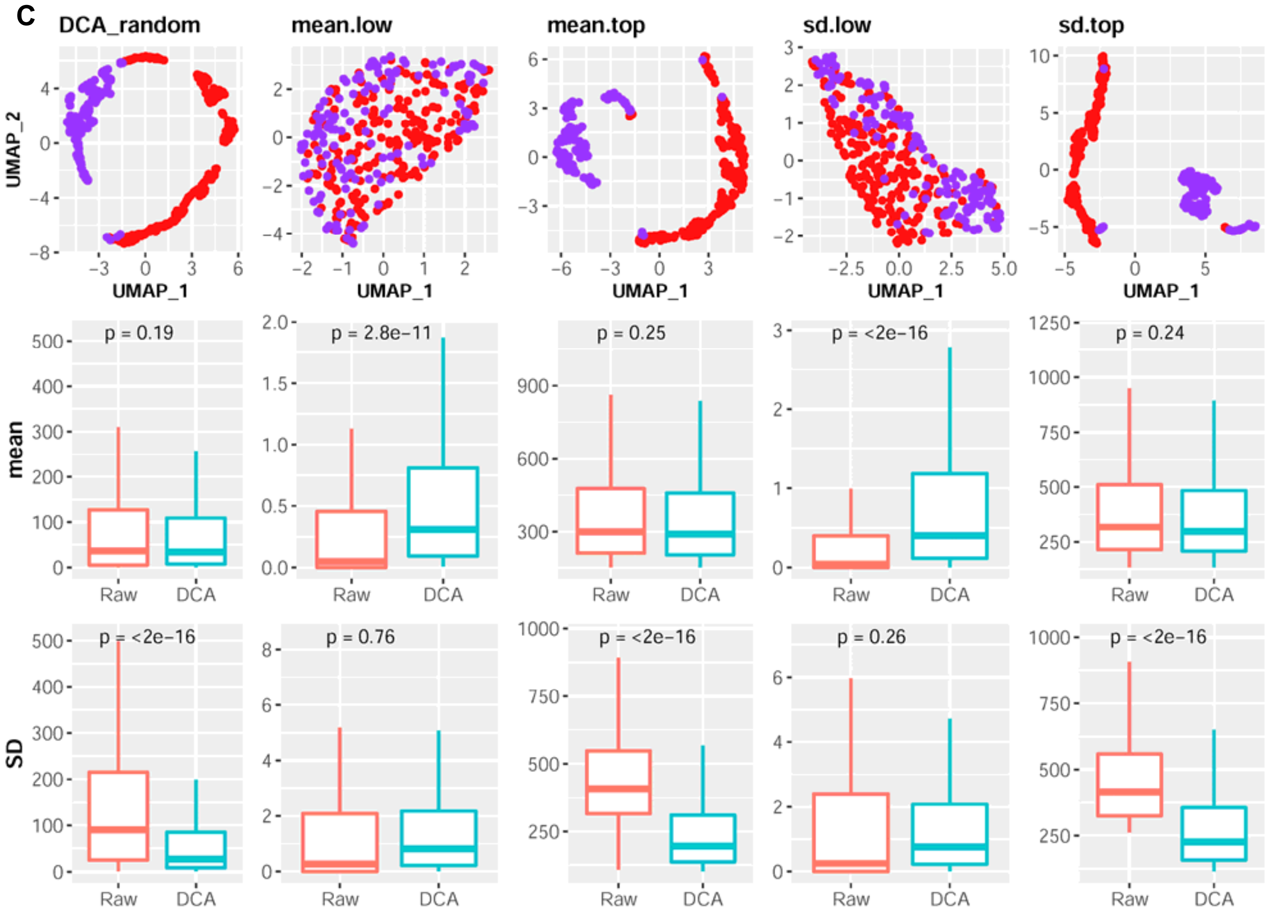

**Figure S9 (continue)**

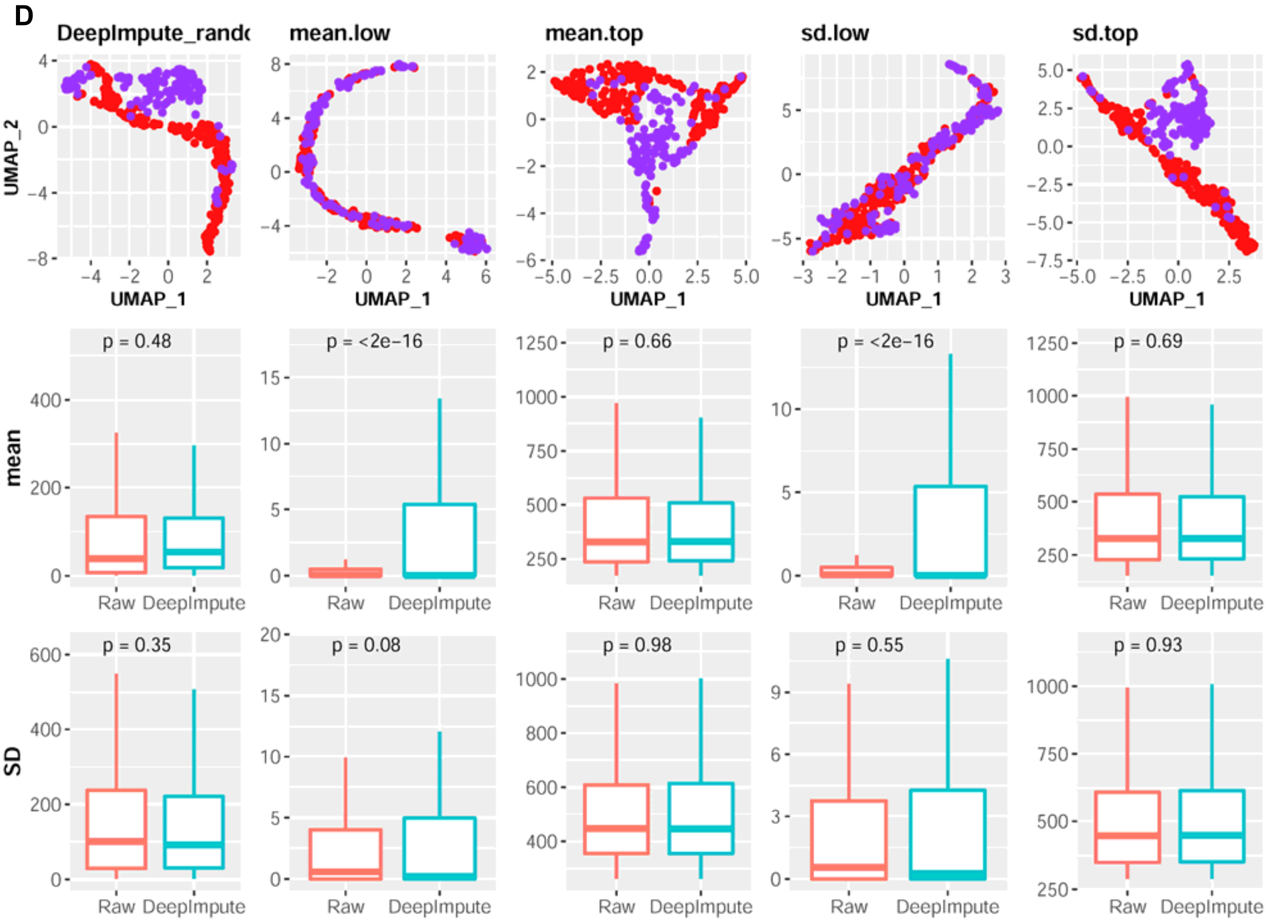

**Figure S9 (continue)**

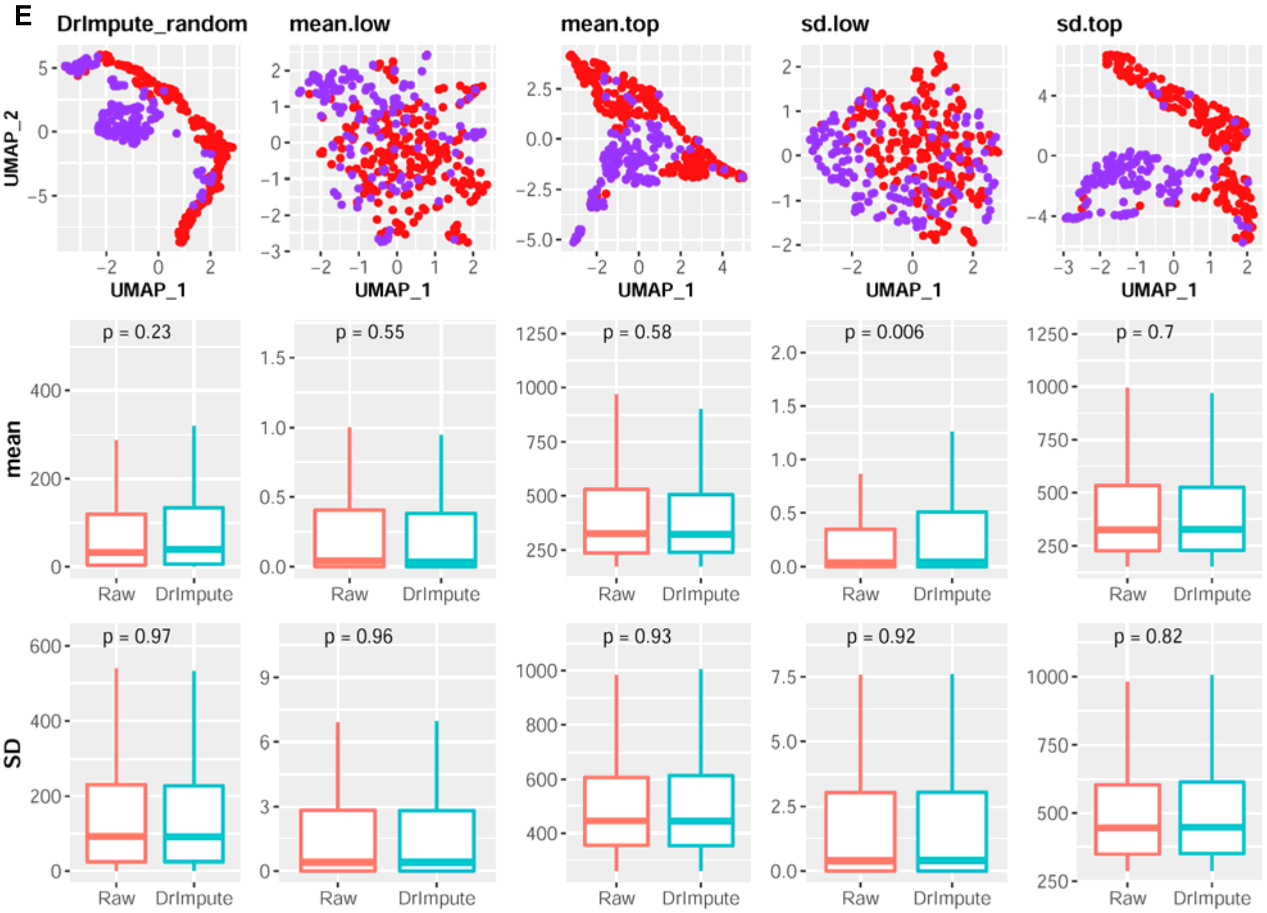

**Figure S9 (continue)**

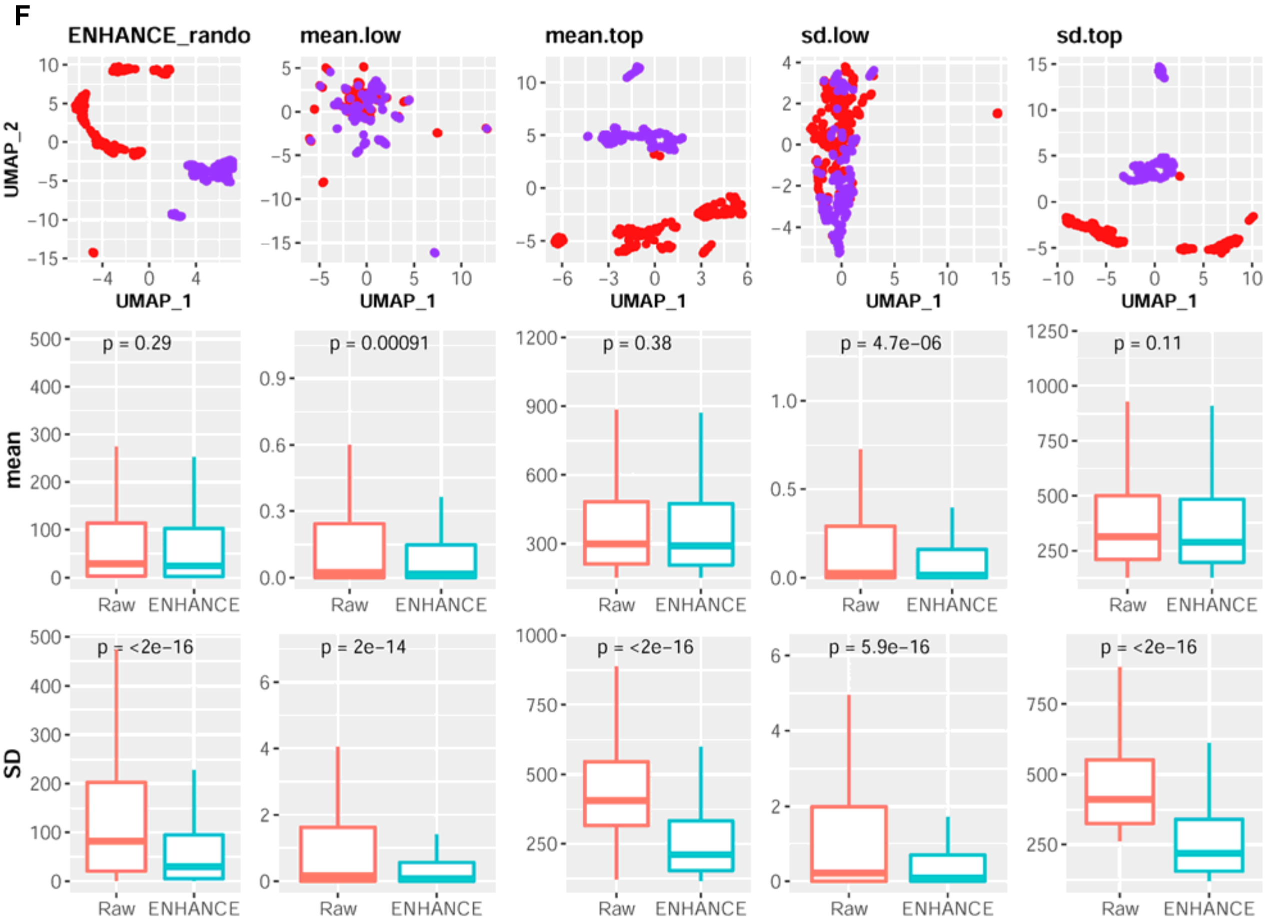

**Figure S9 (continue)**

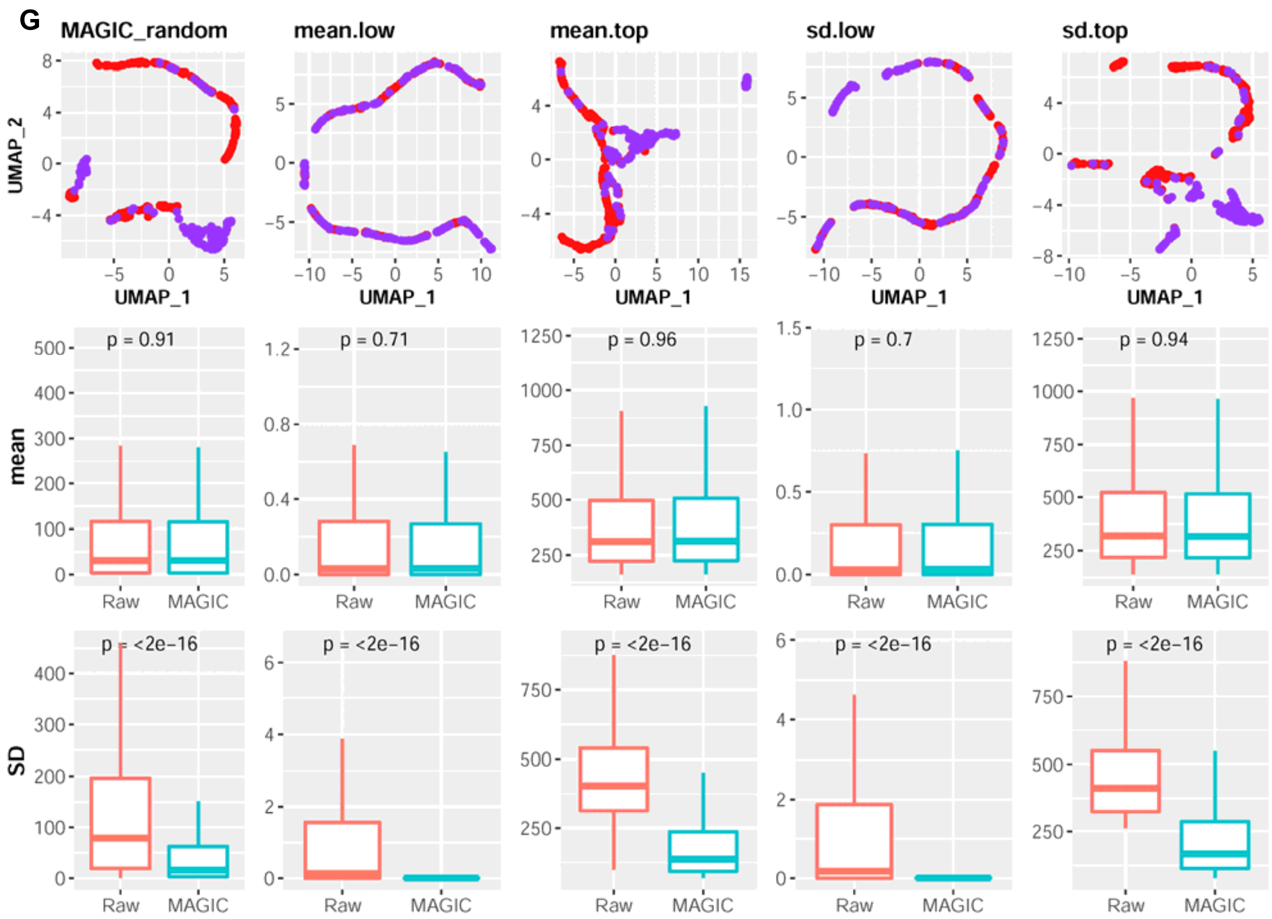

**Figure S9 (continue)**

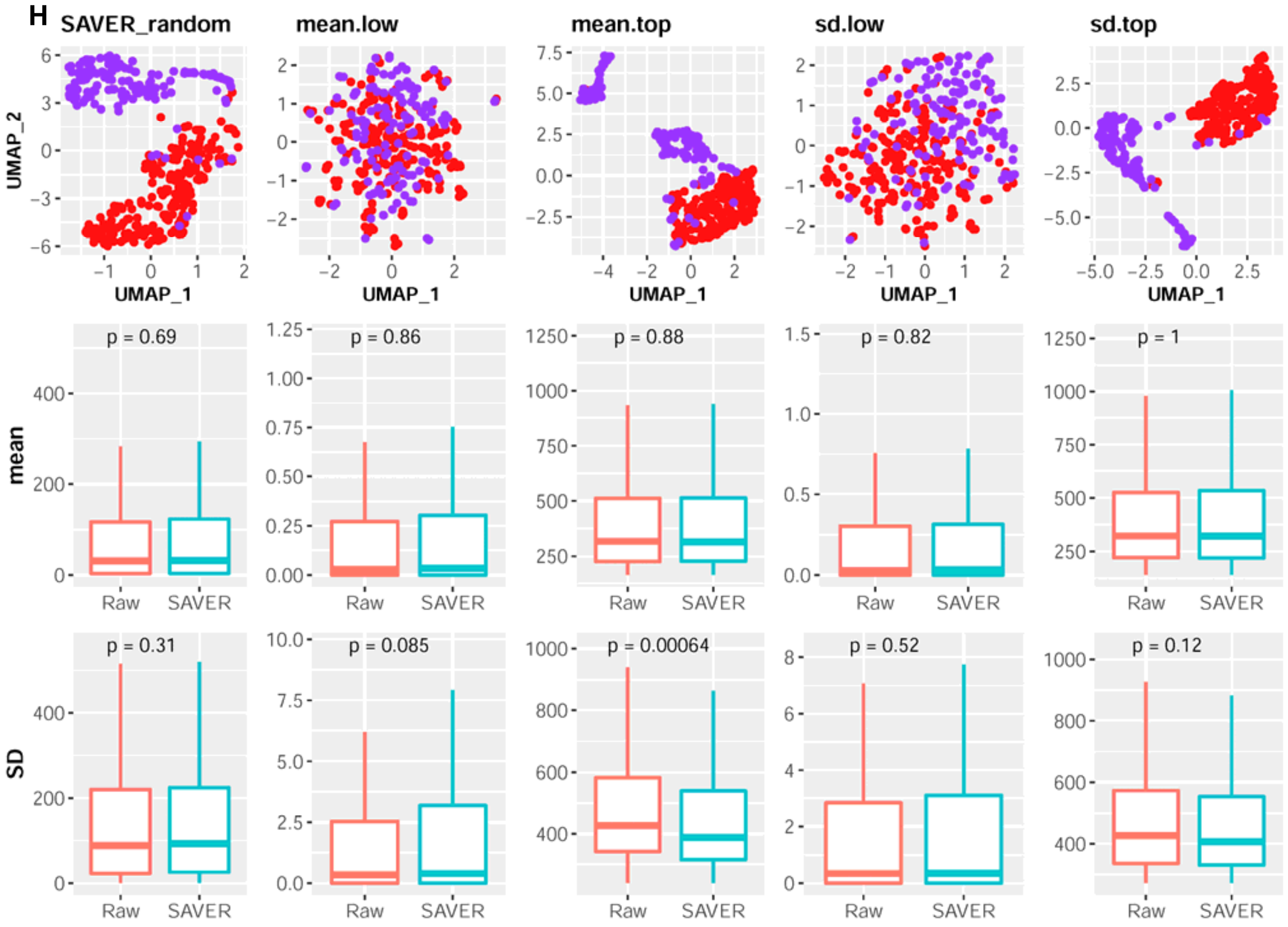

**Figure S9 (continue)**

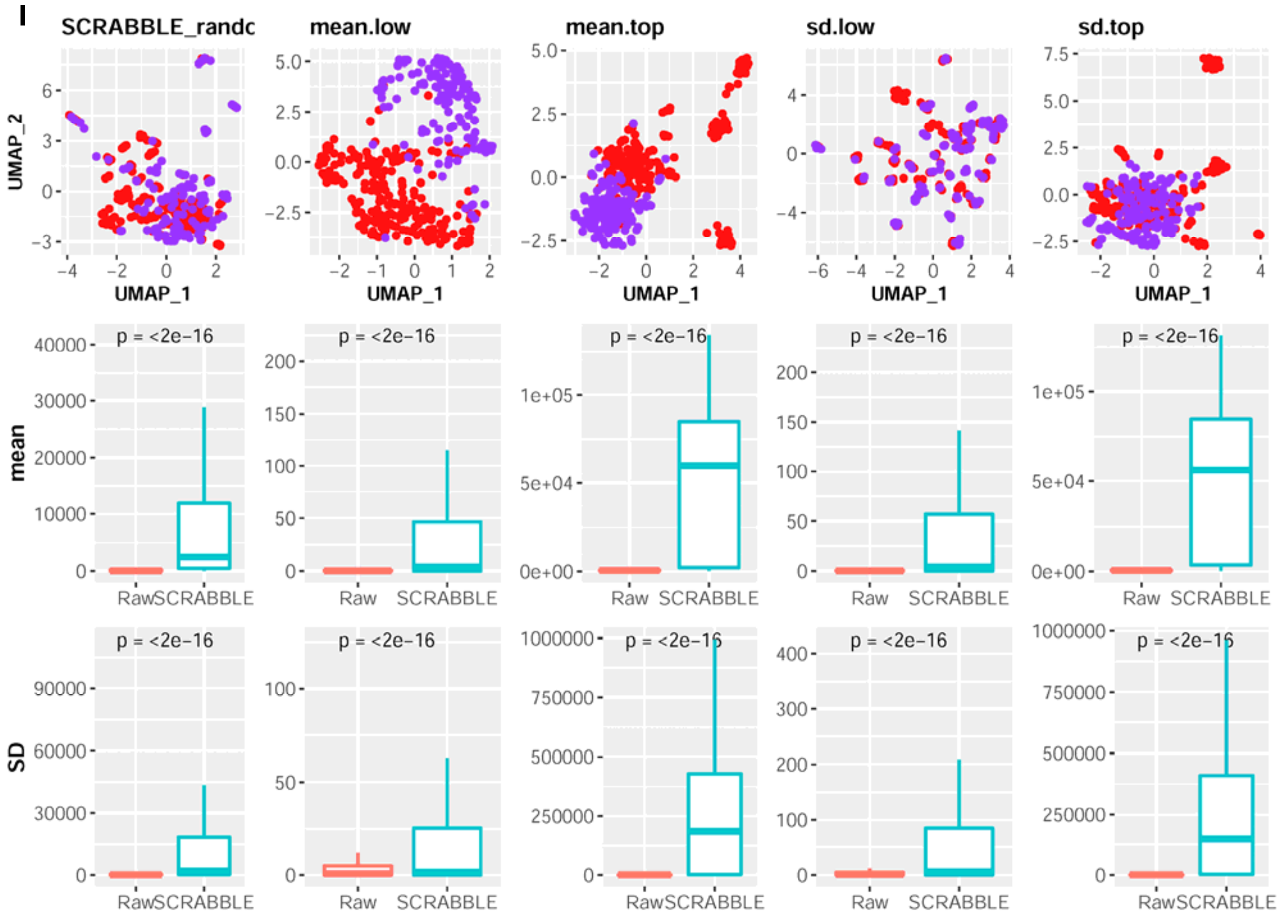

**Figure S9 (continue)**

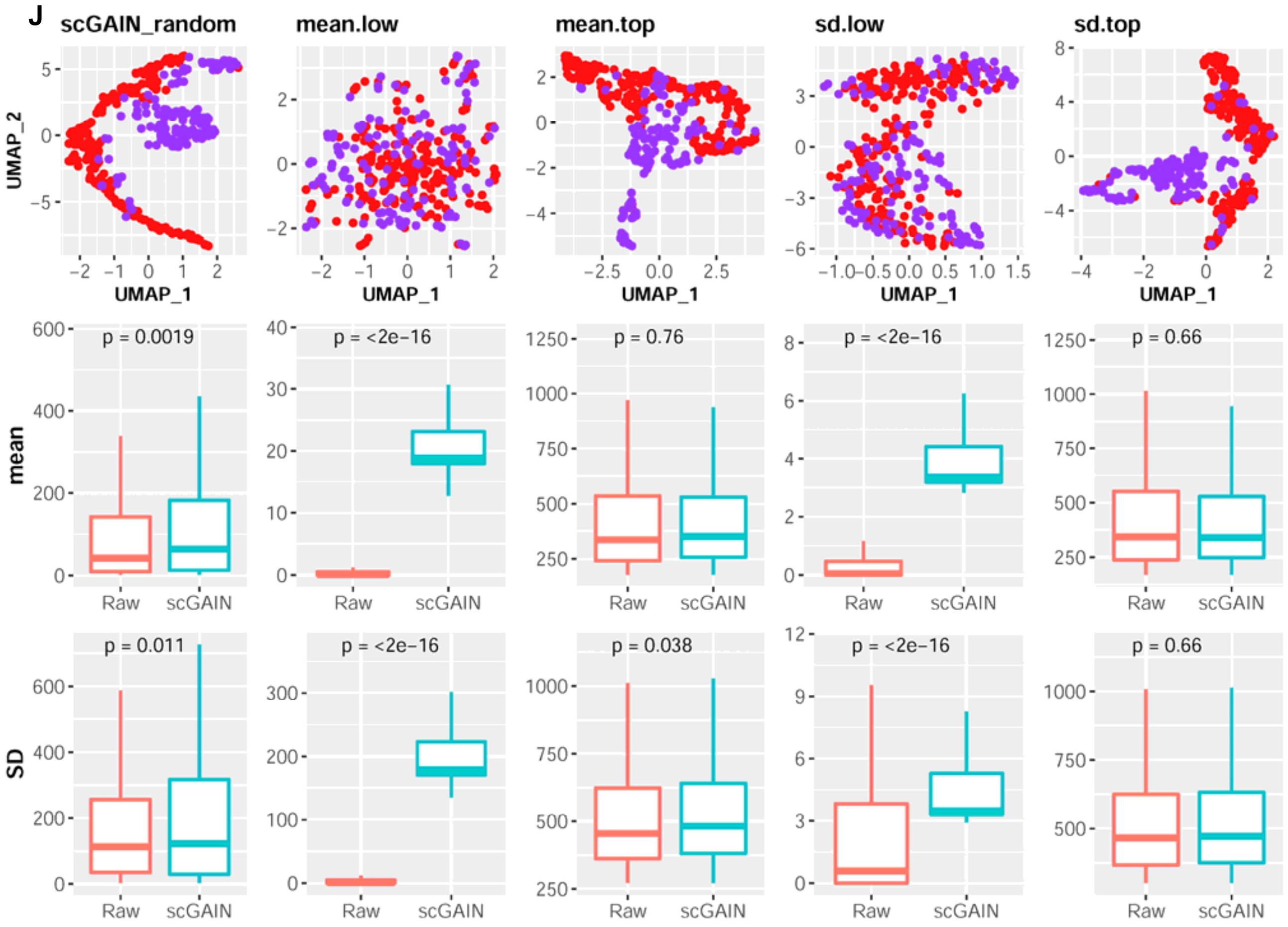

**Figure S9 (continue)**

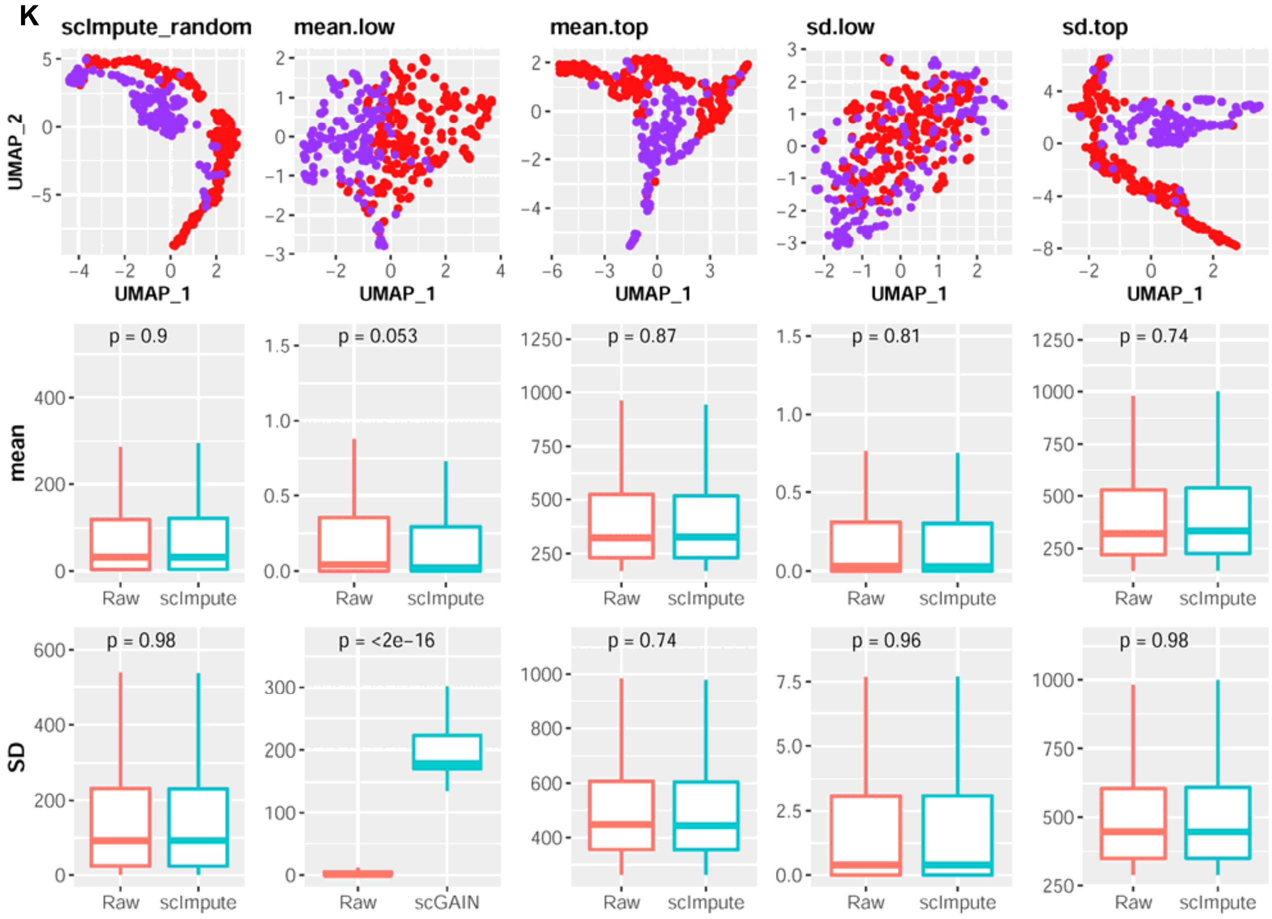

**Figure S9 (continue)**

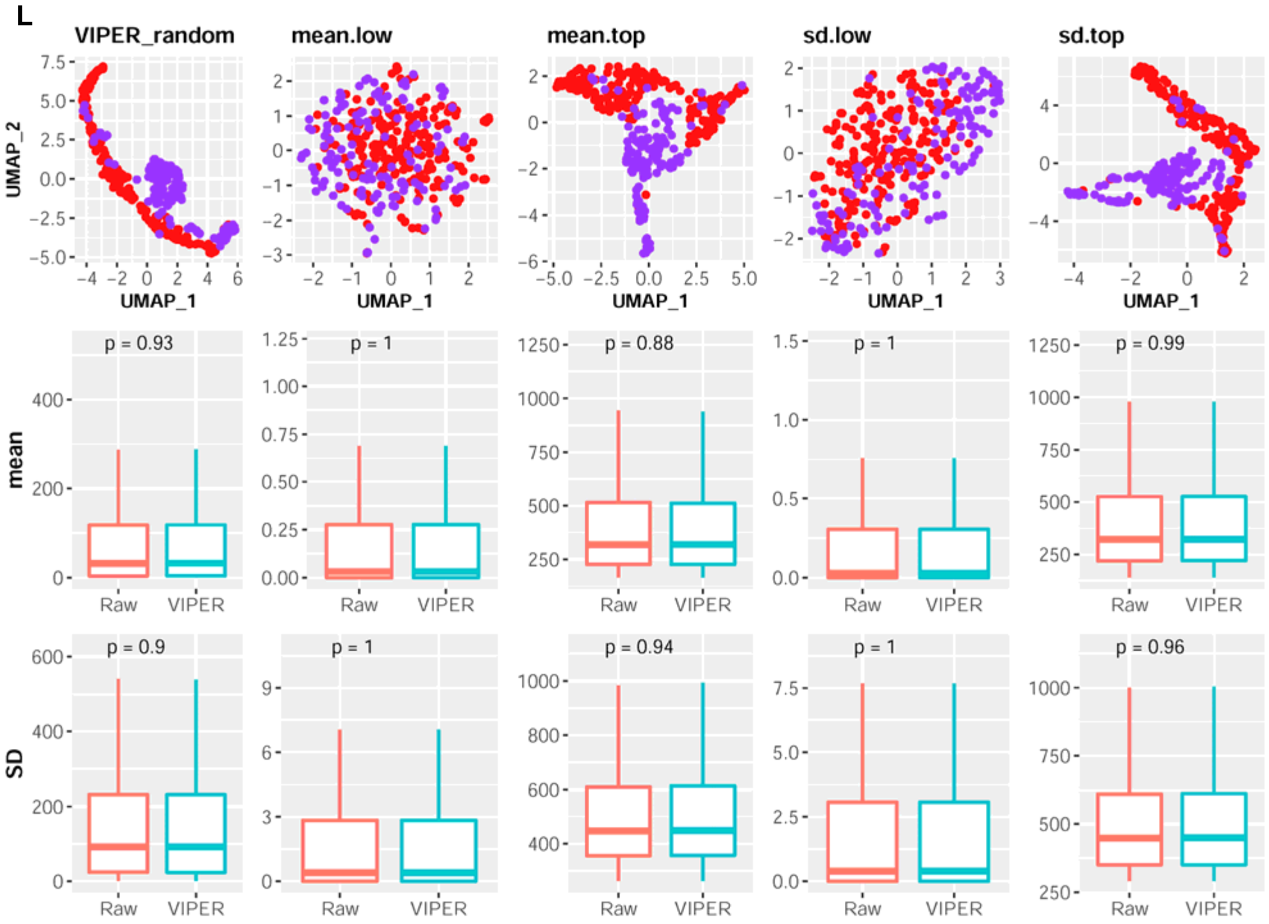

**Figure S10. The UMAP visualizations of the same cell populations sequenced by three different scRNA-seq methods.** The extended plots of **Figure 7A**. 10x, 10X Genomics/Chromium; celseq2, CEL-seq2/Fluidigm; dropseq, Drop-seq/Droplet.

**Related to Figure 7A.**

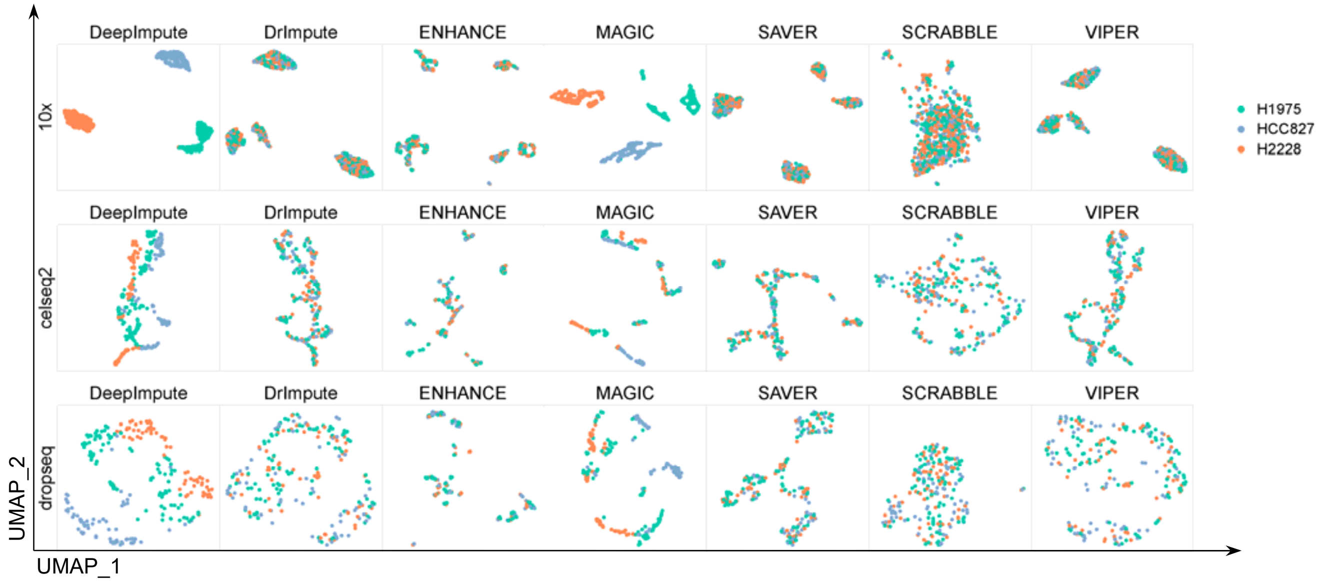

**Figure S11. Cross-platform performances of different methods.** The extended plots of **Figure 7B**, shown as the other six clustering metrics. The source data for all clustering metrics are provided in **Supplementary Table S8**.

**Related to Figure 7B.**

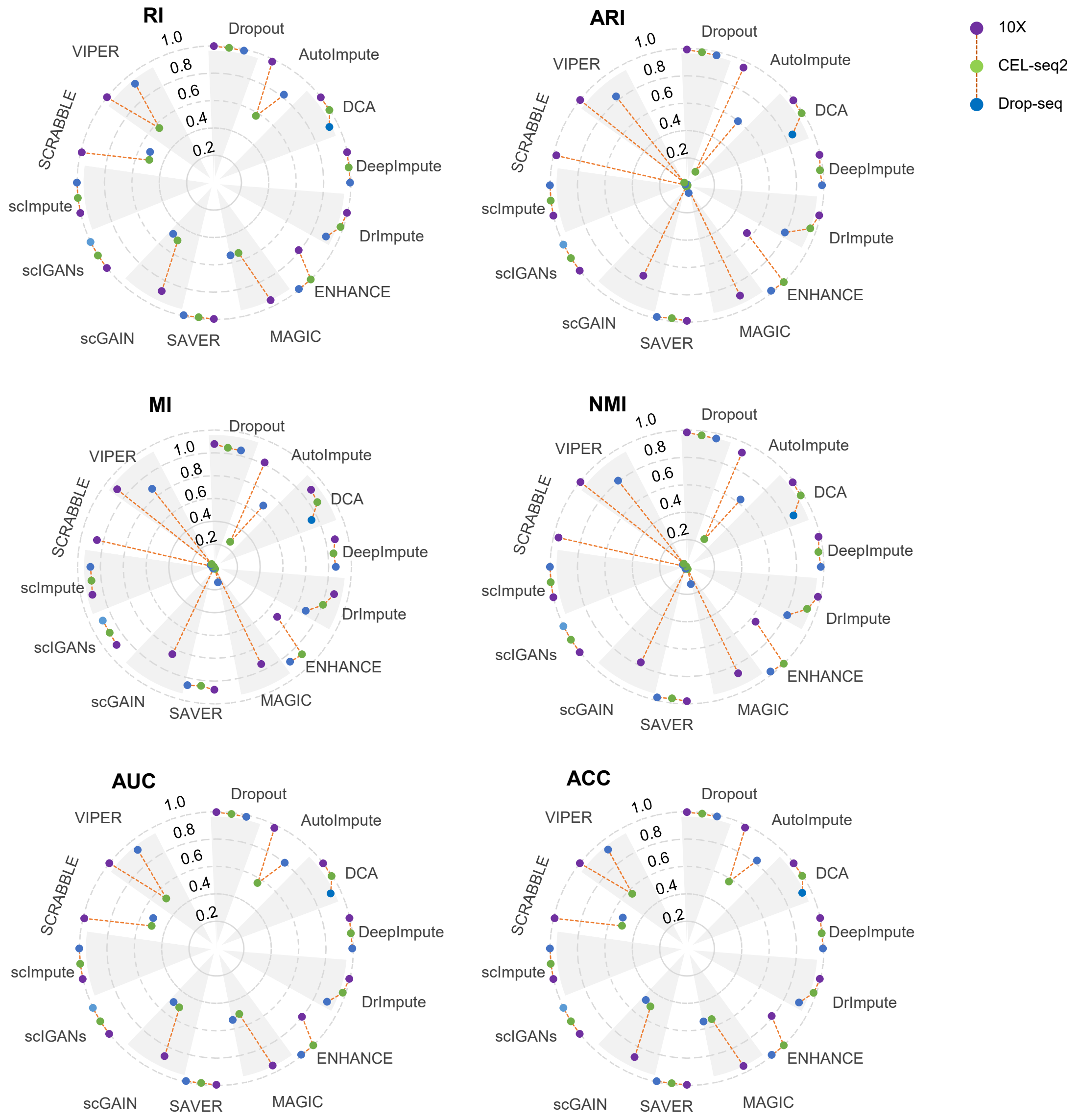

**Figure S12. scIGANs is scalable to scRNA-seq methods and data sizes. A.** The cell-cell similarity of the expression profiles from different scRNA-seq methods represented as the Spearman correlation coefficients from the expression profiles of the same cell type generated by different sequencing methods. **B.** The average memory usage (in GB) of the imputation methods on datasets with different sizes (cell numbers). The failed jobs in **Supplementary Table S9** are assigned auxiliary values of 65 GB for plotting purposes.

**Related to Figure 7C-D.**

A

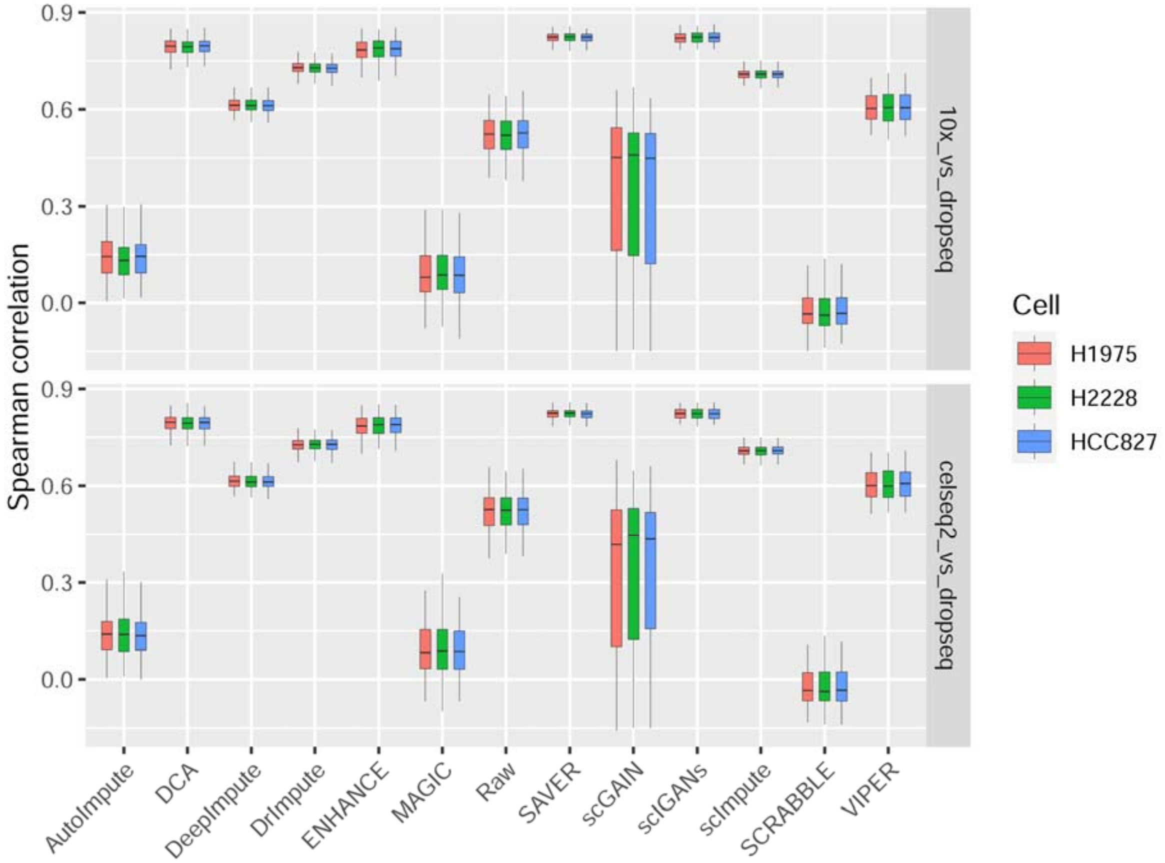


B

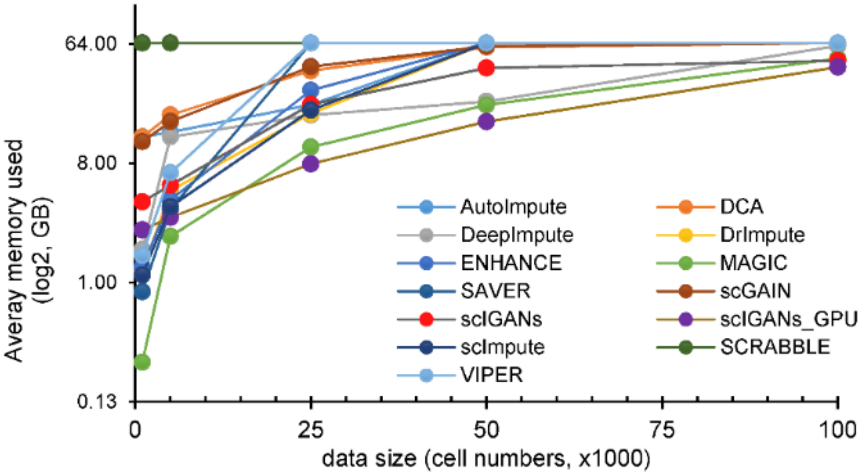
