## Additional file 2: Tables S1-S3, and Tables S5-S9. for "scIGANs: single-cell RNA-seq imputation using generative adversarial networks"

### Table S1. The summary of the simulated and real scRNA-seq datasets used in this work.

| Method sections | Dataset name | Protocol | Platform | Count type | Source | Dataset details | Result sections | Derived Figures and Tables |
|---|---|---|---|---|---|---|---|---|
| Simulated datasets | CIDR_sim | Simulated with CIDR | NA | short reads | Simulation | 150 cells × 20180 genes; 3 cell types | Section 2 | Figures 2A, S2A and Table S2 |
| | Splatter_sim | Simulated with Splatter | NA | short reads | Simulation | 1000 cells × 800 genes; 3 cell types; three datasets of different dropout rates (71%, 83%, and 87%) | | Figures S2B-E and Table S3 |
| | Splatter_sim100 | Simulated with Splatter | NA | short reads | Simulation | 1000 cells × 800 genes; 3 cell types; three datasets of different dropout rates (71%, 83%, and 87%) with 100 replicate for each | | Figures 2B, S3A-F; Table S4 |
| Real scRNA-seq datasets | Human_brain | SMARTer/ Fluidigm | plate-based | short reads | GEO: GSE67835 | 420 cells × 22085 genes; 8 cell types | | Figures 2C-D and S3G; Table S5 |
| | ERCC | SMARTer/ Fluidigm | plate-based | short reads | ArrayExpress: E-MTAB-2805 | 289 cells × 92 spike-in RNAs; 3 cell-cycle phases: G1, S, G2M | | Figures 2E and S3H-I; Table S6 |
| | Cell_cycle_phase | SMARTer/ Fluidigm | plate-based | short reads | ArrayExpress: E-MTAB-2805 | 288 cells × 38293 transcripts; 3 cell-cycle phases: G1, S, G2M | Section 3 | Figures 3A-B and S3; Table S7 |
| | Mouse_ESC | CEL-seq/ Droplet | droplet-based | UMI | GEO: GSE65525 | 6885 cells × 24175 genes; 1 cell type with 3 replicates | | Figures 3C and S4C-N |
| | Human_ESC | SMARTer/ Fluidigm | plate-based | short reads | GEO: GSE75748 | scRNA-seq: 350 cells × 19097 genes; 2 cell types: H1 ESC and DEC Bulk RNA-seq: 4 replicates for H1 ESC and 2 replicates for DEC with 19097 genes | Section 4 | Figures 4 and S5-S7 |
| | Time_course | SMARTer/ Fluidigm | plate-based | short reads | GEO: GSE75748 | 758 cells × 19189 genes; 6 time points | Section 5 | Figures 5 and S8 |
| Subsampled datasets | mean.top | SMARTer/ Fluidigm | plate-based | short reads | Subsampled from dataset Human_ESC | 350 cells × 1024 genes; 2 cell types: H1 ESC and DEC | Section 6 | Figures 6 and S9 |
| | mean.low | SMARTer/ Fluidigm | plate-based | short reads | Subsampled from dataset Human_ESC | 350 cells × 1024 genes; 2 cell types: H1 ESC and DEC | | |
| | sd.top | SMARTer/ Fluidigm | plate-based | short reads | Subsampled from dataset Human_ESC | 350 cells × 1024 genes; 2 cell types: H1 ESC and DEC | | |
| | sd.low | SMARTer/ Fluidigm | plate-based | short reads | Subsampled from dataset Human_ESC | 350 cells × 1024 genes; 2 cell types: H1 ESC and DEC | | |
| | global.random | SMARTer/ Fluidigm | plate-based | short reads | Subsampled from dataset Human_ESC | 350 cells × 1024 genes; 2 cell types: H1 ESC and DEC | | |
| Cross-platform datasets | sc_10X | 10X Genomics/ Chromium | droplet-based | UMI | https://github.com/LuyiTian/sc_mixology/tree/master/data/csv | 902 cells × 16468 genes; 3 cell types:H1975, H2228, HCC827 | Section 7 | Figure 7A, 7B, S10, and S11; Tables S8 |
| | sc_CEL-seq2 | CEL-seq2/ Fluidigm | plate-based | UMI | https://github.com/LuyiTian/sc_mixology/tree/master/data/csv | 274 cells × 28204 genes; 3 cell types:H1975, H2228, HCC827 | | |
| | sc_Drop-seq | Drop-seq/ Droplet | droplet-based | UMI | https://github.com/LuyiTian/sc_mixology/tree/master/data/csv | 225 cells × 15127 genes; 3 cell types:H1975, H2228, HCC827 | | |
| Scalability datasets | pmbc_1k | 10X Genomics/ Chromium | droplet-based | UMI | https://support.10xgenomics.com/single-cell-gene-expression/datasets/3.0.0/pbmc_10k_v3 | 1000 cells × 10000 genes; 10 clusters | | Figures 7C, 7D, S12; Tables S9 |
| | pmbc_5k | | | | | 5000 cells × 10000 genes; 10 clusters | | |
| | pmbc_25k | | | | | 25000 cells × 10000 genes; 10 clusters | | |
| | pmbc_50k | | | | | 50000 cells × 10000 genes; 10 clusters | | |
| | pmbc_100k | | | | | 100000 cells × 10000 genes; 10 clusters | | |

**Table S2. Multiple metrics are used to measure the clustering output for each of the compared methods on CIDR simulated data.** ARI, adjusted rand index; NMI, normalized mutual information; F score, the harmonic mean of precision and recall; AUC, area under the ROC curve; ACC, accuracy; RI, rand index; MI, mutual information; VI, variation of information; NVI, normalized variation of information; ID, information distance; NID, normalized information distance. Underline-highlighted metrics are used to plot **Figure S2A**.

| | **ARI** | **NMI** | **F score** | **AUC** | **ACC** | **RI** | **MI** | **VI** | **NVI** | **ID** | **NID** |
|---|---|---|---|---|---|---|---|---|---|---|---|
| Dropout | 0.637 | 0.570 | 0.761 | 0.821 | 0.841 | 0.840 | 0.626 | 0.944 | 0.601 | 0.473 | 0.430 |
| ENHANCE | -0.001 | 0.015 | 0.414 | 0.504 | 0.494 | 0.490 | 0.017 | 1.842 | 0.991 | 1.082 | 0.985 |
| scGAIN | 0.005 | 0.014 | 0.363 | 0.509 | 0.551 | 0.548 | 0.015 | 2.105 | 0.993 | 1.084 | 0.986 |
| DCA | 0.014 | 0.026 | 0.352 | 0.514 | 0.567 | 0.564 | 0.028 | 2.140 | 0.987 | 1.070 | 0.974 |
| AutoImpute | 0.091 | 0.122 | 0.409 | 0.552 | 0.597 | 0.594 | 0.134 | 1.903 | 0.934 | 0.964 | 0.878 |
| SCRABBLE | 0.024 | 0.039 | 0.362 | 0.519 | 0.570 | 0.567 | 0.043 | 2.098 | 0.980 | 1.055 | 0.961 |
| VIPER | 0.059 | 0.062 | 0.390 | 0.536 | 0.581 | 0.578 | 0.068 | 2.032 | 0.968 | 1.030 | 0.938 |
| DeepImpute | 0.132 | 0.139 | 0.435 | 0.572 | 0.615 | 0.613 | 0.152 | 1.873 | 0.925 | 0.946 | 0.861 |
| MAGIC | 0.368 | 0.402 | 0.628 | 0.713 | 0.682 | 0.680 | 0.442 | 1.003 | 0.694 | 0.657 | 0.598 |
| DrImpute | 0.450 | 0.431 | 0.657 | 0.742 | 0.743 | 0.741 | 0.474 | 1.110 | 0.701 | 0.625 | 0.569 |
| SAVER | 0.845 | 0.799 | 0.898 | 0.924 | 0.932 | 0.931 | 0.878 | 0.441 | 0.334 | 0.221 | 0.201 |
| scIGANs(w/o) | 0.941 | 0.911 | 0.961 | 0.971 | 0.974 | 0.974 | 1.001 | 0.196 | 0.164 | 0.098 | 0.089 |
| scImpute | 1.000 | 1.000 | 1.000 | 1.000 | 1.000 | 1.000 | 1.099 | 0.000 | 0.000 | 0.000 | 0.000 |
| scIGANs(w/) | 1.000 | 1.000 | 1.000 | 1.000 | 1.000 | 1.000 | 1.099 | 0.000 | 0.000 | 0.000 | 0.000 |
| Full | 1.000 | 1.000 | 1.000 | 1.000 | 1.000 | 1.000 | 1.099 | 0.000 | 0.000 | 0.000 | 0.000 |

**Table S3**. **Multiple metrics are used to measure the clustering outputs for each of the compared methods on the Splatter simulated dataset with three different dropout rates.** ARI, adjusted rand index; NMI, normalized mutual information; F score, the harmonic mean of precision and recall; AUC, area under the ROC curve; ACC, accuracy; RI, rand index; MI, mutual information; VI, variation of information; NVI, normalized variation of information; ID, information distance; NID, normalized information distance. Underline-highlighted ARIs are used to plot **Figure S2E**.

| | | RI | ARI | MI | NMI | VI | NVI | ID | NID | F score | AUC | ACC |
|---|---|---|---|---|---|---|---|---|---|---|---|---|
| Dropout rate (71.0%) | Full | 0.998 | 0.995 | 1.027 | 0.987 | 0.026 | 0.025 | 0.014 | 0.013 | 0.997 | 0.997 | 0.998 |
| | Dropout | 0.530 | 0.007 | 0.004 | 0.004 | 2.014 | 0.998 | 1.035 | 0.996 | 0.389 | 0.504 | 0.530 |
| | AutoImpute | 0.741 | 0.446 | 0.551 | 0.531 | 0.972 | 0.638 | 0.488 | 0.469 | 0.654 | 0.724 | 0.741 |
| | DCA | 0.544 | 0.003 | 0.006 | 0.005 | 2.121 | 0.997 | 1.088 | 0.995 | 0.356 | 0.502 | 0.545 |
| | DeepImpute | 0.526 | 0.004 | 0.002 | 0.001 | 2.014 | 0.999 | 1.038 | 0.999 | 0.391 | 0.503 | 0.527 |
| | DrImpute | 0.781 | 0.551 | 0.451 | 0.434 | 0.985 | 0.686 | 0.589 | 0.566 | 0.736 | 0.789 | 0.781 |
| | ENHANCE | 0.416 | -0.016 | 0.008 | 0.008 | 1.435 | 0.995 | 1.032 | 0.992 | 0.496 | 0.491 | 0.416 |
| | MAGIC | 0.829 | 0.653 | 0.598 | 0.575 | 0.654 | 0.522 | 0.441 | 0.425 | 0.797 | 0.845 | 0.829 |
| | SAVER | 0.543 | 0.013 | 0.025 | 0.023 | 2.055 | 0.988 | 1.040 | 0.977 | 0.373 | 0.507 | 0.544 |
| | scGAIN | 0.475 | 0.007 | 0.003 | 0.003 | 1.639 | 0.998 | 1.036 | 0.997 | 0.465 | 0.504 | 0.475 |
| | scImpute | 0.502 | -0.001 | 0.000 | 0.000 | 1.773 | 1.000 | 1.039 | 1.000 | 0.423 | 0.500 | 0.502 |
| | SCRABBLE | 0.946 | 0.884 | 0.870 | 0.824 | 0.355 | 0.290 | 0.186 | 0.176 | 0.927 | 0.940 | 0.946 |
| | VIPER | 0.545 | 0.015 | 0.009 | 0.009 | 2.090 | 0.996 | 1.060 | 0.991 | 0.373 | 0.508 | 0.546 |
| | scIGANs(w/o) | 0.812 | 0.599 | 0.542 | 0.521 | 0.995 | 0.648 | 0.498 | 0.479 | 0.749 | 0.800 | 0.813 |
| | scIGANs(w/) | 1.000 | 1.000 | 1.039 | 1.000 | 0.000 | 0.000 | 0.000 | 0.000 | 1.000 | 1.000 | 1.000 |
| Dropout rate (83.0%) | Full | 0.998 | 0.995 | 1.027 | 0.987 | 0.026 | 0.025 | 0.014 | 0.013 | 0.997 | 0.997 | 0.998 |
| | Dropout | 0.527 | -0.004 | 0.001 | 0.001 | 2.054 | 1.000 | 1.038 | 0.999 | 0.379 | 0.499 | 0.527 |
| | AutoImpute | 0.711 | 0.378 | 0.447 | 0.425 | 1.198 | 0.728 | 0.606 | 0.575 | 0.608 | 0.689 | 0.711 |
| | DCA | 0.528 | 0.001 | 0.002 | 0.002 | 2.030 | 0.999 | 1.037 | 0.998 | 0.384 | 0.501 | 0.528 |
| | DeepImpute | 0.537 | 0.008 | 0.006 | 0.005 | 2.071 | 0.997 | 1.037 | 0.995 | 0.378 | 0.505 | 0.538 |
| | DrImpute | 0.500 | -0.008 | 0.002 | 0.002 | 1.889 | 0.999 | 1.037 | 0.998 | 0.417 | 0.497 | 0.500 |
| | ENHANCE | 0.490 | 0.002 | 0.001 | 0.001 | 1.808 | 1.000 | 1.039 | 0.999 | 0.442 | 0.502 | 0.491 |
| | MAGIC | 0.811 | 0.609 | 0.539 | 0.518 | 0.871 | 0.618 | 0.501 | 0.482 | 0.765 | 0.815 | 0.811 |
| | SAVER | 0.515 | 0.008 | 0.005 | 0.005 | 1.895 | 0.997 | 1.034 | 0.995 | 0.415 | 0.505 | 0.515 |
| | scGAIN | 0.419 | -0.008 | 0.002 | 0.002 | 1.405 | 0.998 | 1.037 | 0.998 | 0.502 | 0.496 | 0.419 |
| | scImpute | 0.518 | 0.033 | 0.022 | 0.021 | 1.728 | 0.987 | 1.017 | 0.979 | 0.443 | 0.518 | 0.519 |
| | SCRABBLE | 0.940 | 0.870 | 0.857 | 0.809 | 0.385 | 0.310 | 0.202 | 0.191 | 0.917 | 0.932 | 0.940 |
| | VIPER | 0.544 | 0.004 | 0.005 | 0.005 | 2.121 | 0.998 | 1.087 | 0.995 | 0.357 | 0.503 | 0.545 |
| | scIGANs(w/o) | 0.507 | -0.004 | 0.007 | 0.006 | 1.844 | 0.996 | 1.033 | 0.994 | 0.412 | 0.499 | 0.507 |
| | scIGANs(w/) | 1.000 | 1.000 | 1.039 | 1.000 | 0.000 | 0.000 | 0.000 | 0.000 | 1.000 | 1.000 | 1.000 |
| Dropout rate (87.0%) | Full | 0.998 | 0.995 | 1.027 | 0.987 | 0.026 | 0.025 | 0.014 | 0.013 | 0.997 | 0.997 | 0.998 |
| | Dropout | 0.526 | -0.001 | 0.001 | 0.001 | 2.039 | 1.000 | 1.039 | 0.999 | 0.385 | 0.500 | 0.526 |
| | AutoImpute | 0.692 | 0.332 | 0.365 | 0.340 | 1.383 | 0.791 | 0.709 | 0.660 | 0.573 | 0.665 | 0.692 |
| | DCA | 0.519 | -0.005 | 0.003 | 0.003 | 2.011 | 0.998 | 1.036 | 0.997 | 0.392 | 0.498 | 0.519 |
| | DeepImpute | 0.542 | -0.001 | 0.001 | 0.001 | 2.128 | 0.999 | 1.090 | 0.999 | 0.354 | 0.501 | 0.542 |
| | DrImpute | 0.499 | -0.001 | 0.003 | 0.003 | 1.784 | 0.998 | 1.037 | 0.997 | 0.426 | 0.500 | 0.500 |
| | ENHANCE | 0.433 | 0.014 | 0.006 | 0.006 | 1.413 | 0.996 | 1.033 | 0.994 | 0.511 | 0.509 | 0.434 |
| | MAGIC | 0.758 | 0.488 | 0.424 | 0.408 | 1.176 | 0.735 | 0.615 | 0.592 | 0.685 | 0.748 | 0.758 |
| | SAVER | 0.501 | -0.007 | 0.001 | 0.001 | 1.908 | 0.999 | 1.038 | 0.999 | 0.416 | 0.497 | 0.502 |
| | scGAIN | 0.457 | 0.001 | 0.002 | 0.002 | 1.662 | 0.999 | 1.038 | 0.998 | 0.477 | 0.501 | 0.458 |
| | scImpute | 0.494 | 0.000 | 0.008 | 0.008 | 1.704 | 0.995 | 1.032 | 0.992 | 0.435 | 0.501 | 0.494 |
| | SCRABBLE | 0.950 | 0.892 | 0.861 | 0.825 | 0.361 | 0.295 | 0.182 | 0.175 | 0.932 | 0.946 | 0.950 |
| | VIPER | 0.512 | -0.002 | 0.003 | 0.003 | 1.969 | 0.999 | 1.037 | 0.997 | 0.406 | 0.499 | 0.512 |
| | scIGANs(w/o) | 0.474 | -0.003 | 0.003 | 0.003 | 1.714 | 0.998 | 1.036 | 0.997 | 0.455 | 0.499 | 0.474 |
| | scIGANs(w/) | 0.995 | 0.990 | 1.021 | 0.983 | 0.035 | 0.033 | 0.018 | 0.017 | 0.993 | 0.995 | 0.995 |

**Table S4. Multiple metrics are used to measure the clustering output for each of the compared methods on Splatter simulated datasets with three different dropout rates (100 replicates for each dropout rate).** ARI, adjusted rand index; NMI, normalized mutual information; F score, the harmonic mean of precision and recall; AUC, area under the ROC curve; ACC, accuracy; RI, rand index; MI, mutual information; VI, variation information; NVI, normalized variation information; ID, information distance; NID, normalized information distance.

**Large tables are provided in the separate XLSX file**

**Table S5. Multiple metrics are used to measure the clustering output for each of the compared methods on real scRNA-seq data from the human brain.** ARI, adjusted rand index; NMI, normalized mutual information; F score, the harmonic mean of precision and recall; AUC, area under the ROC curve; ACC, accuracy; RI, rand index; MI, mutual information; VI, variation of information; NVI, normalized variation of information; ID, information distance; NID, normalized information distance. Underline-highlighted metrics are used to plot **Figure 2D**.

| | NMI | ARI | F score | AUC | ACC | RI | MI | VI | NVI | ID | NID |
|---|---|---|---|---|---|---|---|---|---|---|---|
| Dropout | 0.582 | 0.355 | 0.487 | 0.675 | 0.798 | 0.798 | 1.059 | 1.489 | 0.584 | 0.762 | 0.418 |
| SCRABBLE | 0.126 | 0.083 | 0.367 | 0.575 | 0.481 | 0.480 | 0.225 | 2.207 | 0.908 | 1.562 | 0.874 |
| DCA | 0.496 | 0.328 | 0.472 | 0.669 | 0.781 | 0.780 | 0.886 | 1.743 | 0.663 | 0.900 | 0.504 |
| MAGIC | 0.615 | 0.390 | 0.512 | 0.689 | 0.812 | 0.812 | 1.169 | 1.349 | 0.536 | 0.731 | 0.385 |
| DeepImpute | 0.568 | 0.364 | 0.492 | 0.678 | 0.803 | 0.802 | 1.057 | 1.534 | 0.592 | 0.805 | 0.432 |
| scIGANs(w/) | 0.540 | 0.364 | 0.492 | 0.678 | 0.802 | 0.802 | 0.999 | 1.638 | 0.621 | 0.850 | 0.460 |
| AutoImpute | 0.666 | 0.559 | 0.634 | 0.745 | 0.876 | 0.876 | 1.366 | 1.104 | 0.447 | 0.684 | 0.334 |
| DrImpute | 0.642 | 0.471 | 0.580 | 0.732 | 0.834 | 0.833 | 1.189 | 1.260 | 0.515 | 0.663 | 0.358 |
| ENHANCE | 0.706 | 0.538 | 0.626 | 0.751 | 0.862 | 0.862 | 1.359 | 0.993 | 0.422 | 0.565 | 0.294 |
| SAVER | 0.602 | 0.575 | 0.671 | 0.804 | 0.856 | 0.856 | 1.076 | 1.231 | 0.534 | 0.711 | 0.398 |
| scGAIN | 0.138 | 0.092 | 0.332 | 0.562 | 0.626 | 0.625 | 0.246 | 2.655 | 0.915 | 1.540 | 0.862 |
| scImpute | 0.672 | 0.545 | 0.638 | 0.767 | 0.857 | 0.857 | 1.223 | 1.161 | 0.487 | 0.598 | 0.328 |
| VIPER | 0.544 | 0.306 | 0.460 | 0.662 | 0.768 | 0.767 | 0.972 | 1.592 | 0.621 | 0.815 | 0.456 |
| scIGANs(w/o) | 0.349 | 0.243 | 0.395 | 0.620 | 0.766 | 0.765 | 0.631 | 2.330 | 0.787 | 1.175 | 0.651 |

**Table S6. Multiple metrics are used to measure the clustering output for each of the compared methods on real scRNA-seq data for ERCC spike-in RNAs.** ARI, adjusted rand index; NMI, normalized mutual information; F score, the harmonic mean of precision and recall; AUC, area under the ROC curve; ACC, accuracy; RI, rand index; MI, mutual information; VI, variation of information; NVI, normalized variation of information; ID, information distance; NID, normalized information distance.

| | **ARI** | **NMI** | **F score** | **AUC** | **ACC** | **RI** | **MI** | **VI** | **NVI** | **ID** | **NID** |
|---|---|---|---|---|---|---|---|---|---|---|---|
| Dropout | 0.038 | 0.050 | 0.430 | 0.524 | 0.516 | 0.514 | 0.055 | 1.751 | 0.969 | 1.043 | 0.950 |
| AutoImpute | 0.145 | 0.183 | 0.453 | 0.578 | 0.608 | 0.607 | 0.201 | 1.717 | 0.895 | 0.897 | 0.817 |
| DCA | 0.065 | 0.072 | 0.434 | 0.538 | 0.542 | 0.541 | 0.079 | 1.834 | 0.959 | 1.019 | 0.928 |
| DeepImpute | 0.098 | 0.094 | 0.454 | 0.557 | 0.559 | 0.558 | 0.104 | 1.699 | 0.943 | 0.995 | 0.906 |
| DrImpute | 0.194 | 0.181 | 0.513 | 0.610 | 0.604 | 0.603 | 0.199 | 1.501 | 0.883 | 0.900 | 0.819 |
| ENHANCE | 0.429 | 0.387 | 0.624 | 0.718 | 0.746 | 0.745 | 0.425 | 1.333 | 0.758 | 0.674 | 0.613 |
| MAGIC | 0.429 | 0.387 | 0.423 | 0.542 | 0.563 | 0.745 | 0.425 | 1.333 | 0.758 | 0.674 | 0.613 |
| SAVER | 0.041 | 0.052 | 0.425 | 0.526 | 0.525 | 0.524 | 0.057 | 1.771 | 0.969 | 1.041 | 0.948 |
| scGAIN | 0.001 | 0.018 | 0.491 | 0.501 | 0.359 | 0.357 | 0.019 | 1.247 | 0.985 | 1.079 | 0.982 |
| scImpute | 0.416 | 0.399 | 0.649 | 0.734 | 0.711 | 0.710 | 0.439 | 1.010 | 0.697 | 0.660 | 0.601 |
| SCRABBLE | 0.035 | 0.033 | 0.404 | 0.522 | 0.540 | 0.538 | 0.037 | 1.937 | 0.981 | 1.062 | 0.967 |
| VIPER | 0.008 | 0.035 | 0.404 | 0.507 | 0.511 | 0.509 | 0.038 | 1.809 | 0.979 | 1.060 | 0.965 |
| scIGANs(w/o) | 0.034 | 0.048 | 0.416 | 0.521 | 0.527 | 0.525 | 0.053 | 1.798 | 0.971 | 1.046 | 0.952 |
| scIGANs(w/) | 0.042 | 0.052 | 0.405 | 0.525 | 0.546 | 0.544 | 0.057 | 1.916 | 0.971 | 1.041 | 0.948 |

**Table S7. Multiple metrics are used to measure the clustering output for each of the compared methods on real scRNA-seq data for cell-cycle states of mESCs.** ARI, adjusted rand index; NMI, normalized mutual information; F score, the harmonic mean of precision and recall; AUC, area under the ROC curve; ACC, accuracy; RI, rand index; MI, mutual information; VI, variation of information; NVI, normalized variation of information; ID, information distance; NID, normalized information distance. The metrics are used to plot **Figure S4A**.

| | **ARI** | **NMI** | **F score** | **AUC** | **ACC** | **RI** | **MI** | **VI** | **NVI** | **ID** | **NID** |
|---|---|---|---|---|---|---|---|---|---|---|---|
| Dropout | 0.025 | 0.038 | 0.449 | 0.517 | 0.476 | 0.474 | 0.042 | 1.640 | 0.975 | 1.057 | 0.962 |
| AutoImpute | 0.097 | 0.109 | 0.469 | 0.558 | 0.541 | 0.540 | 0.120 | 1.598 | 0.930 | 0.978 | 0.891 |
| DCA | 0.128 | 0.106 | 0.471 | 0.573 | 0.574 | 0.573 | 0.116 | 1.734 | 0.937 | 0.983 | 0.894 |
| DeepImpute | 0.002 | 0.024 | 0.492 | 0.502 | 0.360 | 0.357 | 0.027 | 1.225 | 0.979 | 1.072 | 0.976 |
| DrImpute | 0.033 | 0.091 | 0.486 | 0.523 | 0.426 | 0.424 | 0.100 | 1.304 | 0.929 | 0.998 | 0.909 |
| ENHANCE | 0.006 | 0.048 | 0.483 | 0.505 | 0.386 | 0.383 | 0.053 | 1.313 | 0.961 | 1.046 | 0.952 |
| MAGIC | 0.006 | 0.048 | 0.400 | 0.538 | 0.576 | 0.383 | 0.053 | 1.313 | 0.961 | 1.046 | 0.952 |
| SAVER | 0.099 | 0.106 | 0.472 | 0.559 | 0.541 | 0.539 | 0.116 | 1.619 | 0.933 | 0.983 | 0.894 |
| scGAIN | -0.003 | 0.006 | 0.404 | 0.501 | 0.498 | 0.496 | 0.006 | 1.812 | 0.997 | 1.092 | 0.994 |
| scImpute | 0.561 | 0.590 | 0.740 | 0.820 | 0.779 | 0.778 | 0.648 | 0.566 | 0.466 | 0.451 | 0.410 |
| SCRABBLE | 0.523 | 0.534 | 0.716 | 0.796 | 0.762 | 0.761 | 0.587 | 0.717 | 0.550 | 0.512 | 0.466 |
| VIPER | 0.003 | 0.030 | 0.493 | 0.502 | 0.358 | 0.356 | 0.032 | 1.206 | 0.974 | 1.066 | 0.970 |
| scIGANs(w/o) | 0.110 | 0.159 | 0.448 | 0.562 | 0.577 | 0.576 | 0.174 | 1.710 | 0.908 | 0.924 | 0.841 |
| scIGANs(w/) | 1.000 | 1.000 | 1.000 | 1.000 | 1.000 | 1.000 | 1.099 | 0.000 | 0.000 | 0.000 | 0.000 |

**Table S8. Multiple metrics are used to measure the clustering output for each of the compared methods on scRNA-seq datasets from three different protocols.** ARI, adjusted rand index; NMI, normalized mutual information; F score, the harmonic mean of precision and recall; AUC, area under the ROC curve; ACC, accuracy; RI, rand index; MI, mutual information; VI, variation of information; NVI, normalized variation of information; ID, information distance; NID, normalized information distance. Underline-highlighted columns are used to plot **Figure 7B and S11**.

| Dataset | Method | RI | ARI | MI | NMI | VI | NVI | ID | NID | F score | AUC | ACC |
|---|---|---|---|---|---|---|---|---|---|---|---|---|
| **10X Genomics (zero rate 45.0%)** | Dropout | 0.996 | 0.990 | 1.076 | 0.981 | 0.042 | 0.037 | 0.021 | 0.019 | 0.993 | 0.995 | 0.996 |
| | AutoImpute | 0.979 | 0.953 | 1.014 | 0.925 | 0.165 | 0.140 | 0.083 | 0.075 | 0.969 | 0.977 | 0.979 |
| | DCA | 0.997 | 0.994 | 1.083 | 0.988 | 0.027 | 0.024 | 0.013 | 0.012 | 0.996 | 0.997 | 0.997 |
| | DeepImpute | 0.997 | 0.993 | 1.082 | 0.986 | 0.030 | 0.027 | 0.015 | 0.014 | 0.996 | 0.997 | 0.997 |
| | DrImpute | 0.996 | 0.990 | 1.076 | 0.981 | 0.042 | 0.037 | 0.021 | 0.019 | 0.993 | 0.995 | 0.996 |
| | ENHANCE | 0.791 | 0.560 | 0.704 | 0.642 | 0.643 | 0.477 | 0.393 | 0.358 | 0.726 | 0.800 | 0.791 |
| | MAGIC | 0.953 | 0.894 | 0.945 | 0.862 | 0.297 | 0.239 | 0.152 | 0.138 | 0.930 | 0.948 | 0.953 |
| | SAVER | 0.996 | 0.990 | 1.076 | 0.981 | 0.042 | 0.037 | 0.021 | 0.019 | 0.993 | 0.995 | 0.996 |
| | scGAIN | 0.880 | 0.733 | 0.848 | 0.774 | 0.470 | 0.356 | 0.248 | 0.226 | 0.825 | 0.872 | 0.880 |
| | scImpute | 1.000 | 1.000 | 1.097 | 1.000 | 0.000 | 0.000 | 0.000 | 0.000 | 1.000 | 1.000 | 1.000 |
| | SCRABBLE | 0.990 | 0.977 | 1.055 | 0.961 | 0.084 | 0.074 | 0.043 | 0.039 | 0.985 | 0.988 | 0.990 |
| | VIPER | 0.999 | 0.997 | 1.089 | 0.993 | 0.015 | 0.013 | 0.007 | 0.007 | 0.998 | 0.998 | 0.999 |
| | scIGANs | 1.000 | 1.000 | 1.097 | 1.000 | 0.000 | 0.000 | 0.000 | 0.000 | 1.000 | 1.000 | 1.000 |
| **CEL-seq2 (zero rate 74.7%)** | Dropout | 0.990 | 0.977 | 1.049 | 0.966 | 0.071 | 0.063 | 0.037 | 0.034 | 0.985 | 0.988 | 0.990 |
| | AutoImpute | 0.575 | 0.116 | 0.259 | 0.239 | 1.508 | 0.854 | 0.825 | 0.761 | 0.461 | 0.565 | 0.577 |
| | DCA | 0.995 | 0.989 | 1.064 | 0.981 | 0.041 | 0.037 | 0.021 | 0.019 | 0.993 | 0.994 | 0.995 |
| | DeepImpute | 0.990 | 0.977 | 1.049 | 0.966 | 0.071 | 0.063 | 0.037 | 0.034 | 0.985 | 0.988 | 0.990 |
| | DrImpute | 0.980 | 0.955 | 1.008 | 0.930 | 0.152 | 0.131 | 0.076 | 0.070 | 0.970 | 0.977 | 0.980 |
| | ENHANCE | 1.000 | 1.000 | 1.084 | 1.000 | 0.000 | 0.000 | 0.000 | 0.000 | 1.000 | 1.000 | 1.000 |
| | MAGIC | 0.545 | 0.007 | 0.019 | 0.017 | 2.090 | 0.991 | 1.065 | 0.983 | 0.366 | 0.507 | 0.546 |
| | SAVER | 0.990 | 0.977 | 1.049 | 0.966 | 0.071 | 0.063 | 0.037 | 0.034 | 0.985 | 0.988 | 0.990 |
| | scGAIN | 0.498 | 0.011 | 0.019 | 0.017 | 1.868 | 0.990 | 1.065 | 0.983 | 0.424 | 0.508 | 0.500 |
| | scImpute | 1.000 | 1.000 | 1.084 | 1.000 | 0.000 | 0.000 | 0.000 | 0.000 | 1.000 | 1.000 | 1.000 |
| | SCRABBLE | 0.500 | -0.005 | 0.005 | 0.005 | 1.812 | 0.997 | 1.078 | 0.995 | 0.406 | 0.500 | 0.501 |
| | VIPER | 0.563 | 0.026 | 0.034 | 0.031 | 2.100 | 0.984 | 1.050 | 0.969 | 0.365 | 0.517 | 0.564 |
| | scIGANs | 1.000 | 1.000 | 1.084 | 1.000 | 0.000 | 0.000 | 0.000 | 0.000 | 1.000 | 1.000 | 1.000 |
| **Drop-seq (zero rate 62.1%)** | Dropout | 0.987 | 0.972 | 1.046 | 0.961 | 0.083 | 0.074 | 0.043 | 0.039 | 0.982 | 0.986 | 0.988 |
| | AutoImpute | 0.821 | 0.599 | 0.686 | 0.629 | 0.806 | 0.540 | 0.405 | 0.371 | 0.737 | 0.800 | 0.821 |
| | DCA | 0.935 | 0.854 | 0.944 | 0.863 | 0.291 | 0.236 | 0.150 | 0.137 | 0.905 | 0.926 | 0.935 |
| | DeepImpute | 0.994 | 0.986 | 1.063 | 0.977 | 0.048 | 0.043 | 0.025 | 0.023 | 0.991 | 0.993 | 0.994 |
| | DrImpute | 0.908 | 0.794 | 0.888 | 0.813 | 0.403 | 0.312 | 0.205 | 0.187 | 0.865 | 0.897 | 0.909 |
| | ENHANCE | 0.994 | 0.986 | 1.063 | 0.977 | 0.048 | 0.043 | 0.025 | 0.023 | 0.991 | 0.993 | 0.994 |
| | MAGIC | 0.544 | 0.058 | 0.140 | 0.129 | 1.737 | 0.925 | 0.946 | 0.871 | 0.429 | 0.535 | 0.546 |
| | SAVER | 0.994 | 0.986 | 1.063 | 0.977 | 0.048 | 0.043 | 0.025 | 0.023 | 0.991 | 0.993 | 0.994 |
| | scGAIN | 0.478 | -0.005 | 0.018 | 0.017 | 1.801 | 0.990 | 1.068 | 0.983 | 0.425 | 0.500 | 0.481 |
| | scImpute | 1.000 | 1.000 | 1.086 | 1.000 | 0.000 | 0.000 | 0.000 | 0.000 | 1.000 | 1.000 | 1.000 |
| | SCRABBLE | 0.517 | 0.014 | 0.036 | 0.033 | 1.823 | 0.981 | 1.050 | 0.967 | 0.409 | 0.511 | 0.519 |
| | VIPER | 0.923 | 0.829 | 0.874 | 0.805 | 0.401 | 0.314 | 0.212 | 0.195 | 0.890 | 0.920 | 0.923 |
| | scIGANs | 1.000 | 1.000 | 1.086 | 1.000 | 0.000 | 0.000 | 0.000 | 0.000 | 1.000 | 1.000 | 1.000 |

**Table S9**. **Running time and memory usage of imputation methods on different data sizes.** These values are used to plot Figures 7D and S12B, with the exception that running time >48 hours (Failed_T) or the memory usage > 64 GB (Failed_M) were replaced by auxiliary values of 49 and 65, so that all methods will be plotted in the figures at all data size points.

| | **Cell numbers:** | **1k** | **5k** | **25k** | **50k** | **100k** |
|---|---|---|---|---|---|---|
| **Elapsed time (hour)** | **AutoImpute** | 1.13 | 2.92 | 20.86 | ***Failed_T*** | ***Failed_T*** |
| | **DCA** | 0.10 | 0.48 | 2.87 | 4.77 | ***Failed_T*** |
| | **DeepImpute** | 0.02 | 0.05 | 0.20 | 0.43 | 1.31 |
| | **DrImpute** | 0.04 | 0.51 | 3.01 | ***Failed_M*** | ***Failed_M*** |
| | **ENHANCE** | 0.01 | 0.06 | 0.93 | ***Failed_M*** | ***Failed_M*** |
| | **MAGIC** | 0.01 | 0.03 | 0.17 | 0.42 | 1.05 |
| | **SAVER** | 0.34 | 0.29 | ***Failed_M*** | ***Failed_M*** | ***Failed_M*** |
| | **scGAIN** | 0.05 | 1.07 | 5.18 | 10.42 | ***Failed_M*** |
| | **scIGANs** | 3.40 | 5.61 | 10.07 | 15.53 | 29.79 |
| | **scIGANs_GPU** | 0.07 | 0.22 | 0.92 | 2.09 | 6.03 |
| | **scImpute** | 0.10 | 1.26 | 12.35 | ***Failed_M*** | ***Failed_M*** |
| | **SCRABBLE** | ***Failed_T*** | ***Failed_T*** | ***Failed_T*** | ***Failed_T*** | ***Failed_T*** |
| | **VIPER** | 0.80 | 14.68 | ***Failed_T*** | ***Failed_M*** | ***Failed_M*** |
| **Used memory (GB)** | **AutoImpute** | 12.66 | 13.74 | 22.17 | ***Failed_T*** | ***Failed_T*** |
| | **DCA** | 12.63 | 18.52 | 40.07 | 60.45 | ***Failed_T*** |
| | **DeepImpute** | 1.77 | 12.65 | 18.49 | 23.48 | 61.92 |
| | **DrImpute** | 1.21 | 4.95 | 18.97 | ***Failed_M*** | ***Failed_M*** |
| | **ENHANCE** | 1.30 | 4.22 | 28.48 | ***Failed_M*** | ***Failed_M*** |
| | **MAGIC** | 0.25 | 2.23 | 10.60 | 22.10 | 49.51 |
| | **SAVER** | 0.85 | 3.57 | ***Failed_M*** | ***Failed_M*** | ***Failed_M*** |
| | **scGAIN** | 11.63 | 16.52 | 43.07 | 61.45 | ***Failed_M*** |
| | **scIGANs** | 4.08 | 5.44 | 22.30 | 42.04 | 47.78 |
| | **scIGANs_GPU** | 2.50 | 3.10 | 7.90 | 16.50 | 42.70 |
| | **scImpute** | 1.14 | 3.74 | 20.19 | ***Failed_M*** | ***Failed_M*** |
| | **SCRABBLE** | ***Failed_T*** | ***Failed_T*** | ***Failed_T*** | ***Failed_T*** | ***Failed_T*** |
| | **VIPER** | 1.60 | 6.76 | ***Failed_T*** | ***Failed_M*** | ***Failed_M*** |
